# From pose to behavior: SABER integrates identity-resolved multi-animal pose tracking with large-language-model-based behavioral factor discovery

**DOI:** 10.64898/2026.09.02.749007

**Authors:** Jiaqi Zheng, Chen Peng, Shi-Yuan Zhang, Jie Wang, Wen-Ning Zhou, Hong-Mei Xue, Tian-Qi Cai, Yi-Wen Li, Xiao-Ya Su, Zhiwei Jiang, Yangyi Tan, Xiang-Dong Sun

## Abstract

Quantifying social behavior requires accurately assigning posture and actions to individual animals, but close contact, occlusion, and identity switches often hinder this. Meanwhile, current behavioral recognition pipelines still lack stable predictive accuracy, especially when using a unified multi-class model. Here we developed SABER, a locally deployable framework that couples multi-animal pose estimation with identity-preserving tracking, interpretable behavioral factor mining, and multiscale temporal behavior prediction from a single overhead video stream. Across spontaneous social interaction, mating, aggression, and four-mouse recordings, SABER improved pose-estimation accuracy and tracking continuity relative to comparator pipelines. Its behavioral factor-mining procedure identified interpretable kinematic, postural, and social descriptors, and temporal integration refined behavioral-category classification. SABER offers an intuitive, open-source interface to facilitate use. Applied to social-defeat-stress mice, SABER detected reduced approach behavior and a multivariate behavioral profile that distinguished depression-susceptible from control animals. SABER outputs could also be synchronized with miniscope calcium recordings, enabling joint analysis of behavioral states and neuronal population activity. SABER therefore provides an accessible, identity-resolved route from single-view social-interaction video to behavioral phenotyping and brain–behavior analysis.

## Introduction

Social behavior is a fundamental feature of animal life and provides critical insights into how individuals communicate, cooperate, compete, and adapt to their environment^1, 2^. Social interactions are inherently dynamic processes in which each individual simultaneously generates and responds to behavioral cues from others^3^.

These reciprocal processes are central to understanding how neural circuits generate social decisions and how disruptions in social behavior contribute to neuropsychiatric disorders^4^. Therefore, comprehensive analysis of social behavior requires quantitative measurements that capture not only where animals move but also how individual body configurations, identities, and behavioral states evolve over time^5, 6^.

Recent computational approaches have greatly improved the ability to track animal pose and quantify their behavior from video recordings. Markerless pose estimation methods have replaced invasive tracking strategies and enabled detailed measurement of body configurations in freely moving animals^7–12^. However, current approaches remain insufficient for analyzing complex social interactions, in which multiple individuals frequently overlap, make contact, and rapidly change their postures^13, 14^. One major limitation is that pose estimation and identity assignment become inseparable challenges in multi-animal environments. Although existing methods perform well when analyzing individual animals, close interactions such as fighting and mating often introduce severe occlusion and ambiguous visual cues, leading to keypoint displacement, identity switches, and fragmented trajectories^14, 15^. These errors are particularly problematic because behavioral interpretation depends not only on detecting movement but also on correctly assigning actions to specific individuals. Three-dimensional systems offer a partial solution by combining multiview acquisition with synthetic occlusion augmentation to improve pose reconstruction, identification, and behavioral embedding during close social interactions^16^. However, these multiview frameworks require substantial hardware and remain susceptible to error propagation across sequential processing stages^17, 18^. A second challenge is accurately transforming pose trajectories into biologically meaningful behavioral representations. While machine-learning approaches have increasingly been used to map pose trajectories onto behavioral categories, existing strategies often rely on manually designed kinematic and postural features or unsupervised clustering approaches^3, 8, 9, 12, 16^. However, handcrafted features require substantial domain expertise, limiting the range of higher-order combinations that can be systematically explored and potentially compromising behavioral prediction accuracy and model generalization^19, 20^. While unsupervised approaches offer complementary opportunities to discover unexpected behaviors, they generally yield clusters or modules rather than direct semantic predictions, and may identify statistically distinct movement patterns that lack clear biological interpretation^12, 21, 22^. Thus, a computational framework that integrates reliable individual tracking with automated discovery of interpretable behavioral representations remains needed.

To address these challenges, we developed SABER (Spatio-temporal Action Behavior Recognition), a framework designed to quantify complex social behaviors at the individual-animal level. SABER integrates three complementary components, including an accurate multi-animal pose estimation and identity-preserving tracking module, a large language model-based behavioral factor mining framework, and a multiscale temporal behavior prediction model. We evaluated SABER across diverse mouse social interaction paradigms, including freely interacting pairs, aggressive encounters, mating behaviors, and group social interactions. By combining individual-resolved pose dynamics with automated behavioral representation discovery and temporal modeling, SABER enables scalable characterization of complex social behaviors and provides a quantitative framework for studying behavioral organization in health and disease models.

## Results

### SABER integrates pose estimation, identity tracking, factor mining, and temporal behavior prediction

We developed SABER as an integrated behavioral analysis framework consisting of three sequential components: multi-animal pose estimation using the Token Mixed Pose (TMP) model and identity-preserving trajectory reconstruction using multi-object tracking (Fig. 1a, b), behavioral feature generation from pose dynamics using large-language-model (LLM)-based behavioral factor mining (Fig. 1c), and multiscale temporal behavior prediction (Fig. 1d).

**Fig 1.**
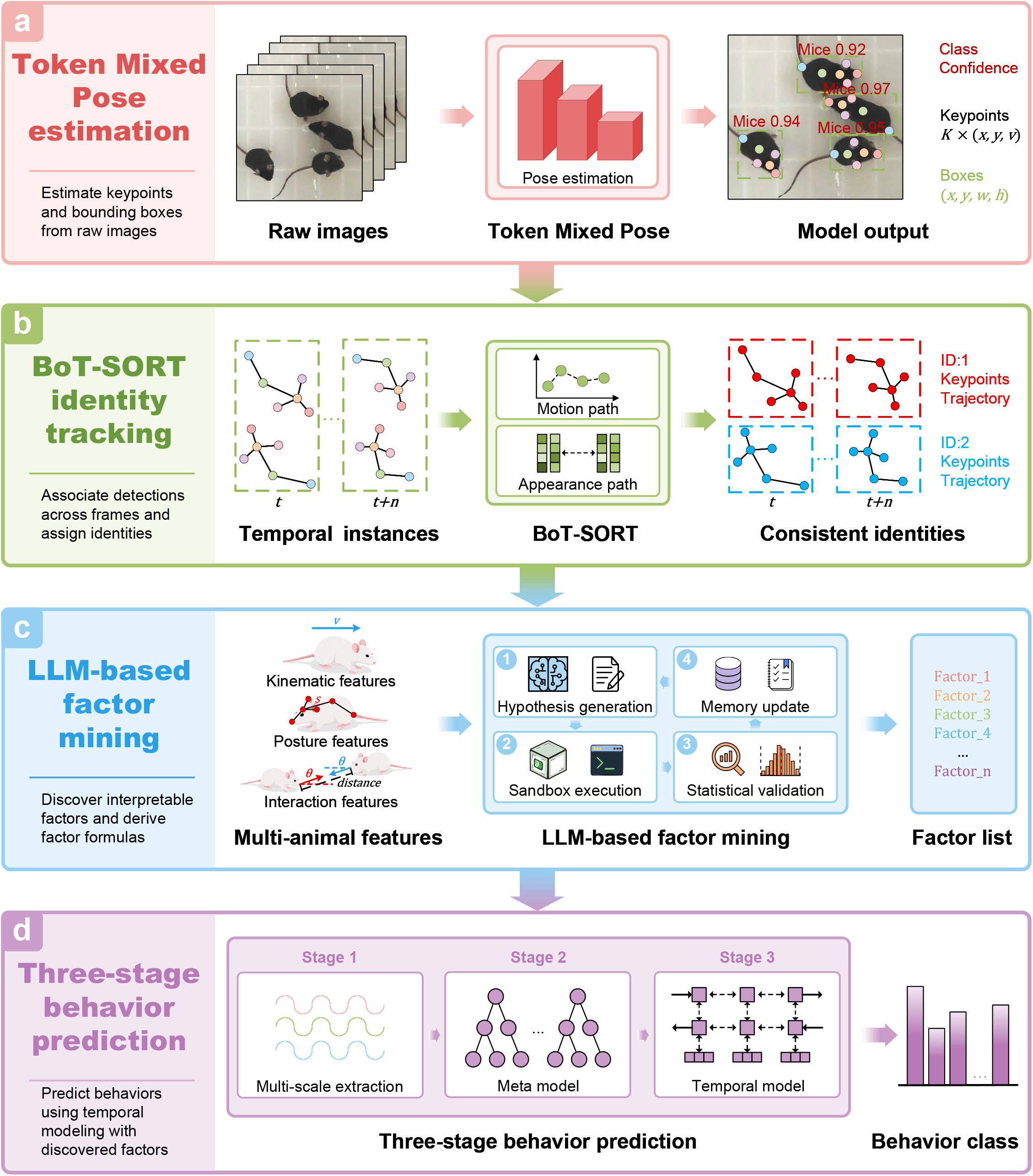
Overview of the workflow of SABER. **a**, Token Mixed Pose estimation. Overhead videos of multi-animal interactions are acquired. The Token Mixed Pose (TMP) model detects six anatomical landmarks in each animal. **b**, BoT-SORT identity tracking. BoT-SORT maintains multi-animal identities across frames. Identity-linked keypoints are converted into kinematic, within-animal postural, and interaction features. **c**, A large-language-model-guided factor-mining procedure proposes executable behavioral descriptors, evaluates them against annotated behaviors, and retains validated factors for subsequent rounds. **d**, A three-stage temporal prediction model integrates short-, intermediate-, and long-range factors to produce frame-resolved behavior probabilities and labels for each animal.

We generated behavioral datasets from freely interacting mice, including spontaneous two-mouse social interactions, aggressive encounters, mating behaviors, and four-animal group interactions, viewed from a top-down perspective. Unlike conventional single-animal behavioral assays, these paradigms involve frequent physical contact and transient occlusion, posing challenging conditions for automated behavioral analysis.

Reliable behavioral quantification first requires accurate reconstruction of individual body configurations. We therefore developed TMP, a pose estimation model optimized for multi-animal social interactions (Fig. 1a). TMP follows a one-stage supervised-learning design. The convolutional backbone extracts hierarchical visual features, the neck integrates features across scales, and the prediction head outputs keypoint locations, animal bounding boxes, and class predictions (Fig. 2a). We optimized the model end-to-end using a joint loss across these prediction tasks.

**Fig 2.**
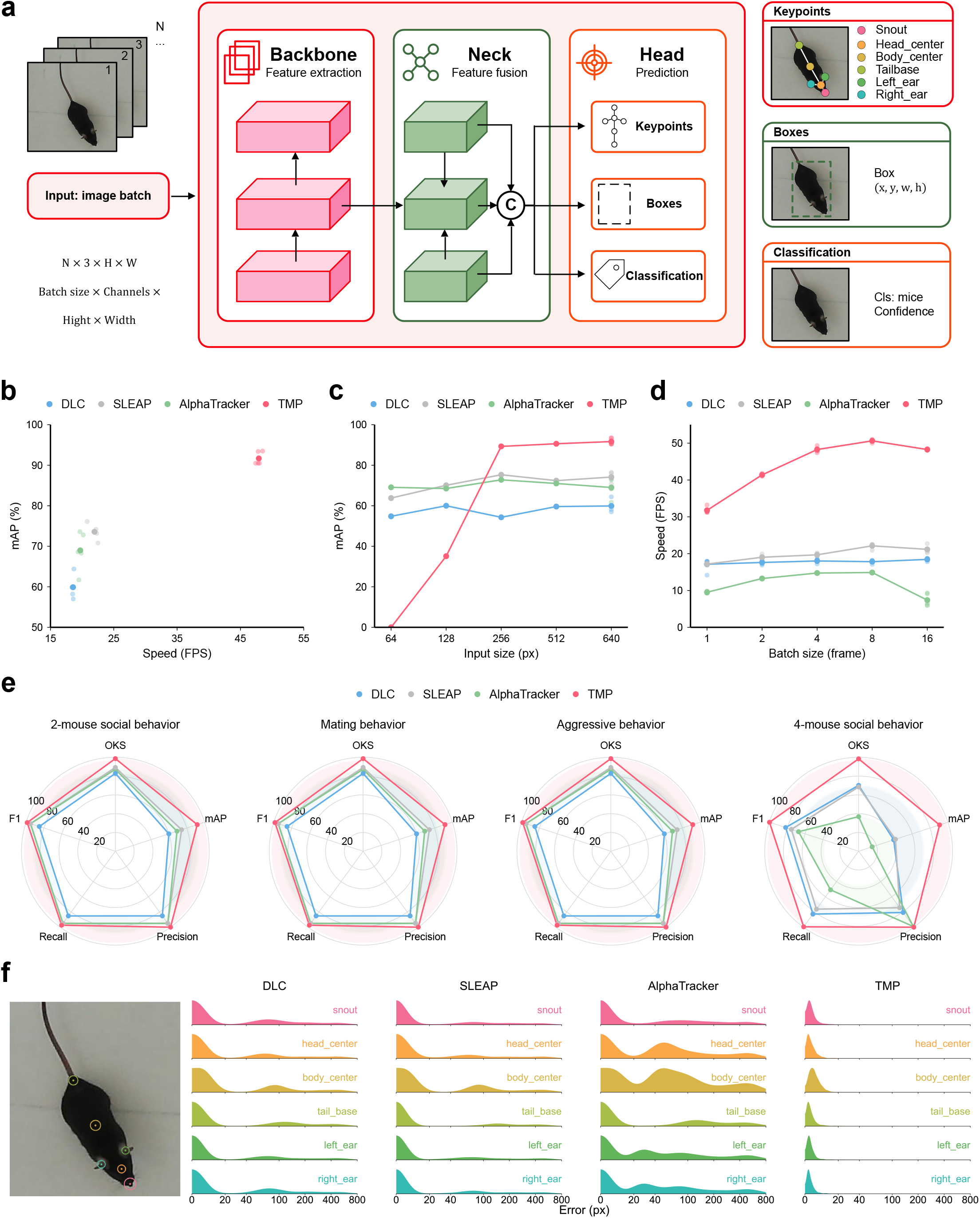
TMP enables accurate and high-throughput multi-animal pose estimation. **a**, TMP architecture. Batches of RGB images pass through a backbone, a multiscale feature-fusion neck, and prediction heads for keypoints, bounding boxes, and animal class. Training uses object keypoint similarity (OKS), complete intersection-over-union and distribution focal losses for localization, and binary cross-entropy for classification. **b**, Mean average precision (mAP) plotted against inference throughput for DeepLabCut (DLC), SLEAP, AlphaTracker, and TMP at the benchmark input settings. n = 5 times of repeated training per model. **c**, mAP across input resolutions from 64 to 640 pixels. **d**, Throughput at 640 × 640-pixel input resolution across batch sizes of 1, 2, 4, 8 and 16. N = 5 times of repeated inferences per batch size. **e**, Radar plots summarizing OKS, mAP, precision, recall and F1-score for two-mouse social behavior, mating behavior, aggressive behavior and four-mouse social behavior. **f**, Representative keypoint annotation and per-landmark pixel-error distributions for the snout, head center, body center, tail base, left ear and right ear. Error distributions are shown for DLC, SLEAP, AlphaTracker and TMP. Points and lines in **b**-**d** indicate the mean.

For identity tracking, TMP detections were passed to BoT-SORT, a multi-object tracking system^23^. A motion branch used Kalman filter-based state prediction to estimate the target’s position from preceding frames, whereas an appearance branch provided visual features for re-identification. The association incorporated motion and appearance information to maintain short-term trajectory continuity and to recover identities following occlusions. Pose sequences were subsequently reordered by tracked identity and converted into kinematic, within-animal postural, and social-interaction features (Fig. 1b, 3b).

To reduce reliance on manually specified behavioral factors, we developed an LLM-based factor-mining procedure (Fig. 1c). The system operated as a closed loop in which candidate factor formulas were proposed, calculated from behavioral time series, evaluated statistically, and either retained or rejected. Each iteration returned information to the LLM to guide subsequent proposals. Factors retained by this procedure provided the inputs to the downstream temporal classifier. The final behavior model used a three-stage stacked architecture (Fig. 1d). Separate first-stage models represented behavior over different temporal ranges, a second-stage meta-model integrated their predictions, and a third-stage temporal model incorporated local probability dynamics and temporal continuity. This architecture combines information across multiple temporal resolutions while reducing isolated frame-level classification errors.

### TMP provides accurate and efficient pose estimation across challenging social scenarios

TMP takes batches of three-channel RGB images as input and outputs keypoint coordinates, bounding boxes, and class predictions after backbone feature extraction and multiscale feature fusion (Fig. 2a; Supplementary Note 1). We first evaluated pose-estimation performance on a two-mouse natural social interaction dataset using mean Average Precision (mAP) as the primary metric. We compared TMP with three widely used frameworks, DeepLabCut^9^, SLEAP^8^, and AlphaTracker^10^, under a common input resolution of 640 × 640 pixels. At this resolution, TMP achieved a mAP of 92 ± 1.5%, outperformed DLC (60.7 ± 3.7%), SLEAP (73.7 ± 2.9%), and AlphaTracker (67.7 ± 6.0%) (Fig. 2b). Pose-estimation accuracy depended strongly on TMP input resolution (Fig. 2c). At 64- and 128-pixel inputs, TMP mAP was very low, whereas increasing the input resolution led to a steep increase, ultimately reaching a mAP above 90%. In contrast, DLC, SLEAP, and AlphaTracker showed substantially less variation across input resolutions within the configurations (Fig. 2c).

We next measured throughput at 640 × 640-pixel input resolution. TMP was the fastest among the four implementations across all tested batch sizes (Fig. 2d). Throughput increased with batch size, reaching 48.0 ± 0.6 frames s⁻^1^ at batch size 16. By contrast, the throughput of DLC and SLEAP changed relatively little across the tested batch sizes, processing approximately 18-21 frames s⁻^1^. AlphaTracker throughput increased from batch size 1 to 8 but dropped sharply at batch size 16.

To further evaluate complementary aspects of pose performance, we compared mAP, Object Keypoint Similarity (OKS), Precision, Recall, and F1-Score. TMP had the highest values across all five metrics in the overall social-behavior benchmark. SLEAP showed the second-most balanced profile, whereas AlphaTracker showed comparatively lower mAP and OKS, and DLC produced the lowest overall profile on the tested dataset (Fig. 2e, left).

We then examined three additional conditions—mating with extensive body overlap, aggressive interaction with rapid movement, and four-mouse social interaction. In the mating dataset, TMP achieved a mAP of 87.0 ± 3.5% and had the highest reported profile across all five-evaluation metrics. SLEAP and AlphaTracker showed lower pose-estimation performance under extensive overlap, and DLC had the lowest profile in this comparison.

During aggressive behavior, TMP again had the highest reported values across all five metrics. AlphaTracker performed better than SLEAP in this specific scenario, suggesting that relative model ranking depended on the form of behavioral difficulty. In the four-mouse condition, DLC and SLEAP showed broadly comparable profiles, whereas AlphaTracker had lower values on most metrics except precision (Fig. 2e). Collectively, these results indicate that TMP retained high pose-estimation performance across varying overlap, rapid movement, and increased animal numbers under the conditions evaluated here.

We further characterized localization errors for six body landmarks to visually depict the degree of disturbance in the model’s predictions and evaluate its positioning reliability. Error-density distributions showed distinct patterns among models. The 95th-percentile pixel-error radii of TMP were concentrated near zero across all six landmarks (12.63 pixels for the snout, 8.61 pixels for the head center, 10.98 pixels for the body center, 9.94 pixels for the tail base, 7.84 pixels for the left ear, and 8.24 pixels for the right ear), with narrower distributions than the other methods (Fig. 2f). DLC and SLEAP showed longer error tails, including observations extending beyond 400 pixels. AlphaTracker showed broader and, for some landmarks, multimodal distributions, particularly for the body and head centers. These distributions indicate that the TMP model is superior to the other three methods, not only in achieving the lowest systematic bias but also in substantially reducing estimation variance.

### Identity-preserving tracking reduces errors that confound behavioral attribution

Accurate pose estimation alone is insufficient for analyzing social behavior because behavioral events must be assigned to the correct individual throughout an interaction. This requirement becomes particularly challenging when animals physically contact each other, partially disappear from view, or rapidly change direction. We therefore assessed identity continuity after linking TMP detections with the tracking module (Fig. 3a).

**Fig 3.**
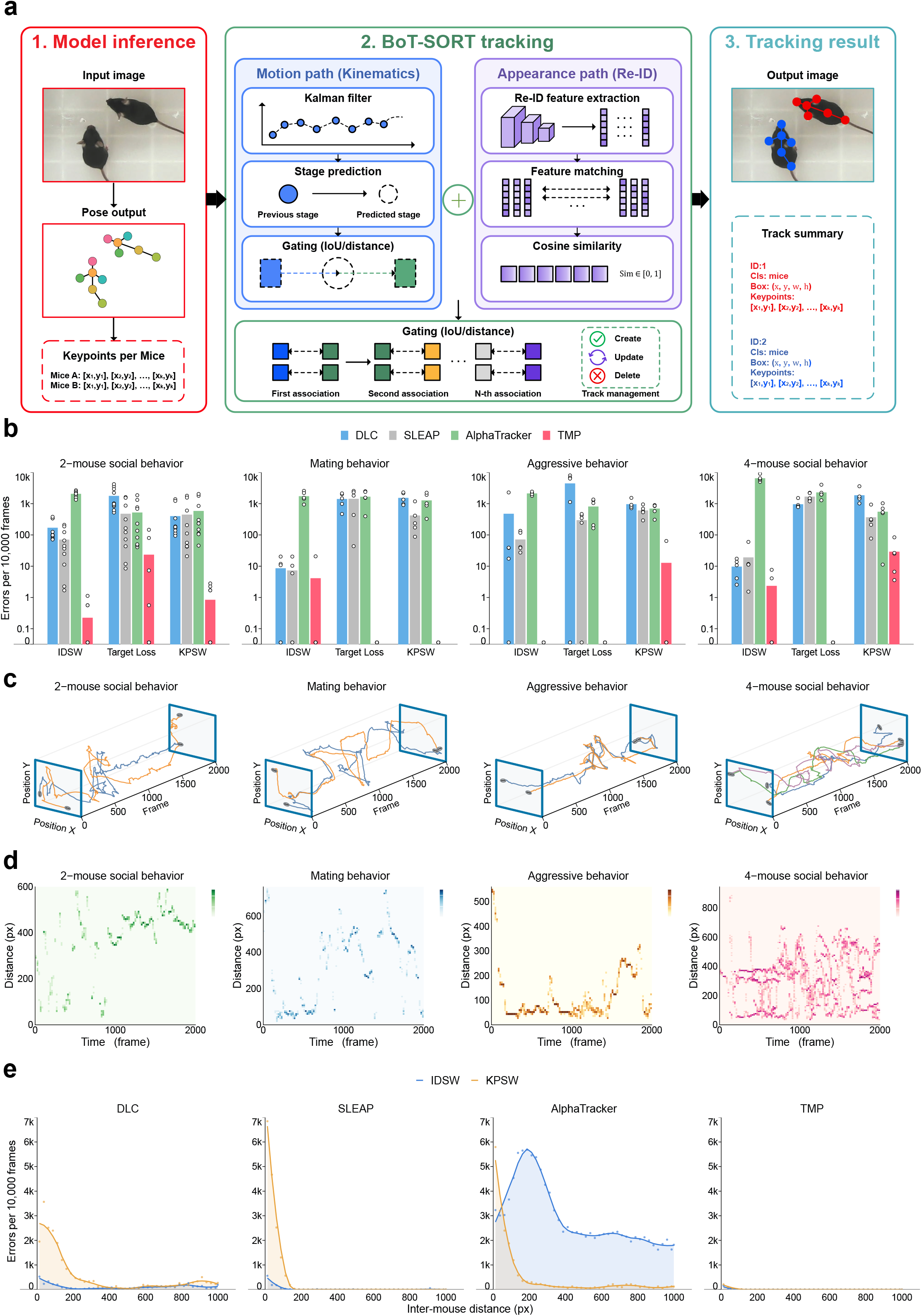
Identity-preserving tracking reduces assignment errors during multi-animal interactions. **a**, Tracking workflow. TMP detections and keypoints are passed to BoT-SORT, which combines Kalman-filter motion prediction, geometric association, and appearance-based re-identification to produce identity-linked pose trajectories. **b**, Errors of Identity switches (IDSW), target-loss events and keypoint switches (KPSW), normalized per 10,000 frames, for AlphaTracker, DLC, SLEAP and TMP across two-mouse social, mating, aggressive and four-mouse social recordings. Bars show mean ± s.e.m.; open circles denote individual recordings. n = 10 videos for 2-mouse social behavior; n = 5 videos for the rest conditions. **c**, Representative identity-linked trajectories plotted against frame number for the four interaction scenarios. **d**, Time-resolved inter-animal distance for the recordings shown in **c**. Lower values indicate closer spatial proximity between the tracked animals. **e**, Distance-dependent distributions of IDSW and KPSW for DLC, SLEAP, AlphaTracker, and TMP. Inter-mouse distance was used as a proxy for interaction proximity.

SABER integrates the TMP pose estimator with the BoT-SORT tracking framework, which combines motion prediction with appearance-based re-identification to maintain individual identity across frames. The tracker first predicts animal trajectories based on temporal continuity and subsequently incorporates appearance features to resolve ambiguous associations during close interaction^8^. We evaluated tracking performance using three error metrics that directly influence downstream behavioral interpretation: identity switches (IDSW) for cross-frame identity consistency, target loss for continuity of animal detection, and keypoint switches (KPSW) for incorrect assignment of body landmarks during tracking (Fig. 3b). Across two-mouse social interaction, mating, aggression, and four-mouse interaction, SABER showed the lowest error counts for IDSW, target loss, and KPSW in the plotted comparisons (Fig. 3b). AlphaTracker exhibited frequent identity switches, particularly in two- and four-mouse datasets. DLC showed fewer identity switches than AlphaTracker but more target loss and KPSW events, whereas SLEAP generally showed intermediate performance (Fig. 3b). These results indicate that TMP can more effectively suppress multi-level tracking errors, ranging from macroscopic trajectory interruptions to microscopic key-point confusion, in multi-target interactions and complex dynamic scenarios, thereby providing a more reliable data foundation for subsequent high-precision quantitative behavioral analysis.

Identity-linked x–y trajectories remained continuous throughout the representative interactions (Fig. 3c). These tracked identities enabled calculation of inter-animal distance over time (Fig. 3d). For example, aggressive interactions contained repeated epochs of small, rapidly changing inter-animal distance, consistent with repeated close-range interactions. The four-mouse recording showed a broader distribution of pairwise separations, which is consistent with dynamically changing spatial subgroups^3, 24^. Using inter-mouse distance as a proxy for interaction proximity and overlap severity, we found that IDSW and KPSW errors in the comparator methods increased markedly as animals approached one another (Fig. 3e). DLC and SLEAP showed pronounced KPSW accumulation at short inter-mouse distances, whereas AlphaTracker exhibited a particularly strong increase in IDSW that extended across a broader range of separations. By contrast, TMP maintained consistently low IDSW and KPSW rates across the full distance range, including the closest-contact regime in which body overlap and landmark ambiguity were greatest (Fig. 3e). These results indicate that SABER’s improved tracking performance is not simply driven by lower overall error rates, but is retained specifically under close-contact conditions that are most likely to disrupt identity continuity and keypoint assignment during social interactions.

### LLM-based behavioral factor mining generates interpretable representations from pose dynamics

Although individual-resolved pose trajectories provide detailed descriptions of animal movement, converting these high-dimensional signals into biologically meaningful behavioral representations remains challenging. We asked whether behavioral features could be automatically discovered from pose dynamics without relying exclusively on manually engineered descriptors. We therefore developed an LLM-based behavioral factor mining framework that generates, evaluates, and refines candidate behavioral factors through an iterative, closed-loop process. We annotated seven classic behavioral categories^3, 25, 26^, including approach, explore, climb, sniffing, self-grooming, stand, and blank for behavioral mining (Supplementary Note 2). Rather than directly using raw pose coordinates for classification, the system first generates candidate mathematical descriptors from available kinematic, postural, and inter-animal variables. We evaluated candidate factors on the behavioral dataset and retained those with one-versus-rest performance exceeding AUC ≥ 0.65 (Fig. 4a).

**Fig 4.**
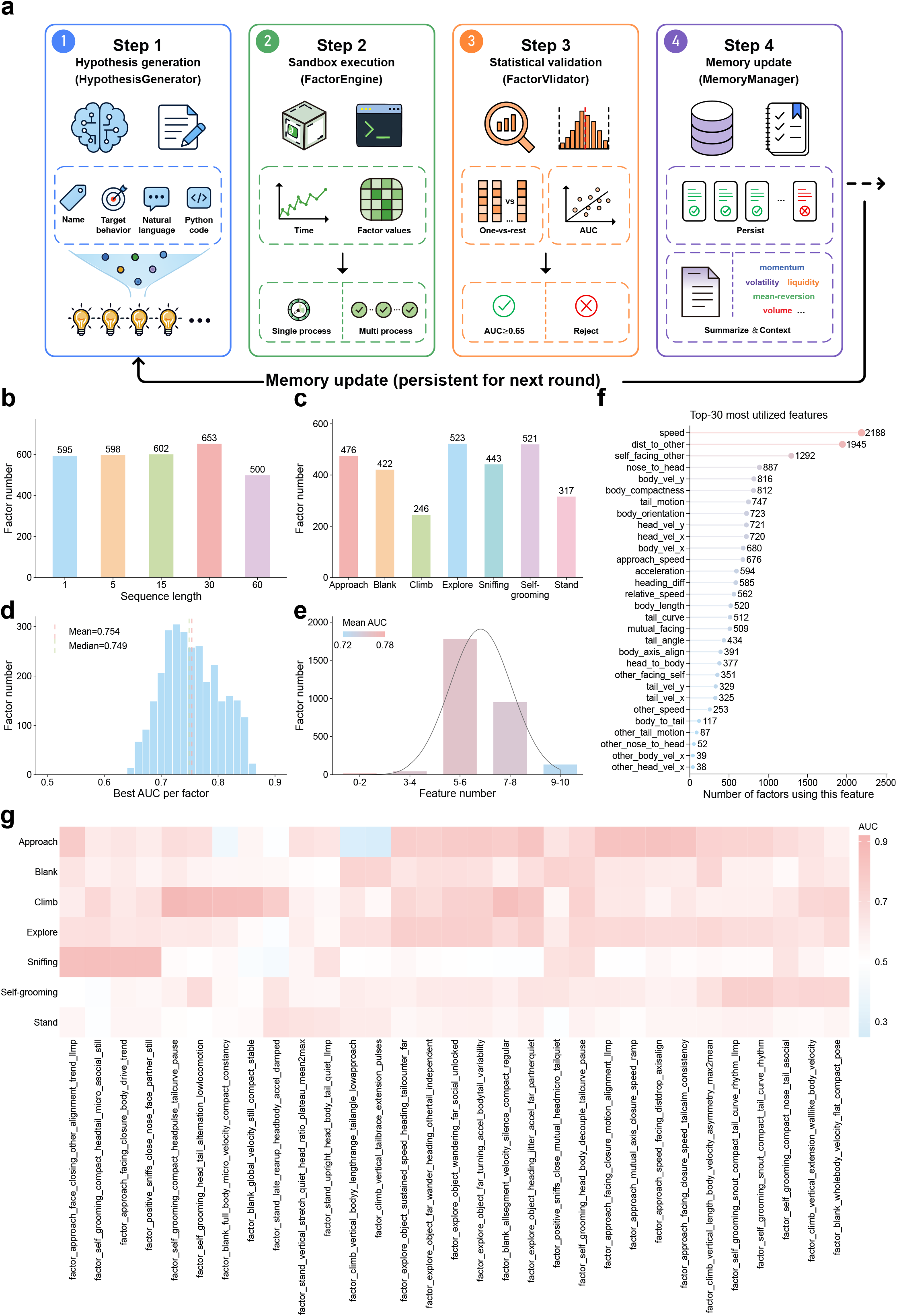
Closed-loop discovery and validation of interpretable behavioral factors. **a**, Four-step factor-mining workflow. A language model generates a named factor, target behavior, natural-language rationale, and executable Python expression; the factor is evaluated in a sandbox; one-versus-rest classification quantifies AUC and F1; and accepted or rejected proposals are summarized in persistent memory to guide the next round. **b**, Number of accepted factors at sequence lengths of 1, 5, 15, 30 and 60 frames. **c**, Number of accepted factors for approach, blank, climb, explore, sniffing, self-grooming, and stand. **d**, Distribution of the best one-versus-rest AUC obtained for each accepted factor. Dashed lines indicate the mean (0.754) and median (0.749). **e**, Distribution of the number of features used per factor. Bars show factor number within the indicated feature-number bins; the line shows a fitted normal density scaled to the counts. **f**, Top twenty features used in the accepted factor library; values indicate the number of factors containing each feature. **g**, Cross-behavior AUC matrix for the five leading factors selected for each target behavior. Rows denote behavior classes and columns denote factor definitions; color indicates one-versus-rest AUC.

Across multiple rounds of exploration, the system identified a large collection of behavioral factors spanning different temporal scales (Fig. 4b). The number of factors that met the classification criteria varied substantially across behavior classes. Behaviors of explore, self-grooming, and approach were associated with 523, 521, 476 effective factors, respectively, whereas sniffing, blank, stand, and climb were associated with 443, 422, 317, and 246 factors, respectively (Fig. 4c), suggesting that distinct behaviors are represented by different combinations of spatial and temporal features. The distribution of the best AUC obtained by each factor had a mean of 0.754 and a median of 0.749 (Fig. 4d). Most factors therefore provided moderate one-versus-rest discrimination under the factor-validation procedure, with a smaller set extending toward higher AUC values. Increasing the number of constituent variables did not produce a monotonic increase in factor AUC (Fig. 4e). Inspection of frequently used variables showed contributions from locomotor magnitude, inter-animal distance, body orientation, and relative alignment, etc. (Fig. 4f), indicating that the mining process repeatedly combined global locomotion, body configuration, and inter-animal spatial relationships. Notably, explore, climb, self-grooming, sniffing, and approach classes contained multiple factors with comparatively high one-versus-rest AUC values, whereas blank and stand showed less uniformly high factor performance across the displayed set (Fig. 4g). Together, these results indicate that LLM-based factor mining provides a scalable strategy for extracting interpretable behavioral representations while reducing dependence on manually designed feature engineering.

### Three-stage temporal modeling improves multi-animal behavior prediction

Behavioral states unfold across different temporal scales^27^. Brief changes in velocity or posture may represent transient behavioral events, whereas sustained movement patterns and social interactions often require longer temporal contexts^28^. We therefore developed a three-stage temporal prediction framework to integrate behavioral information across multiple time scales (Fig. 5a). The three-stage temporal prediction model (3-STBP) processes behavioral factors according to their temporal receptive fields. In the first stage, the model independently analyzes short-, medium-, and long-duration factors to capture behavior-specific temporal signatures. The model then integrates the resulting predictions through a meta-learning framework that combines information across temporal resolutions. Finally, a temporal refinement module incorporates sequential continuity to generate frame-level behavioral predictions.

**Fig 5.**
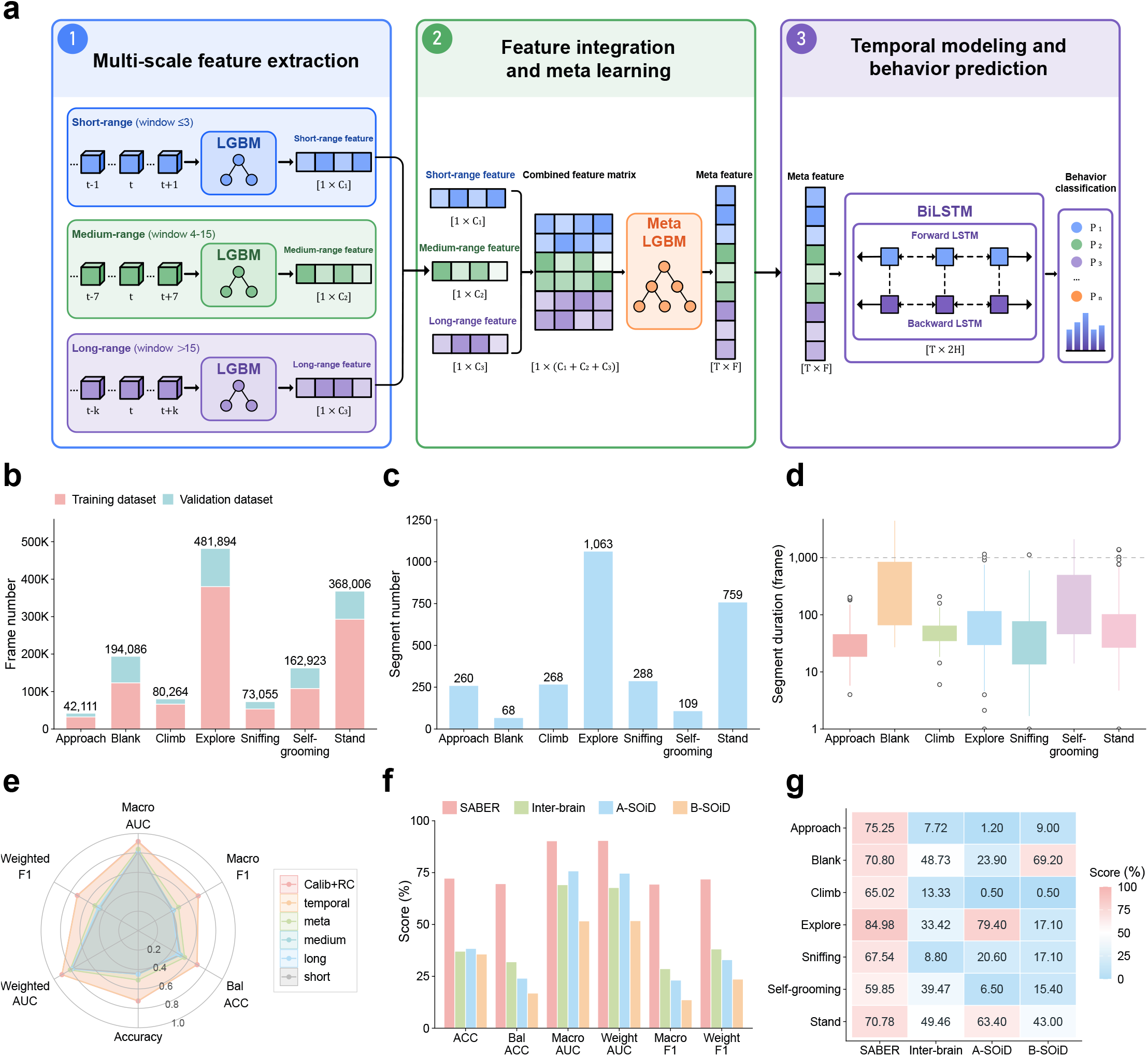
Multiscale temporal integration improves behavior classification. **a**, The architecture of the three-stage temporal behavior prediction. Short-, intermediate-, and long-range factors are first processed by separate LightGBM classifiers. Their class-probability vectors are concatenated and integrated by a meta-LightGBM model, after which a 31-frame sliding window supplies temporal context for final frame-level prediction. **b**, Training and validation frame counts for the seven annotated behavior classes. **c**, Segment number count for each behavior classes. **d**, Per-class segment-duration distributions on a logarithmic y axis. Boxes show the median and interquartile range, whiskers extend to 1.5 times the interquartile range, and points show values outside the whiskers. **e**, Radar plots summarizing accuracy, balanced accuracy, macro AUC, weighted AUC, macro F1-Score and weighted F1-Score across the short-, intermediate- and long-range models, the meta-model, temporal model and calibrated rule-corrected model (Calib+RC). **f**, Overall validation performance of SABER, Inter-brain, A-SOiD and B-SOiD across accuracy, balanced accuracy, macro AUC, weighted AUC and macro F1. **g**, Per-class precision for the four behavior-analysis methods. Values within cells are percentages.

The behavior dataset comprised 40 annotated two-mouse social-interaction videos, yielding 2,815 continuous behavioral segments. The class distribution was strongly imbalanced: explore and stand behaviors accounted for about 64% of frames, whereas approach accounted for <3% (Fig. 5b). Behavioral segments also differed markedly in duration, with blank and self-grooming segments being relatively long (Fig. 5c, d). We compared successive stages using macro F1-Score, balanced Accuracy, Accuracy, weighted AUC, weighted F1-Score, and macro AUC. The short-, medium-, and long-range first-stage models showed similar overall profiles, with no single temporal range dominating all metrics. The meta-model provided comparatively modest additional improvement. The largest increase, particularly for weighted AUC and weighted F1-Score, occurred after temporal context was incorporated in the third stage, and peaked with the further calibration and resampling (Calib+RC) approach (Fig. 5e). These results indicate that behavioral classification benefits from explicitly modeling the hierarchical temporal structure of animal actions rather than treating each frame as an independent observation.

We next compared SABER with alternative behavioral classification approaches, including supervised classifiers-A-SoiD^12^, SVM-based classification (Referred to as Inter-brain)^3^, and unsupervised B-SoiD^29^. SABER achieved overall accuracy above 70%, macro-AUC close to 90%, and balanced accuracy and macro F1 of approximately 70% in the displayed benchmark (Fig. 5f). B-SOiD had the lowest displayed scores, with all reported aggregate metrics below approximately 25%. A-SOiD achieved an overall Accuracy of approximately 38%. Inter-brain was the strongest of the three comparator implementations, with macro and weighted AUC values in the approximately 60–70% range but balanced Accuracy of approximately 30%. Class-level results further showed that the relative advantage varied by behavior (Fig. 5g). For explore behavior, the accuracies were 84.98% for SABER, 33.42% for Inter-brain, and 79.4% for A-SOiD. In terms of approach, SABER achieved 75.25%, whereas the other models were below 10%. Together, these data demonstrate that SABER improved behavioral prediction performance.

### SABER detects a stress-associated behavioral profile and aligns behavior with neural activity

To make the analysis workflow usable without assembling separate computational components, we implemented SABER as a locally installable desktop application (Fig. 6a). The interface allows users to import animal videos and perform pose estimation, identity-preserving tracking, behavioral factor mining, and frame-resolved behavior prediction within a unified environment. Pose trajectories, individual identities, and predicted behavioral timelines can be inspected directly in the application, enabling researchers to proceed from raw video to quantitative behavioral output without constructing separate computational pipelines (Fig. 6a).

**Fig 6.**
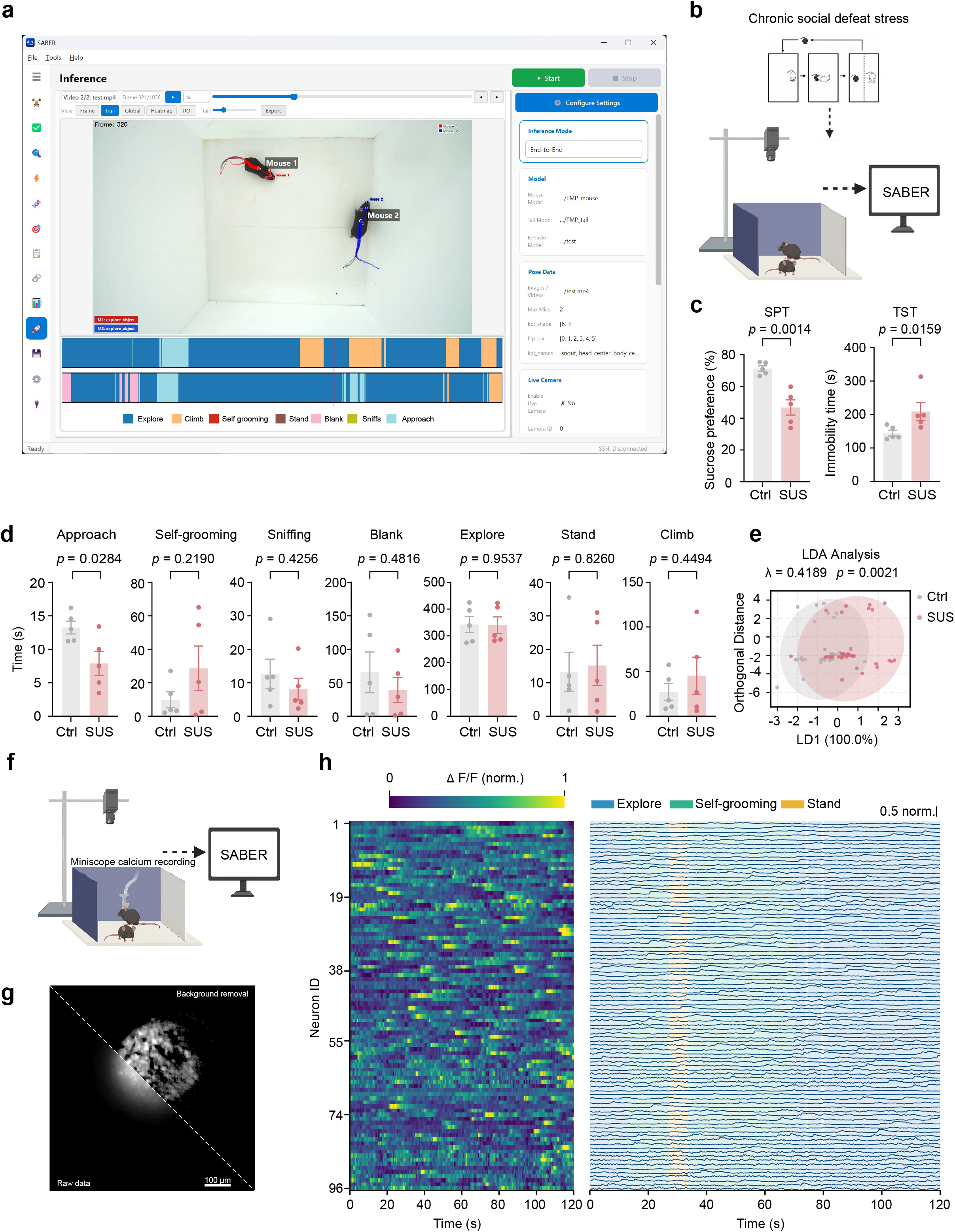
SABER identifies a stress-associated behavioral profile and aligns behavior with miniscope calcium activity. **a**, SABER desktop interface showing identity-linked pose estimates, behavior labels, and frame-resolved timelines for two interacting mice. **b**, Chronic social defeat stress and social-interaction recording workflow used for SABER analysis. **c**, Decreased sucrose preference ratio in the sucrose preference test (SPT) and increased immobility time in the tail suspension test (TST) in stress-susceptible ^38^ mice compared with control (Ctrl) mice. n = 5 mice per group. Student t-test for SPT, t_(8)_ = 4.773, *p* = 0.0014; Mann-Whitney test for TST, *p* = 0.0159. **d**, Time spent in approach, self-grooming, sniffing, blank, explore, stand, and climb during the 10-min social-interaction recording. Student t-test, for approach, t_(8)_ = 2.669, *p* = 0.0284; for self-grooming, t_(8)_ = 1.334, *p* = 0.219; for sniffing, t_(8)_ = 0.8393, *p* = 0.4256; for blank, t_(8)_ = 0.738, *p* = 0.4816; for explore, t_(8)_ = 0.0599, *p* = 0.9537; for stand, t_(8)_ = 0.2272, *p* = 0.826; for climb, t_(8)_ = 0.7952, *p* = 0.4494. **e**, Linear discriminant analysis of the seven behavioral-duration variables. Points show observations from Ctrl and SUS groups, crosses indicate group centroids, and shaded regions show 95% confidence ellipses (Wilks’ λ = 0.4189, *p* = 0.0021). **f**, Concurrent miniscope calcium imaging and overhead social-behavior recording, with the behavioral video processed by SABER. **g**, Representative miniscope field of view shown before and after background removal. Scale bar, 100 μm. **h**, Calcium activity from 96 accepted neuronal components during a representative 120-s recording. The heat map and traces are ordered by the original neuron identifier and aligned row by row. Color indicates within-neuron normalized ΔF/F. Shading denotes SABER-predicted explore, self-grooming, and stand epochs; vertical scale bar, 0.5 normalized units. In **c**,**d**, bars show mean ± s.e.m. and points denote individual mice. \**p* < 0.05, \*\**p* < 0.01; ns, not significant.

We next tested whether SABER-derived measurements could identify behavioral differences in mice exposed to chronic social defeat stress, a mouse model of depression^30, 31^. After chronic social defeat stress, we paired stress-susceptible and control mice with a conspecific and recorded a 10-min free-interaction session (Fig. 6b). Compared with controls, SUS mice showed reduced sucrose preference and increased immobility time in the TST (Fig. 6c), confirming the depression-related phenotype. We then quantified the time spent in each of the seven predefined behavioral categories. SUS mice spent significantly less time approaching their social partner than control mice (Fig. 6d). Self-grooming was numerically increased, whereas sniffing was numerically reduced, although neither difference reached statistical significance. Because the phenotype may involve coordinated changes across several behaviors, we also evaluated the seven behavioral measurements using linear discriminant analysis (LDA). The resulting discriminant dimension separated the control and SUS behavioral profiles along the discriminant axis (λ = 0.4189, *p* = 0.0021; Fig. 6e). Thus, SABER-derived behavioral composition captured a multivariate signature that distinguished stress-susceptible mice from controls.

Finally, we synchronized SABER-derived behavioral outputs with miniscope calcium imaging (Fig. 6f). Background removal and denoising produced cellular-resolution imaging data, from which 96 neuronal traces were extracted in the representative recording (Fig. 6g). Neural activity and SABER-predicted behavior were registered to a common time base, allowing activity patterns to be inspected during explore, self-grooming, and stand epochs (Fig. 6h). These results establish a unified workflow in which disease-related behavioral phenotypes can be quantified and subsequently related to population neuronal dynamics at frame-level temporal resolution.

## Discussion

Reliable quantification of social behavior requires maintaining information across several stages of measurement. Pose errors alter body-geometry estimates, identity errors assign measurements to the wrong animal, and both errors propagate into the variables used for behavioral classification. SABER was designed around this dependency by linking multi-animal pose estimation, identity-preserving trajectory reconstruction, quantitatively screened behavioral factors, and temporal classification within a single workflow. Across the social-interaction conditions evaluated here, this organization produced accurate pose estimates, fewer observed tracking errors, and higher behavioral prediction accuracy that captured both individual actions and inter-animal relationships. The model weights have been publicly released, thereby enabling researchers to either apply them directly, perform transfer learning with limited data, or train models de novo using their own datasets. The choice of strategy can be made according to the researchers’ specific requirements.

A key advance of SABER is its approach to behavioral factor discovery. Traditional behavioral analysis often relies on manually defined features selected based on prior knowledge of specific behaviors^12, 21^. Although expert-designed features remain valuable, they may not scale efficiently as behavioral repertoires become more complex or as subtle phenotypic differences require identification. Conversely, fully unsupervised approaches can reveal latent movement structures but may generate representations that are difficult to interpret biologically^5^. The LLM-based factor mining framework in SABER offers an intermediate strategy that combines automated exploration with quantitative validation. Rather than replacing behavioral expertise, this approach substantially reduces the time required to formulate and screen complex behavioral descriptors. More importantly, because the language model can draw on broad prior knowledge while exploring combinations of kinematic, postural, and social variables, the resulting factors may expose non-obvious relationships between measurable features and behavioral states. These hypotheses remain subject to empirical validation, but they can provide investigators with interpretable starting points for mechanistic analysis and thereby extend the role of SABER from automated behavior prediction to data-driven behavioral hypothesis discovery.

The temporal organization of behavior represents another important consideration for automated behavioral analysis. Animal behaviors are not independent frame-by-frame events but hierarchical processes that unfold across multiple time scales^25, 27^. Short-lived changes in posture or velocity may indicate transient actions, whereas prolonged patterns may represent sustained behavioral states. The multiscale temporal prediction strategy implemented in SABER explicitly models this temporal hierarchy by integrating information from different behavioral time windows. This design improves behavioral prediction while maintaining interpretability by allowing individual temporal components to contribute to the final classification.

Reliable behavioral inference is only possible when posture and identity are measured accurately. Another advance of SABER is the ability to maintain reliable behavioral attribution during complex social interactions. Previous computer vision approaches have substantially improved the automated measurement of animal behavior, particularly for individual animals^7, 32^. However, social behaviors add complexity because the behavioral output of interest often emerges from interactions among individuals. During physical contact, rapid movement, or partial occlusion, errors in pose estimation and identity assignment can propagate into downstream behavioral interpretation. The TMP module of SABER is optimized for overlapping animals.

Because severe-overlap frames are rare and hard to annotate accurately, we developed a Geometry-Aware Mouse Synthesis Algorithm that generates many pixel-accurate synthetic overlap examples with controlled overlap ratios and relative orientations. TMP also incorporates Semantic-Aware Grouping Attention, which maintains cross-image semantic prototypes of body structures and uses these representations to recover partially occluded landmarks. By integrating high-resolution pose estimation with identity-preserving tracking, SABER therefore provides a framework for assigning behavioral events to specific individuals throughout dynamic interactions. This capability is particularly important for studies investigating social decision-making, dominance relationships, affiliative behaviors, and disease-associated alterations in social phenotypes.

SABER complements, rather than replaces, existing behavioral analysis frameworks. Current pose estimation platforms, including DeepLabCut, SLEAP, and related approaches, have established the foundation for markerless behavioral measurement by enabling flexible extraction of animal body configurations^5^. Approaches such as Lightning Pose further improve robustness by incorporating temporal constraints and semi-supervised learning^33^, whereas behavioral discovery methods such as A-SOiD emphasize efficient annotation and active learning^12^. SABER builds upon these advances by focusing on a different but complementary problem: how to transform reliable individual-level behavioral measurements into scalable and interpretable descriptions of complex social behaviors. Future integration with more general behavioral foundation models may further enhance cross-species applicability and reduce dependence on task-specific training.

Several limitations should be considered. First, SABER currently relies primarily on two-dimensional video recordings and predefined anatomical landmarks. Although this representation is sufficient for many behavioral analyses, three-dimensional reconstruction from multi-view imaging would provide additional information about body orientation, physical contact, and spatial organization during social interactions. Second, although the LLM-based factor mining framework reduces dependence on manual feature engineering, the quality of the discovered factors still depends on the underlying feature representation and the objective-function design.

Incorporating larger behavioral datasets and more advanced multimodal models may further improve automated behavioral representation learning. Third, we developed and evaluated SABER primarily in laboratory mouse social paradigms. Extending the framework to other species, naturalistic environments, and long-duration recordings will require improved generalization strategies, including self-supervised learning and large-scale pretraining.

Looking forward, integrating quantitative behavioral analysis with neural recordings represents an important direction for systems neuroscience^34, 35^. Emerging approaches that jointly model brain activity and behavioral dynamics highlight the need for behavioral representations that are sufficiently precise, interpretable, and scalable^5^. By providing individual-resolved behavioral trajectories and automatically derived behavioral representations, SABER may facilitate the investigation of how neural circuits generate, coordinate, and adapt social behaviors, particularly when employed alongside closed-loop neuromodulation paradigms.

Together, our work establishes a computational framework for quantitatively analyzing social behavior, bridging the gap between raw behavioral videos and interpretable behavioral states. By combining accurate measurement with automated representation discovery, SABER provides a foundation for systematically investigating social behavior across physiological, genetic, and disease contexts.

## Methods

### Mice

Male and female C57BL/6J mice aged 8–10 weeks were obtained from the Experimental Animal Center of Southern Medical University. Mice were group-housed on a 12-h light–dark schedule, with lights on at 08:00 and off at 20:00, at 22 ± 2 °C and 40–60% relative humidity. No more than five animals were housed per cage, with food and water available ad libitum. Mice were handled twice daily for 3–4 days before behavioral experiments. Experimental procedures were approved by the animal ethics committee of Southern Medical University.

### Datasets

A dataset combining keypoint and behavior annotations under standardized conditions was constructed and openly released (see Data availability), as public datasets provide either relatively uniform scenes or do not cover both keypoint annotations and behavioral labels. This dataset complements existing resources and enables joint pose–behavior analyses under various social conditions.

### Behavioral video acquisition for training and validation sets

Four types of videos were used to develop and evaluate the pose-tracking pipeline: spontaneous two-mouse social behavior (40 videos, each for 10 min), aggressive behavior (12 videos, lasting for 22-76 sec), mating behavior (7 videos, 23-113 sec), and spontaneous four-mouse social behavior (5 videos, each for 10 min). For two-animal recordings, mice moved freely in a white plastic open-field arena (42 × 42 × 30 cm). Four-mouse recordings were conducted in a 27 × 39 × 27.5 cm arena. Videos were acquired from above under white LED illumination of 100–300 lux using a KM2010-1080P camera (Kaimeng vision) controlled with VideoCap software. The camera supported acquisition at up to 1,920 × 1,080 pixels and 30 frames s⁻¹. Animals acclimated to the arena for 30 min before recording. Ambient temperature and humidity were maintained at 22 ± 2 °C and 40–60%, respectively. Two- and four-mouse social recordings were acquired at 1,920 × 1,080 pixels for 10 min with an image scale of 0.4295 mm pixel⁻¹. Aggressive and mating behavior datasets were stored at 1,280 × 720 pixels with an image scale of 0.5048 mm pixel⁻¹. The training/validation split and the per-set frame counts are summarized in Supplementary Note 1.

### Keypoint definition and pose annotation

Six anatomical landmarks were defined: snout, head_center, body_center, tail_base, left_ear, and right_ear. The head center was operationally defined as the center of the triangle formed by the two ears and snout. The skeleton connected the snout and both ears to the head_center, the head_center to the body_center, and the body_center to the tail_base. To increase the number of raw features, we additionally include keypoints for the tail portion: tail_start, tail_middle, and tail_end. The body and tail parts are annotated as two distinct categories, which are used to train two separate TMP models for inference on the body and tail parts, respectively. Trained annotators used LabelMe to annotate keypoint coordinates, animal bounding boxes, and identity/group information. Fully occluded keypoints were left unlabeled. Annotations were exported in JSON format. A total of 3000 randomly sampled frames, consisting of 1000 frames from the 2-mouse social behavior dataset, 1400 from the aggressive behavior dataset, and 600 from the mating behavior dataset, were used for keypoint annotation. The training-to-validation dataset ratio was 3:1.

### Behavioral categories and frame-level annotation

The behavioral classification dataset comprised 40 10-min videos of freely interacting mouse pairs. A single trained researcher annotated all recordings using Behavior Annotator and exported the onset and offset times of each behavior in TXT format. Labels described an individual mouse’s state rather than a dyadic interaction class. Seven mutually exclusive frame-level behavior classes, including approach, blank, climb, explore, sniffing, self-grooming, and stand, were defined based on previous reports^3, 26^. Approach denoted directed movement toward the partner; blank denoted immobility with all four paws on the floor; climb denoted rearing with an attempt to climb the arena wall; sniffing denoted close-range sniffing of the partner; self-grooming denoted self-directed grooming; stand denoted a bipedal upright posture; and explore denoted sustained movement without an obvious direction. The frame-level class distribution of the behavioral dataset was summarized in Supplementary Note 1.

### SABER system architecture and software implementation System overview

SABER is a locally deployable, open-source platform for markerless mouse behavior analysis. It combines keypoint detection, multi-object tracking, feature construction, LLM-based factor mining, and multiresolution behavior classification. The same trained models support offline batch analysis of recorded videos and streaming analysis of live camera feeds. Functionality is exposed through a desktop graphical user interface, command-line interface (CLI), and Python application programming interface (API).

The analysis chain contains four principal computational modules. First, TMP, a single-stage keypoint detector, localizes anatomical landmarks in each frame. Second, a BoT-SORT-based tracker links frame-level detections into identity-indexed trajectories. Third, the secondary-feature engine converts keypoint coordinates into standardized kinematic, postural, and social descriptors (Supplementary Note 1). Fourth, the factor-mining and behavior-prediction engine uses an LLM to propose executable factor definitions and a three-stage classifier to convert the resulting factor matrix into frame-level behavior probabilities and labels.

### Analysis workflow

Video input. The user supplies one or more video files, an image directory, or a live camera stream.

TMP pose estimation. For every detected animal, TMP returns 6 keypoint coordinates and confidence scores. An optional tail-specific pose model can run in parallel and fuse with the body-pose output.

Identity association. Frame-level detections are linked over time by a BoT-SORT-based tracker. The tracker maps stable track identifiers to the target animal and its partner for downstream feature calculation.

Secondary-feature calculation. Identity-linked landmarks are converted into normalized coordinates, centered finite-difference velocities and accelerations, angular variables, and engineered social or kinematic descriptors.

LLM-based factor mining. On labeled data, the LLM proposes short, executable code fragments that define candidate factors. Each candidate is evaluated in a restricted sandbox and screened quantitatively before being added to the factor library. Genetic-programming-style mutations and parameter tuning can further extend the library.

Three-stage behavior prediction. Validated factors are partitioned by temporal receptive field into short-, intermediate-, and long-range groups. Group-specific gradient-boosting models generate posterior probabilities, a meta-learner combines the three groups, and a temporal second-level model refines the frame-level sequence.

Output and spatial summaries. SABER writes a frame-resolved behavior timeline, calibrated class probabilities, rendered pose and behavior overlays, identity-specific trajectories, occupancy maps, user-defined region-of-interest summaries, and quantitative model reports.

Offline and streaming modes share the same serialized model bundle. In offline mode, recorded videos or previously extracted keypoints are processed at full precision, and body and tail pose models can run on parallel CUDA streams. In streaming mode, each arriving frame is processed by FP16 pose estimation and identity tracking, while behavior classification runs asynchronously in a separate process that consumes a queue of identity-linked keypoints. Predictions are overlaid on the video and checkpointed periodically as frame-level CSV files and incrementally updated behavior rasters. Separating the acquisition–tracking loop from the prediction process prevents behavior classification from blocking frame capture.

### Data interfaces between modules

Modules exchange explicit serialized objects rather than relying only on shared in-memory state. For a frame containing N animals and K=6 landmarks, TMP produces a coordinate tensor of shape [N, K, 2] and a confidence matrix of shape [N, K]. The tracker returns identity-indexed trajectories, which are stored as per-animal keypoint files after canonical track mapping. MouseBehaviorDataset converts these files into a dense float32 feature matrix *X* ∈ ℝ*^T^*^×*D*^, an optional label vector *y* ∈ ℤ*^T^*, and a length-D list of feature names (flat attributes). A behavioral factor is represented as a pure function mapping a window *W* ∈ ℝ*^L^*^×*D*^ to one scalar in row mode, or a batch *W* ∈ ℝ*^N^*^×*L*×*D*^ to an N-vector in batch mode. Stacking F factors across T frames yields a factor matrix in ℝ*^T^*^×*F*^ ; the three-stage classifier converts this matrix into group-level posterior probabilities in ℝ*^T^*^×*C*^, a *T×3C* meta-representation, and a final label sequence in ℤ*^T^*.

### User interface and interactive operation

The desktop application was implemented with PySide6, the official Python binding for Qt 6, from a single cross-platform codebase for Windows, Linux, and macOS. Long-running operations, including pose-model training, factor mining, behavior-model training, and inference, execute in background QThread workers so that the interface remains responsive and operations can be monitored or interrupted. The application runs locally and does not require a separate application server.

The main window contains a collapsible navigation sidebar organized around training, factor mining, factor engineering, prediction, and utility functions. Training panels launch pose training and three-stage behavior-model training. Factor-mining panels support interactive and resource-aware batch mining. Factor-engineering panels provide evolution, parameter tuning, library management, correlation analysis, and factor analysis.

Prediction panels execute inference and validation. Utility panels provide configuration editing and optional SSH-based remote execution. Each panel uses a common layout with typed controls on the left and plots, tables, and streaming logs on the right.

Frame-level behavior timelines and plotted variables are exported as CSV. Factor definitions, validation reports, pipeline manifests, and identity-linked trajectories are stored as JSON. Run configurations are stored in YAML, and confusion matrices, behavior rasters, occupancy maps, and calibration plots are stored as PNGs. Inference additionally saves rendered pose and behavior overlays that can be assembled into annotated videos.

### Application programming interfaces and deployment

The same functionality is accessible programmatically. A pose-predictor object loads a trained pose-model directory and converts video frames into keypoint arrays. A tracker associates detections into identity-indexed trajectories. MouseBehaviorDataset converts trajectories into the standardized feature matrix and feature-name index. A hypothesis-generator object drives LLM-based factor proposal and validation for a selected target behavior. InferencePipeline loads a trained run bundle and executes the three-stage predictor to return a frame-level behavior timeline. These objects can be composed into an end-to-end script or invoked independently.

For batch or headless use, each functional mode is exposed as a CLI entry point. Dedicated commands cover poses training, factor discovery, three-stage behavior-model training, and inference on new recordings.

Additional entry points include factor mutation, LLM-based parameter tuning, correlation-based redundancy filtering, factor analysis, factor library management, and held-out validation. GUI panels are lightweight wrappers around the corresponding CLI functions and therefore write the same outputs.

Hyperparameters are managed declaratively in hierarchical YAML files. The source implementation stores LLM endpoint, dataset, preprocessing and output settings in common.yaml; discovery thresholds and loop control in discovery.yaml; and classifier or temporal-model settings in validation.yaml. JSON resources store dataset manifests, class mappings, and behavioral priors. Command-line flags can override individual fields, and the fully resolved configuration is persisted with each run.

### Data model and storage format

A dataset configuration is a list of segment records, each pointing to the target- and tail-keypoint directories for a recording, the maximum number of co-housed animals, and the behavioral annotation file. Loading the configuration constructs a MouseBehaviorDataset containing a dense *T×D* feature matrix, an aligned T-vector of labels, and a feature-name index. Each factor-record stores a name, a natural-language description, a target behavior, executable code, an evaluation mode (row or batch), a temporal window length, and measured discrimination metrics. A collection of factor records constitutes a reusable factor library.

All files required to reproduce a run—including configurations, model weights, normalizers, factor definitions, and class mappings—are placed in a single run directory that can be copied, archived, or version-controlled.

Preprocessed datasets and normalized factor matrices are cached with content-derived hashes based on the input data, factor code, and normalization settings. Identical inputs therefore reuse the same intermediate objects. The merged configuration and factor library are stored with the model weights so that reported predictions can be traced to the exact factors and settings used to generate them.

### Pose-estimation and identity-tracking models TMP architecture

TMP follows a backbone–neck–head architecture. The network was designed for small, visually similar animals that frequently overlap. It combines feature-dependent receptive fields with attention-based cross-scale fusion to reduce ambiguity in crowded frames.

An RGB input tensor first passes through an MSPatchEmb stem for feature embedding, normalization, and nonlinear transformation. Four successive stride-2 stages produce feature maps C1–C4 at 1/4, 1/8, 1/16, and 1/32 of the input resolution. Dynamic-convolution modules are inserted at implementation layers 3, 5, 7, and 9. Each module mixes 3×3 and 5×5 convolutional responses with feature-dependent weights, thereby adapting the effective receptive field to the current image representation. At the deepest stage, a spatial-pyramid-pooling-fast (SPPF) module concatenates pooled features from multiple scales. A 1×1 convolution compresses the channels before a fusion block containing multi-head attention and a feed-forward transformation.

The neck performs top-down multiscale fusion. Deeper semantic features are upsampled and concatenated with higher-resolution features to combine global context with finer spatial detail. Fused branches are standardized to 256 channels. Additional upsampling or downsampling aligns the spatial dimensions before the prediction head. Dynamic residual–bottleneck modules are placed after fusion nodes.

The head contains separate classification, bounding-box, and keypoint branches, each built from independent 3×3 convolutions with SiLU activation. The classification branch predicts foreground and background channels. The box branch predicts the center coordinates (*c_x_*,*c_y_*), width, and height. The landmark branch predicts one confidence heatmap for each of the six keypoints. Ground-truth keypoints are represented by two-dimensional Gaussian heatmaps centered at the manual annotation, with a scale-adaptive standard deviation. A Proto branch upsamples the highest-resolution head input by a factor of two to generate auxiliary pose features for spatial refinement. All outputs are optimized jointly with their corresponding loss terms.

### Semantic-Aware Grouping Attention

Semantic-Aware Grouping Attention (SAGA) models relationships between spatially separated but semantically related image tokens. SAGA combines a cross-stage-partial bottleneck pathway with a transformer-style context pathway. The outer module follows a C3k2-like two-branch design. A 1×1 convolution divides the input channels. The main path contains bottleneck blocks with an optional residual connection controlled by the add parameter, followed by ConvT1 operations for spatial or channel adjustment, then GeLU and 1×1 convolutions. The second path bypasses the main transformations, and the two branches are concatenated at the output.

The context pathway contains a Hierarchical Context Routing Block (HCRB), which comprises Semantic-Aware Sparse Aggregation (SASA), Partitioned Self-Attention (PSA), Global Cross-Cluster Attention (GCCA), and a final 1×1 channel-mixing convolution. Padding preserves spatial alignment during grouping, and a PushBack operation restores grouped tokens to their original positions in the image.

### SASA

SASA maintains **M** global learnable token centers **C** ={*cj* ∈ ℝ*^d^*}*j*=1⋯*M* across the training set rather than recomputing cluster centers independently for every image. For image tokens X = {x_i_ ∈ ℝd}_i = 1 ⋯ N_, cosine similarity is calculated as**D** = ℳ({*x_i_*}, {*c_j_* })//**D** ∈ ℝ*^N^*^×*M*^.Each token is assigned to the center with the largest similarity, forming semantic groups{G*_j_*} = {{*x^i^* ∣ ∗ *arg max_k_* **D***_ik_* = *j*}}. During training, provisional centers are calculated from the current assignments and the persistent centers are updated by exponential moving average, 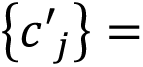 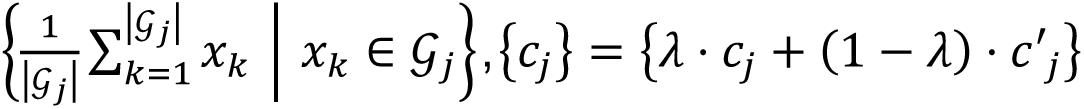. At inference, the learned centers are reused without per-image center optimization. To avoid large differences in group size that would reduce parallel efficiency, grouped tokens are concatenated and repartitioned into fixed-size subgroups before attention is applied.

### Partitioned Self-Attention

For a balanced subgroup*S_j_* , PSA forms learned*Q_j_*, *K_j_* and *V_j_* projections. To reduce discontinuities introduced by semantic regrouping, a query subgroup attends to its own keys and values and those of the adjacent subgroup *j+1*,

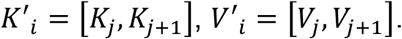

Multi-head scaled dot-product attention is then applied, 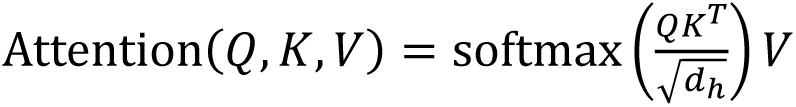, and PushBack maps the output tokens to their original spatial positions to form *X_local_*. This step retains local semantic interactions while preserving the geometry needed by subsequent convolutional layers.

### Global Cross-Cluster Attention

GCCA exposes each local subgroup to the persistent global centers. A local subgroup supplies the query, whereas the global center matrix supplies the keys and values,

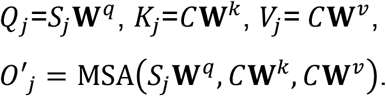

PushBack restores global-context outputs to image order, forming *X_global_*. Local and global outputs are combined by residual addition and a 1×1 convolution,

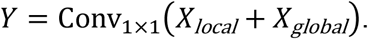

Because *M* is substantially smaller than *N*, GCCA introduces a low-cardinality global context without applying full attention over all image tokens.

### Geometry-Aware Mouse Synthesis Algorithm

The Geometry-Aware Mouse Synthesis Algorithm (GAMSA) was used to better represent severe overlap and occlusion during pose-model training. Starting from annotated single-mouse regions, the algorithm uses the bounding-box annotation as an anchor, binarizes the region to derive a local mouse mask, and extracts the corresponding image patch. The algorithm then transforms and composites the masked patch near another mouse in the image. The number of synthetic mice per image and the target overlap between synthetic instances are configurable, enriching the training distribution with contact configurations that occur infrequently in naturally sampled frames (Supplementary Note 2).

### BoT-SORT-based identity tracking

TMP detections were linked over time with a BoT-SORT-based multi-object tracker configured with appearance-based re-identification^23^. The tracker first predicts the current bounding-box state from the previous trajectory using a Kalman filter. The tracker associates high-confidence detections with predicted trajectories using spatial costs based on IoU and/or Mahalanobis distance. Unmatched trajectories are then compared with lower-confidence detections to recover animals whose detection confidence decreased during occlusion. A ReID encoder provides appearance embeddings, and cosine distance between embeddings supplements the motion-based association. Finally, trajectory management updates matched tracks, initializes new tracks, and removes tracks that remain unmatched for longer than the configured buffer. The conceptual association cost was written as

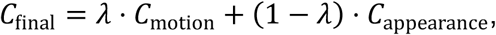

where *C_motion_*denotes the geometry-based association cost, Cappearance denotes the cosine distance between ReID embeddings, and *λ* denotes the balance coefficient.

### LLM-based automated behavioral-factor mining system System architecture and closed-loop mechanism

Factor mining was implemented as a closed-loop with four stages: hypothesis generation, sandboxed execution, statistical validation, and memory update. Each iteration generated a set of candidate behavioral factors, measured their predictive value, and returned the outcome to the next LLM request.

During hypothesis generation, the LLM received the current factor-library state, a summary of earlier successful and unsuccessful attempts, the complete feature catalog organized by semantic group, and the operational definition of each target behavior. The prompt highlighted underused feature combinations and behavior classes for which the current library contained few valid factors. Each response specified a mathematical transformation of a pose-derived temporal sequence and executable vectorized Python/NumPy code. The complete system and user prompt texts, together with the sources of every dynamic placeholder, are provided in Supplementary Note 3.

Generated code was evaluated in a restricted numerical sandbox. Import statements, file-system access, and network access were disabled, and only a whitelist of safe numerical operations was available. For *N* windows of length *L* and *D* input variables, *W* ∈ ℝ*^N^*^×*L*×*D*^, a batch-mode factor was required to return *f* ∈ ℝ*^N^*. Candidates had to produce finite values for at least 50% of windows. Factors with execution errors in more than 10% of windows were rejected.

Each surviving factor was evaluated independently with a one-versus-rest LightGBM classifier on a held-out validation set. A factor was retained if it reached AUC ≥ 0.65 for at least one behavior class. F1-Score was recorded for description but was not used as an acceptance gate. A factor meeting the threshold for several classes was registered for all qualifying classes (Extended Data Fig. 1a).

Accepted factors were stored with their class-specific AUC and F1-Score values in a structured library (Extended Data Fig. 1b). Rejected factors and their failure reasons were also recorded. Before the next request, the memory manager supplied the five most recent discovery rounds and the 15 highest-AUC factors verbatim; older history was summarized by target class to control context length. A diversity tracker identified overused and unexplored feature pairs and added that information to the prompt.

### LLM configuration and context management

The software exposes a common client interface for OpenAI-compatible chat-completion endpoints, Anthropic Claude, and GLM. Candidate factors were generated with temperature 0.7, a maximum output length of 16,384 tokens, and five requested factors per round. At least (n/2) floor of the requested factors was directed toward underrepresented behavior classes. The system prompt specified the normalized [0,1] input format, a vectorized batch convention, a prohibition on Python loops, and a parameterization convention in which tunable constants were declared as _P0, _P1, and so forth, along with default values and search ranges (Supplementary Note 3).

The user prompt contained the catalog of 42 variables, organized into seven semantic groups—subject skeleton, partner skeleton, subject motion, partner motion, subject tail, partner tail, and social variables—along with the formal class definitions, weak-class priorities, sequence-length setting, and hierarchical experience summary (Supplementary Note 4). The most recent five rounds and the top 15 valid factors were retained verbatim.

Earlier history was compressed by target class. The experience summary had a soft limit of 5,000 characters; when this budget was exceeded, the oldest detailed entries were converted to aggregate summaries rather than truncated. max_rounds was nominally 200, and discovery stopped after five consecutive rounds without a newly accepted factor.

### Statistical validation of candidate factors

Factor mining used a video-level training–validation holdout. LightGBM was fitted on designated training videos and evaluated on held-out validation videos; K-fold cross-validation was not used within the mining loop. The single-factor classifier used n_estimators = 200, max_depth = 4, learning_rate = 0.05, num_leaves = 31, random_state = 42 and n_jobs = 1. Early stopping monitored validation AUC with early_stopping_rounds = 20. For each class c, a separate one-versus-rest classifier distinguished class c from all other frames using only the candidate factor. Negative frames were randomly undersampled to achieve a 1:1 positive-to-negative ratio.

For class c, AUC was calculated from the one-versus-rest receiver-operating-characteristic curve, and F1-Score was calculated from precision and recall,

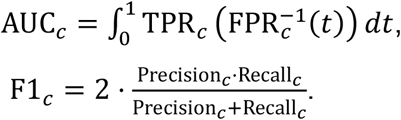

The AUC threshold of 0.65 was selected in preliminary analyses as a compromise between admitting many weak factors and restricting exploration when the factor space was sparsely sampled. F1-Score values were stored but were not used for factor acceptance.

### Correlation-based redundancy filtering

Different formulas can generate nearly identical factor time series. Redundancy was therefore quantified using Pearson correlation between factor outputs on the training data. After imputing missing values with the corresponding training-set mean, correlation between factors a and b was calculated as

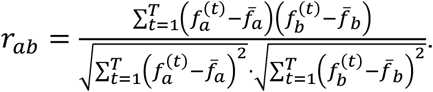

A constant factor was assigned zero correlation with all other factors. Pairs with |*r_ab_*| ≥ 0.95 were marked as redundant. Filtering was performed independently within the short-, intermediate- and long-range groups.

Factors were sorted by best AUC, and a factor was retained only if its absolute correlation with all previously retained factors in the same group was below 0.95. When two factors conflicted, the higher-AUC factor was retained. The resulting recommended factor list was used for downstream joint validation (Extended Data Fig. 1c).

### Factor mutation and parameter tuning

LLM-generated factors could expose tunable constants _P0, _P1, and so forth. Parameter templates were assigned according to parameter type: weight-like terms used a default of 1.0 and range [0.05,5.0]; threshold-like terms used a default of 0.5 and range [0.01,0.95]; scale-like terms used a default of 1.0 and range [0.1,5.0]; and exponents used a default of 1.0 and range [0.5,3.0].

Tuning proceeded in two stages. First, five mutation operators generated structural variants: AggregatorMutator replaced temporal aggregators such as mean, standard deviation, maximum, median or percentile; TemporalMutator replaced temporal summaries such as first-to-last difference, fitted trend or coefficient of variation; Cross4Mutator exchanged ratios, differences and products; NonlinearMutator inserted or replaced abs, square, square-root or log1p transformations; and ConstantMutator multiplied hard-coded constants by values in {0.5,0.75,0.9,1.1,1.25,1.5,2.0}. Each variant differed from its seed by one structural operation.

Second, declared parameters were searched by grid or random sampling, with at most 100 combinations per seed. One- or two-parameter factors used a fine grid of up to seven values per parameter; three- or four-parameter factors used four or five levels; and factors with five or more parameters used the center point, one-at-a-time variants, and random supplementation. Parameter values were materialized as numerical literals in the executable factor code before evaluation.

Each variant was evaluated with the same held-out LightGBM procedure used during discovery. A 40-tree pre-screen rejected a variant when its AUC was more than 0.05 below that of the seed. The remaining variants underwent the full 200-tree evaluation, and those that improved seed AUC were retained (Extended Data Fig. 1d). Factor-value calculation was parallelized using a thread pool, and LightGBM fitting was parallelized using a process pool. Compiled code was cached across variants sharing the same formula structure, and temporal-window matrices were cached by sequence length. On an eight-core machine, a seed with approximately 50 variants required about 30–60 s to tune.

### Factor integration and normalization

Validated factors were grouped by temporal receptive field *Lf*: short range, *Lf* ≤3 frames; intermediate range, 4 ≤ *Lf* ≤ 15 frames; and long range, *Lf* > 15 frames. At 30 frames s−1, these groups correspond approximately to ≤ 100 ms, 130–500 ms and > 500 ms. The short group captures instantaneous kinematics, the intermediate group captures brief action fragments, and the long group captures persistent states.

For a recording with *T* frames and *D* source variables, each factor *f* was evaluated on a centered temporal window ***W****_f_* ∈ ℝ*^T^*^×*L*^*^f^*^×*D*^. Boundary frames were replicated at both ends of the sequence. Executing the factor yielded a *T*-vector of values. Column-stacking all ***F*** factors produced a matrix ***F*** ∈ ℝ*^T^*^×*F*^. Because every factor was aligned to the central frame of its window, all factor vectors shared the same time axis. In the optional nan_boundary purity mode, factor values whose window crossed a manually annotated behavior boundary were set to NaN and imputed during normalization.

Factor columns were standardized with training-set statistics,

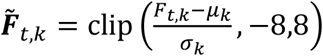

Missing values were imputed with the training-set column mean before scaling. The imputation value, mean, and scale of every factor were saved with the model and applied unchanged during inference. A temporal group typically contained approximately 600–3,000 factors, depending on discovery depth. The implementation optionally retained the top *K* factors ranked by average class-specific AUC; *K*=0 denoted no limit. Correlation filtering was applied before model fitting.

### Three-stage temporal behavior prediction Stage 1: temporal-group classifiers

Stage 1 trained an independent multiclass LightGBM model for each temporal group. For group g, the factor submatrix ***F***^(*g*)^ ∈ ℝ*^T^*^×*F*^*^g^* was standardized with training-set statistics and supplied to its group-specific classifier. Each classifier used n_estimators = 500, max_depth = 4, learning_rate = 0.05, num_leaves = 63, random_state = 42 and class_weight = "balanced". Early stopping monitored validation multiclass log loss with early_stopping_rounds = 50. When GPU execution was enabled, device="gpu" and deterministic=True were set. The three models generated posterior matrices P(short), P(medium), and P(long), each in ℝ*^T^*^×*C*^, and could be evaluated in parallel (Supplementary Note 2).

### Stage 2: meta-level fusion

The three group-level posterior matrices were concatenated to form the meta-representation

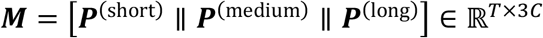

When residual_stage1 was enabled, standardized group-level factor matrices were concatenated with these posterior probabilitie,s allowing the meta-learner to access both factors and first-stage predictions. A LightGBM meta-classifier using the same nominal hyperparameters as the stage-one models was fitted on ***M*** and returned ***P***^(meta)^ ∈ ℝ*^T^*^×*C*^.

For the source implementation, stage-one models generated probabilities for the training and validation data; ***M****_train_*and ***M****_val_* were concatenated, and the meta-classifier was trained on Mtrain with early stopping on Mval.

### Stage 3: temporal-context modeling

Stage 3 incorporated explicit sequence context into P(meta). The software provided two alternative pathways. In the temporal-LightGBM pathway, a centred window of W=31 frames was used to calculate, for every class, the current probability, window mean, standard deviation, maximum, deviation from the window mean, and one-hot-encoded modal class. Additional scalar variables included the normalized number of class changes in the window, normalized current top-1 class identifier and probability entropy, yielding a base representation of 6*C*+3 variables. Optional multiscale windows of 7, 15, 31 and 63 frames could be concatenated. Optional extensions included first- and second-order probability differences, exponential moving averages with half-lives of 3, 10 and 30 frames, top-1–top-2 margin, Gini coefficient, lag-1 to lag-3 autocorrelation, short-versus-long trend differences, volatility ratios, peak differences, skewness and kurtosis. A second LightGBM model was fitted to the resulting temporal representation.

In the BiLSTM pathway, ***P***^(meta)^, optionally concatenated with the meta-level features when seq_use_extra_features was enabled, was supplied directly to a bidirectional recurrent network. The principal architecture contained four bidirectional LSTM layers, each with a hidden dimension of 1024 per direction and an inter-layer dropout of 0.3. The concatenated 2,048-dimensional hidden state was normalized, passed through a dropout layer with rate 0.3, and projected to *C* logits. The implementation also supported an optional two-layer GeLU multilayer perceptron in place of the linear projection and an optional output residual in which the stage-two probabilities were added to the logits before softmax (Supplementary Note 5). The Top-5 behavioral factor per behavior class were provided in Supplementary Note 6.

### Model training and computational environment TMP training

TMP was trained on 640 × 640-pixel RGB inputs, with a batch size of 16 and a nominal maximum of 300 epochs. The first three epochs were used for warm-up, during which momentum increased linearly from 0.8 to 0.937. The main optimizer was Adam with an initial learning rate of 0.01, β1 = 0.937, and weight decay of 5 × 10^−4^. Automatic mixed precision was enabled. Loss gains were 7.5 for bounding-box regression, 0.5 for classification, 12.0 for pose, and 3.0 for keypoint objectness. Augmentation varied hue by ±1.5%, saturation by ±70%, and brightness/value by ±40%. Translation magnitude was 0.1 and scale gain was 0.5. Mosaic augmentation was applied with probability 1.0 and disabled during the final 100 epochs. GAMSA-generated overlap examples were included in the training stream as described above (Extended Data Fig. 2a). The complete training configuration of the body and tail TMP models is provided in Supplementary Note 2.

### Three-stage predictor training

Stage-one group classifiers were fitted independently after Z-score standardization with training-set statistics. Validation data were transformed with the same statistics and used for early stopping. After stage one, group classifiers generated training and validation posterior probabilities, which were concatenated for meta-LightGBM training and early stopping.

The BiLSTM was trained on overlapping 1024-frame chunks with stride 512. Focal loss used γ=2.0 and class-specific α weights derived from inverse class frequency. AdamW used a learning rate of 5 × 10^−4^ and weight decay of 5 × 10^−4^. Gradient norm was clipped at 1.0. The learning rate was reduced on validation plateaus (ReduceLROnPlateau, factor 0.7, patience 8). Early stopping monitored validation balanced accuracy with a patience of 20 and a maximum of 200 epochs. Automatic mixed precision was used on compatible GPU hardware.

The three stages were optimized sequentially rather than end-to-end. This allowed the tree-based group, meta-classifiers, and the gradient-based BiLSTM to use separate objectives and optimization settings. Class imbalance was addressed by class_weight = "balanced" in LightGBM, inverse-frequency α weights in focal loss, and oversampling of chunks containing rare classes during BiLSTM training (Extended Data Fig. 2b). Random seeds, factor-computation parallelism, and the caching configuration used in the reported run are summarized in Supplementary Note 2.

### Hardware and acceleration

Experiments were run on a Linux workstation equipped with one NVIDIA RTX 5090 GPU with 32 GB memory, a 25-core Intel Xeon Platinum 8470Q CPU, and 90 GB system memory. TMP and the BiLSTM temporal model ran on the GPU. Factor evaluation, group and meta LightGBM models, and sequence decoding ran on the CPU. Distributed or multi-GPU training was not used, nor was TensorRT. TMP and BiLSTM training used automatic mixed precision. Group and meta LightGBM models were fitted on CPU because histogram construction for the shallow trees used here was faster across the available CPU cores than GPU offloading. Factor evaluation was parallelized across the same cores, and keypoint arrays were shared among worker processes through memory-mapped files.

TMP was trained at 640 × 640 pixels with a batch size of 16 on the order of 10³ s per epoch, with additional one-time dataset caching and AMP consistency checks during the first epoch. A complete run of approximately 30 epochs was reported to require about 8 h.

For the behavior predictor training, about 1.08 million training frames and 0.36 million validation frames across seven classes were used. The end-to-end run took approximately 1 h 42 min, including loading, three group classifiers, the meta-model, temporal model, validation, figure generation, and calibration. In this run, the group and meta LightGBM stages took about 5 min because the standardized factor matrices were loaded from cache; the BiLSTM trained for 104 epochs with early stopping in about 41 min (∼24 s per epoch); and the post-hoc calibration/decoder grid search took about 49 min.

We used ChatGPT 5.5 for behavioral factor mining for training SABER. LLM-driven mining was dominated by API latency. The source reports 55–80 s per generation request, with sandbox execution and LightGBM validation completed locally within seconds. One hundred mining rounds per temporal group required approximately 2 h, and the reported complete library contained 2948 validated factors accumulated across temporal groups.

Once keypoints were available, the temporal stage processed 360,008 frames in approximately 11 s, corresponding to approximately 3.3 × 10⁴ frames s−1 for model forward evaluation. Sliding-window feature assembly required approximately 4.1 s. Group and meta gradient-boosting stages added only a few seconds of GPU time, making TMP pose estimation the dominant per-frame cost for full-video analysis.

### End-to-end inference and post-processing Online pose estimation and identity tracking

Each frame was resized with letterboxing to 640 × 640 pixels and normalized to (image/255-μ) / σ. Online inference used a batch size of 1. TMP returned keypoint heatmaps, bounding boxes, and class scores. Coarse keypoint coordinates were obtained from the heatmap argmax and multiplied by the output stride of 32. A local center-of-mass calculation around the peak provided subpixel refinement. Keypoints with confidence ≤ 0.3 were discarded. Decoded detections were passed to the BoT-SORT-based tracker for Kalman prediction, cascaded association using IoU/Mahalanobis geometry and ReID appearance features, and track update. The per-frame output was represented as an [*N*, *K*, 4] array containing *x*, *y*, confidence, and track_id.

For contact-rich recordings, track_buffer was set to 300 frames, with_reid=True, appearance_thresh=0.65 and fuse_score=True. A high new-track threshold was used to suppress false trajectories initiated from background detections (Extended Data Fig. 3a). The same inference code supported batched offline evaluation.

### Secondary behavioral features

Identity-linked pose coordinates were transformed into 42 frame-level variables organized into seven semantic groups (Supplementary Note 3). All variables were aligned to the native 30-frame s^−1^ time axis and mapped to [0,1] using feature-wise min–max parameters estimated from the training data. The flat_attributes index exposed each variable by name to the factor sandbox.

For landmark k at frame t, centered five-point velocities were calculated as

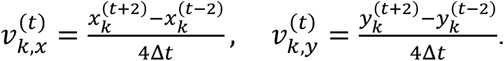

Body-center speed was the Euclidean magnitude of body-center velocity. Acceleration was the magnitude of the temporal derivative of velocity, and movement direction was calculated with 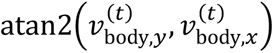. The same velocity, speed, and acceleration variables were calculated for the partner.

Within-animal structural variables included distances between anatomically related landmark pairs—snout to head_center, head center to body center and body center to tail base—body orientation from body center to head center, snout-to-tail-base length, and body compactness, defined as the mean pairwise distance among the four keypoints (snout, head center, body center and tail base); compactness is small when the body is curled up.

#### Corresponding structural variables were calculated for the partner

Social variables included the Euclidean distance between center of bounding boxes; subject-facing-partner and partner-facing-subject scores defined by the cosine between the animal heading vector and the vector to the other animal; mutual facing, calculated as the product of the two facing scores; approach speed, defined as the projection of the relative velocity onto the self-to-other direction (approximately the negative derivative of inter-animal distance); relative speed; absolute angular separation of body orientation; and body-axis alignment, cos(Δ*φ*). The 42 variables formed ***X*** ∈ [0,1]*^T^*^×42^, which was the input to factor computation. Feature-use frequencies in the Results were calculated directly from occurrences of these variables in the final factor library.

### Cascaded inference of the three-stage predictor

At inference, the factor library, normalizers, and trained models were loaded from the run bundle. Each factor was evaluated on its native centered window, producing a *T*-vector of values. Factor vectors were assembled into short-, intermediate- and long-range matrices and standardized with the stored training means and scales. The three group-specific LightGBM models returned ***P***^(short)^, ***P***^(*medium*)^ and ***P***^(*long*)^. These posterior matrices were concatenated, with optional residual factor features, and processed by the meta-LightGBM to produce ***P***^(meta)^. In the BiLSTM pathway, ***P***^(meta)^., optionally concatenated with meta-features, was divided into overlapping 1,024-frame chunks with stride 256. Predictions from overlapping positions were averaged. In the temporal-LightGBM pathway, sliding-window probability features were calculated and processed by the second-level LightGBM.

The final frame-level class was 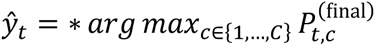.

The output CSV contained frame index, timestamp in seconds, predicted class identifier, class name, and full class-probability vector.

### Probability calibration and rule-based correction

Optional post-processing calibrated the final probabilities before applying label-level corrections. Global temperature scaling transformed logits z as

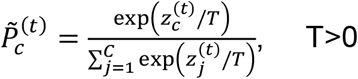

T was selected on the validation data by minimizing the negative log-likelihood. Class-specific calibration added a bias vector b in logit space. When both BiLSTM and meta-LightGBM probabilities were retained, per-class convex mixing was used,

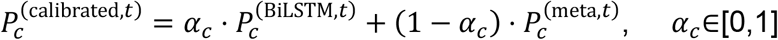

Calibration parameters were selected by coarse-to-fine scalar α search, class-specific α refinement in steps of 0.02, six rounds of class-bias search, 500 joint random-perturbation iterations, and temperature search over [0.75,1.35] in steps of 0.01. The combination maximizing validation accuracy was retained.

The rule-based corrector then operated on discrete labels. Flicker removal targeted segments shorter than max_flicker = 3 frames when the BiLSTM probability of the predicted class was below flicker_prob_threshold = 0.05; high-confidence frames with a top-1 versus top-2 margin > 0.3 were exempt. Boundary refinement examined a 31-frame window around each transition and delayed a class change while the old-class probability remained above 0.20. Segments shorter than min_consecutive = 5 frames were assigned to the adjacent class with the larger mean probability over the short segment. Calibration was applied before label correction (Extended Data Fig. 3b). In addition to the core functionalities described above, this software also supports the analysis of mouse activity heatmaps (Extended Data Fig. 3c). Users can define custom polygonal regions of interest (ROIs) to quantify entry and exit counts for a specified region and the dwell time within that region (Extended Data Fig. 3d).

### Evaluation metrics and benchmarking

#### Pose-estimation metrics

Pose accuracy was evaluated with Object Keypoint Similarity (OKS)-based precision–recall metrics. For visible keypoint i,

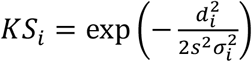

where di is the Euclidean distance between the prediction and manual annotation, s is the object-scale term, and *σ_i_* is a landmark-specific normalization constant estimated from annotation variability. Instance-level OKS was the visibility-weighted mean,

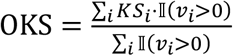

At OKS threshold τ, a matched prediction was counted as a true positive; an unmatched prediction was a false positive, and an unmatched ground-truth instance was a false negative. Precision, recall, and F1-Score

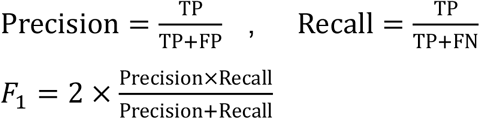

Average precision at threshold τ was the area under the precision–recall curve. Mean average precision was calculated across ten OKS thresholds from 0.50 to 0.95 in increments of 0.05,

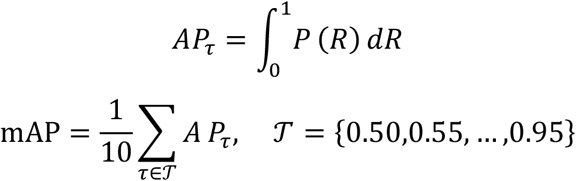

Predictions and ground-truth instances were greedily matched by selecting the highest-OKS unmatched prediction for each ground-truth instance and accepting the pair when OKS ≥ *τ*.

### Tracking-error metrics

Tracking was evaluated with three application-specific error counts. An identity switch (IDSW) occurred when the track identifier assigned to a true individual changed inconsistently between consecutive frames. Target Loss occurred when an animal present in the scene did not receive an accepted pose estimate, such as detections falling below the configured confidence threshold, causing its track to be interrupted. A keypoint switch (KPSW) occurred when one or more landmarks in an otherwise complete predicted skeleton were assigned to another animal. These events were manually or algorithmically compared with identity-annotated ground truth in the evaluated recordings.

### Behavior-classification metrics

Frame-level predictions were compared with manual labels using scikit-learn v1.3 or later. Accuracy was the fraction of correctly labeled frames,

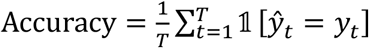

Balanced Acuracy was the unweighted mean of per-class Recall,

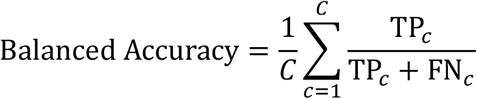

Macro F1-Score was the unweighted mean of class-specific F1-Score values, whereas weighted F1-Score weighted each class by its support. Multiclass AUC was calculated in a one-versus-rest framework and reported as either macro AUC or support-weighted AUC. Raw and row-normalized confusion matrices were generated. All validation frames, including frames close to manually annotated behavior transitions, were retained in the reported metrics.

The implementation used accuracy_score, balanced_accuracy_score, f1_score with average = "macro" or "weighted", roc_auc_score with multi_class = "ovr" and average = "macro" or "weighted", confusion_matrix and classification_report. Per-class precision, recall, F1, support, and AUC were reported with aggregate metrics.

### Inference throughput and latency

All timing measurements were performed on the same single-GPU workstation. Throughput was defined as the number of frames processed per second under the native batch configuration, whereas latency was the time required to process one frame at batch size 1. GPU stages were warmed up with several 640 × 640-pixel dummy inputs before timing to exclude CUDA context creation, cuDNN autotuning, and lazy memory allocation. Throughput was measured over thousands of frames rather than a small number of batches.

Timing scope was recorded explicitly. End-to-end timing included camera acquisition or video decoding, preprocessing, model forward evaluation, tracking, and post-processing. Forward-only timing included model computation and, where specified, feature assembly but excluded image acquisition and decoding. PyTorch models ran in FP16 during the real-time path.

Two representative live-camera sessions contained 4,604 and 2,485 frames and were processed at 30.0 frames s^−1^, requiring 153.5 and 82.8 s, respectively. The corresponding frame budget was 33.3 ms. Camera acquisition, in-memory JPEG transfer, pose estimation, identity association, rendering, and periodic checkpointing were included. Behavior classification ran in a separate process, so the acquisition loop remained capture-limited at the camera frame rate.

Offline pose inference used batch size 16 at 640 × 640 pixels. Body and tail pose models could be dispatched to concurrent CUDA streams. On 360,008 pre-extracted frames, the temporal model required approximately 11 s for forward evaluation (approximately 3.3 × 10⁴ frames s^−1^), and sliding-window feature assembly required approximately 4.1 s. A separate profiling utility measured in-memory JPEG transfer, disk round trips, overlay rendering, and interprocess buffering over 300 frames.

### Benchmarking SABER against multiple pose-tracking and behavior-classification frameworks

To benchmark SABER against established pose-estimation, tracking, and behavior-classification approaches, we implemented three pose-tracking baselines—DeepLabCut (DLC, https://github.com/DeepLabCut/DeepLabCut,v2.3.11), SLEAP (https://github.com/talmolab/sleap, v1.4.1), and AlphaTracker (https://github.com/MVIG-SJTU/AlphaTracker, v0.1.0)—and three behavior-analysis baselines—A-SOiD (https://github.com/YttriLab/A-SoID, v0.3.1.3), B-SoiD (https://github.com/YttriLab/B-S0ID, v2.0), and a support vector machine (SVM)-based decoder (Inter-brain) adapted from the behavioral analysis described in a recent study^3^. DeepLabCut and SLEAP are established multi-animal pose-tracking frameworks, whereas AlphaTracker uses a top-down multi-animal tracking workflow; A-SOiD and B-SOiD provide supervised/active-learning and unsupervised approaches, respectively, for converting pose measurements into behavioral categories.

For the pose-estimation comparison, all methods were trained from the same labeled training images with the same resolution and keypoint annotations used for SABER. The framework-specific recommended configuration was retained, and no additional task-specific hyperparameter search was performed for the benchmark.

DLC used a ResNet-50 backbone. The model was optimized with Adam at an initial learning rate of 5 × 10^−4^, followed by a multistep learning-rate decay schedule, for 78,000 iterations. Training augmentation included random rotations within ±25° and image scaling. DLC was evaluated using 640 × 640-pixel input, and predictions were linked across frames, with the tracker set to the default in the benchmark configuration.

SLEAP used a U-Net architecture with 640 × 640-pixel inputs. Models were trained for 200 epochs with Adam at a learning rate of 1 × 10^−4^. Augmentation comprised random rotations within ±15° and horizontal flipping. Predicted poses were associated across frames using flow-shift tracking.

AlphaTracker was implemented as a two-stage pipeline comprising a YOLO detector and a single-person pose-estimation (SPPE) network. The detector was trained for 60,000 iterations at a learning rate of 5 × 10^−4^, and the SPPE network was trained for 10 epochs at a learning rate of 1 × 10^−4^. The detector retained the native YOLO input dimensions. Consecutive-frame detections were associated using intersection-over-union (IoU) matching.

Behavioral prediction was compared with A-SoiD, B-SOiD, and Inter-brain. All comparator models received the same underlying pose trajectories as SABER. A-SOiD used 100-ms feature bins. Input features were standardized with StandardScaler, and classification was performed with a random forest containing 200 trees, the Gini impurity criterion, and class_weight = "balanced_subsample". B-SOiD used the same 100-ms feature bins and StandardScaler preprocessing. Features were embedded with UMAP using 30 nearest neighbors (n_neighbors = 30), min_dist = 0 and two output dimensions, after which clusters were identified with HDBSCAN using min_samples = 1.

Because code for Inter-brain was not released in the cited study, we reconstructed the classifier from the published methodological description and adapted the feature representation to the landmarks available in our dataset. For each mouse, we constructed an 87-dimensional pose-derived feature vector. This differed from the 75-dimensional representation in the reference analysis because the anatomical keypoints available in the two datasets were not identical. Features were standardized with StandardScaler. Binary classification used LinearSVC with C = 1.0. Positive and negative samples were randomly downsampled to a 1:1 ratio; the analysis required a minimum of 50 frames, and 100 label permutations were used for permutation-based evaluation.

### Chronic social defeat stress model

The chronic social defeat stress (CSDS) mouse model was used as previously described^31^. In brief, mice were acclimated to single-housing for 1 week prior to the chronic social defeat stress (CSDS) procedure. Over the next 10 days, each animal was subjected to a daily 5–10 min encounter with a novel CD1 aggressor, which was replaced daily. Immediately after the defeat session, the experimental mouse was left in the aggressor’s home cage but kept physically apart by a perforated, translucent plastic partition. Control animals were housed in pairs with an identical divider, never exposed to CD1 mice, and moved each day to a cage containing an unfamiliar, non-aggressive mouse. Social interaction was assessed 24 h following the final defeat episode.

### Behavioral tests

The social interaction test (SIT) comprised two 150-s phases. In the first phase, each mouse was placed into a novel open-field arena (42 × 42 × 42 cm) that contained an empty wire-mesh cage (10 × 6.5 × 42 cm) positioned centrally above the floor; this condition was designated as the “no-target” session. After a 30-s interval back in their home cage, the mice returned to the same arena for the second phase, but the wire enclosure now held an unfamiliar aggressive CD-1 mouse (the “target” condition). The area surrounding the enclosure within an 8-cm radius, forming a 14 × 24 cm rectangle, was defined as the interaction zone. Behavior was recorded using Ethovision XT 14.0 (Noldus), and a social interaction ratio was calculated as the duration of interaction with the target divided by the duration without the target. Mice with SIR < 1 were considered Sus.

The sucrose preference test (SPT) was carried out using a two-bottle choice paradigm over two consecutive days, with food available without restriction. Each mouse was singly housed and given simultaneous access to a 2% sucrose solution and plain water; bottle positions were swapped after the first 24 h. Fluid intake from each bottle was measured at the end of every 24-h period. Sucrose preference was expressed as the percentage of total fluid consumed from the sucrose bottle, calculated as [sucrose intake / (sucrose intake + water intake)] × 100%.

For the tail suspension test (TST), mice were suspended individually by affixing adhesive tape to the tail and attaching it to a hook fixed to the ceiling of a rectangular chamber (70 × 30 × 30 cm). The 6-min session was recorded using Ethovision XT 14.0. Immobility was scored when the animal failed to exhibit any active movement during the observation period.

### Stereotaxic Surgery, viral Injection, and implantation

Mice were anesthetized with inhalation of 2-3% isoflurane and secured in a stereotaxic frame (RWD, China). The skull was exposed, and bregma and lambda were leveled horizontally before stereotaxic injection. For viral injection, a craniotomy was made over the left mPFC. Viral vectors were delivered into the mPFC using a glass capillary that connects to a microinjector (Nanoliter 2020, World Precision Instruments) at a rate of 20 nl/min. Injection coordinates were relative to bregma: ML: −0.30 mm; AP: +1.98 mm; DV: −2.00 mm. The capillary was slowly withdrawn 10 minutes after injection. Incision sites were sutured and treated with erythromycin ointment. Following surgery, mice were placed on a heating pad until they regained consciousness, then returned to their home cages for 3 weeks before subsequent experiments. The following AAV vectors were used: AAV2/9-mCaMKIIα-jGCaMP8s-WPRE-pA (1.0 × 10^13^ v.g./ml, 0.2 µl, Taitool). The vector was aliquoted and stored at −80°C until use.

For miniscope imaging, a GRIN lens (0.5 mm diameter, 3.7 mm length) was implanted 0.10 mm above the injection site and secured with cyanoacrylate adhesive and dental cement. After 2–3 weeks of recovery, a baseplate was positioned and fixed once individual cells were in focus.

### Miniscope calcium imaging and analysis

Calcium imaging was performed with a UCLA Miniscope V4 system (Open Ephys; URL) operating at 30 fps. Mice underwent a 30-min habituation in a white plastic open-field chamber immediately before a 10-min recording session on the day of the experiment. Motion correction of the captured videos was accomplished via NoRMCorre^36^, and the extraction of single-neuron components was carried out using CNMF-E^37^, with parameters deliberately set to a conservative range so that roughly 20–50 components were accepted per imaging field. The accepted components and their CNMF-E-derived time series (i.e., C + YrA) were used for all downstream calculations, which included dynamic ΔF/F estimation and subsequent z-score normalization. All plots display z-scored ΔF/F.

### Statistical analysis

Statistical analyses were performed using GraphPad Prism. Normality was assessed with the Shapiro–Wilk test. The unpaired student t-test and the Mann-Whitney test were used to compare behavioral results between Ctrl and SUS mice when data passed or failed normality test, respectively. Linear discriminant analysis was performed using sklearn function-LinearDiscriminantAnalysis with zero-centering and unit-variance scaling of data prior to projection. Inputs to the LDA were the seven behavioral parameters from each biological replicate. For two-group classification, the analysis yields a single discriminant axis (LD1) that maximizes the ratio of between-group to within-group variance (Fisher’s criterion). LD2 was constructed as the orthogonal distance of each sample to its own group centroid in the standardized feature space, representing within-group dispersion for visualization. Ninety-five percent confidence ellipses were computed based on the bivariate chi-squared distribution using the group-specific covariance matrix and its eigen-decomposition. Sample sizes are indicated in the Results and figure legends. Data are presented as mean ± SEM, with significance at *p* < 0.05.

## Data availability

Data access is unrestricted. The full dataset analyzed in this study, comprising per-video TMP keypoints, per-mouse behavior annotations, and the dataset configuration file, can be downloaded from ScienceDB at https://www.scidb.cn/s/UrAjMr. The TMP pose checkpoints and the trained three-stage behavior-prediction model bundle, along with all other model weights, are likewise available from ScienceDB at https://www.scidb.cn/s/RjQRFz. Source data are included with this paper.

## Code availability

All associated resources can be found on our GitHub page at https://kidous2333.github.io/SABER/. SABER’s Python source code is available at https://github.com/kidous2333/SABER. This code is distributed under the ’ACADEMIC OR NON-PROFIT ORGANIZATION NONCOMMERCIAL RESEARCH USE ONLY’ license (see the LICENSE file in the repository). A user-friendly desktop GUI is also provided in the repository; it replicates all command-line functions, including pose-model training, factor mining, factor engineering, and behavior prediction.

## Acknowledgments

We thank the lab members for helpful discussions and suggestions. This work was supported in part by STI2030-Major Projects (2026ZD0223300), Noncommunicable Chronic Diseases-National Science and Technology Major Project (2023ZD0507100), the National Natural Science Foundation of China (grant no. 82271555, 82322024, and 82671967), and the Guangdong Major Project of Basic Research (2026B0303000010).

## Author Contributions

X.-D.S., J.Z., and C.P. designed the study; J.Z. constructed the SABER system; J.W., W.-N.Z., and H.-M.X. recorded videos of mice’s social behaviors, generated the chronic social defeat stress model, and conducted miniscope calcium imaging; S.-Y.Z. annotated keypoints and behavior; T.-Q.C. and Y.-W.L. annotated keypoints; Z.J. helped with the ideas of LLM-based behavioral factor mining; Y.T. assisted in comparison among SABER, SLEAP, DLC, and AlphaTracker; C.P. and J.Z. organized figures; X.-D.S. and J.Z. wrote the manuscript.

## Resource Availability

The corresponding author, Xiang-Dong Sun, will address further information requests and resource requests.

## Declaration of generative AI and AI-assisted technologies in the manuscript preparation process

In the study, we used ChatGPT 5.5 to assist in behavioral factor discovery. In addition, Claude Code (Anthropic) with the DeepSeek-V4 model was used solely to organize project parameters and compile the draft Supplementary Information, the content of which was prepared and verified by the authors. In drafting the manuscript, we used ChatGPT 5.6 to polish the language and improve the paper’s readability. All proposed edits from the AI were subsequently reviewed and adjusted by the authors, who bear full responsibility for the submitted version.

**Extended Data Fig. 1 | GUI-integrated factor mining and experimentation system. a**, Factor mining interface showing factor information and class-wise AUC values. **b**, Factor details page documenting comprehensive information for each factor, including class-wise AUC values, factor hypotheses, factor formulas, and calculation code. **c**, Correlation-based redundancy filtering, calculating pairwise correlations among factors and recording the factors retained after screening. **d**, Factor mutation and parameter tuning showing the AUC improvements of successfully fine-tuned factors and the corresponding fine-tuning methods.

**Extended Data Fig. 2 | Training procedures of the TMP model and the three-stage predictor**. **a**, TMP model training, showing the decreasing curves of each loss term per epoch, the mAP metric, and changes in the learning rate. **b**, Training interface of the three-stage predictor, which shows the model training performance and individual metrics at each stage, enabling visual analysis for researchers.

**Extended Data Fig. 3 | End-to-End Model Inference. a**, Global trajectory map. The movement trajectories of mice are recorded following the start of inference. **b**, End-to-end model inference. The residual trajectories over a selected time period, mice points, and behavioral raster plots for all mice according to the settings was displayed. **c**, Heatmap visualization of mice activity. **d**, Custom region analysis. Users can draw polygonal regions on the GUI to analyze the number of entries into and exits from the regions and the dwell time.

