## Extended Figures for "From pose to behavior: SABER integrates identity-resolved multi-animal pose tracking with large-language-model-based behavioral factor discovery"

### Extended Data Fig. 1

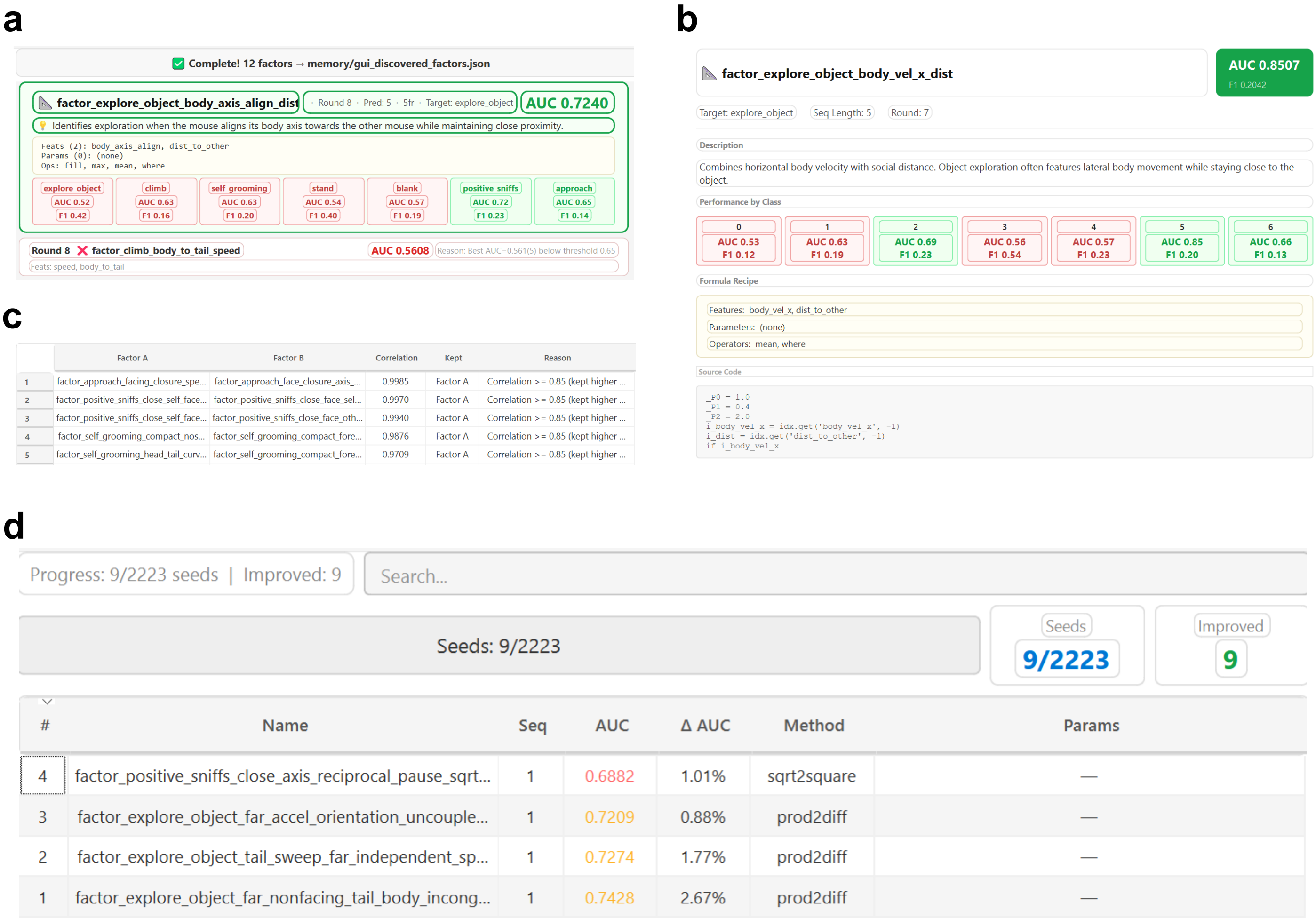

### Extended Data Fig. 2

**a**

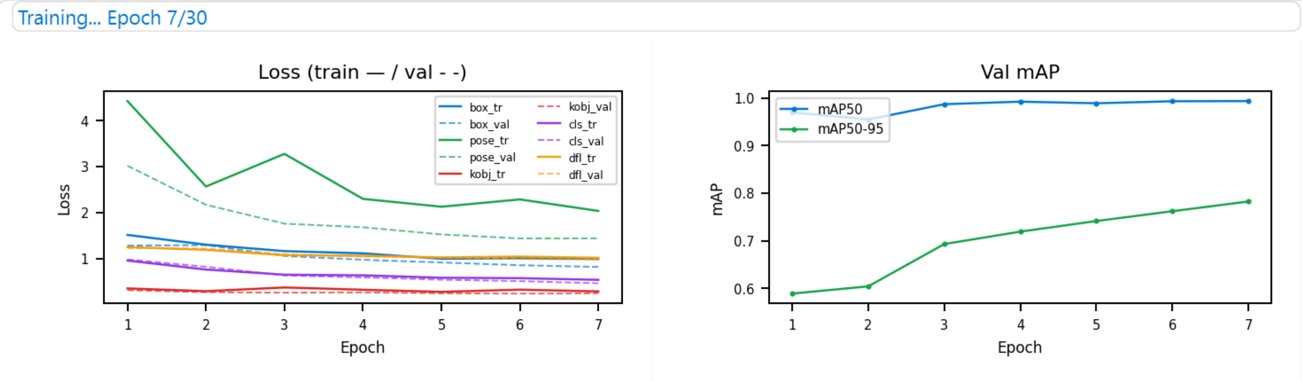


| | Epoch | box | pose | kobj | cls | dfl | mAP50 | mAP50-95 | LR |
|---|---|---|---|---|---|---|---|---|---|
| 2 | 2/30 | 1.2989 | 2.5645 | 0.2875 | 0.7583 | 1.1869 | 0.9548 | 0.6040 | 1.29e-03 |
| 3 | 3/30 | 1.1591 | 3.2742 | 0.3685 | 0.6466 | 1.0712 | 0.9870 | 0.6930 | 1.87e-03 |
| 4 | 4/30 | 1.1106 | 2.2956 | 0.3186 | 0.6325 | 1.0579 | 0.9923 | 0.7191 | 1.80e-03 |
| 5 | 5/30 | 0.9925 | 2.1244 | 0.2729 | 0.5802 | 1.0217 | 0.9889 | 0.7414 | 1.74e-03 |
| 6 | 6/30 | 1.0055 | 2.2837 | 0.3205 | 0.5706 | 1.0400 | 0.9931 | 0.7620 | 1.67e-03 |

**b**

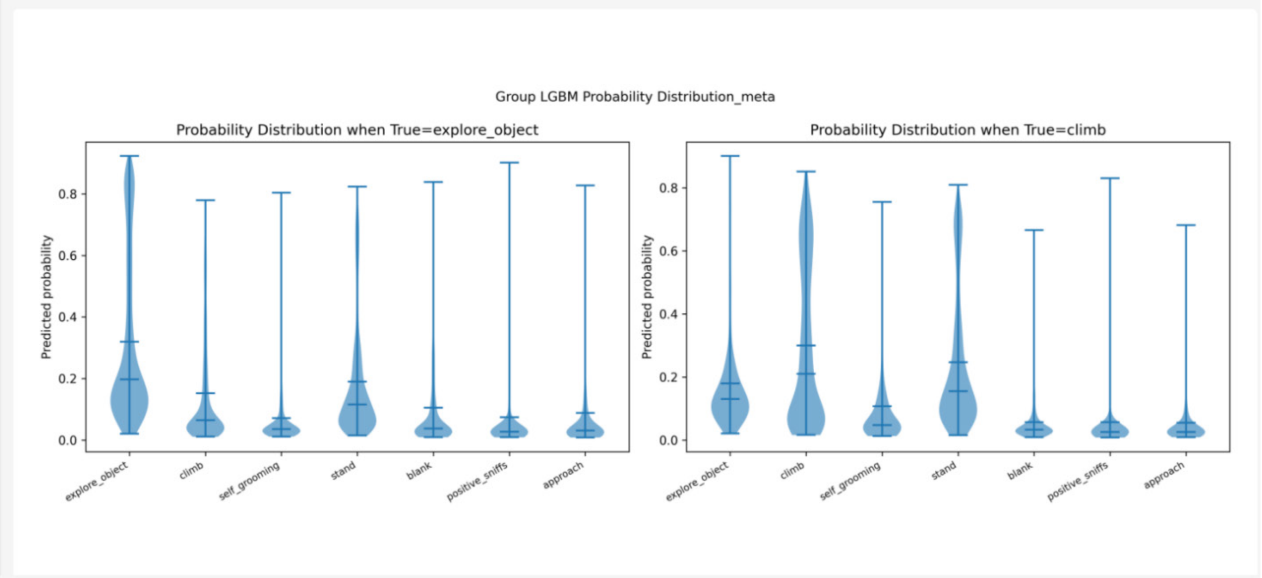


| | Stage | Accuracy | macro-F1 | macro-AUC | weighted-F1 | bal.Acc |
|---|---|---|---|---|---|---|
| 1 | Group (short) | 0.3454 | 0.3363 | 0.7676 | 0.3625 | 0.4459 |
| 2 | Group (medium) | 0.3328 | 0.3184 | 0.7791 | 0.3428 | 0.4296 |
| 3 | Group (long) | 0.3211 | 0.2879 | 0.7324 | 0.3571 | 0.3555 |
| 4 | Meta-LGBM | 0.4084 | 0.3682 | 0.7928 | 0.4298 | 0.4392 |

Image Groups

Visualizations/05 Meta

Visualizations/01 Data

3/4

◂ Prev Next ▸

### Extended Data Fig. 3

a

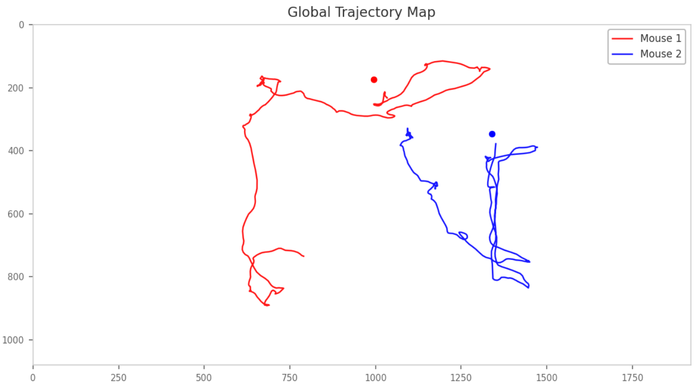


b

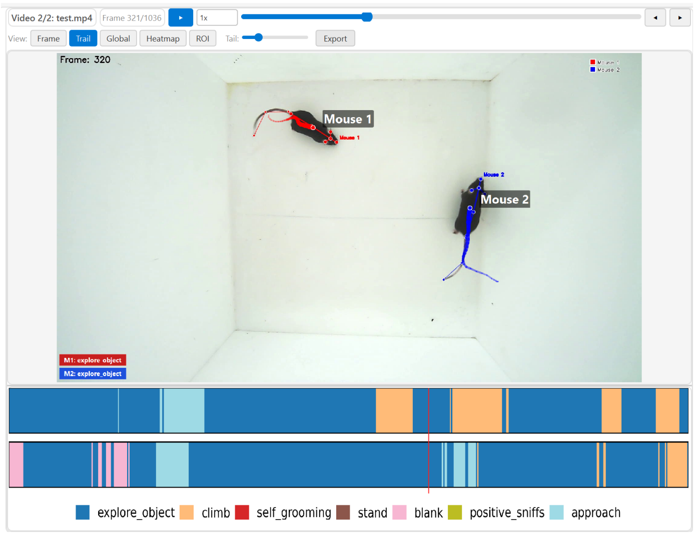


c

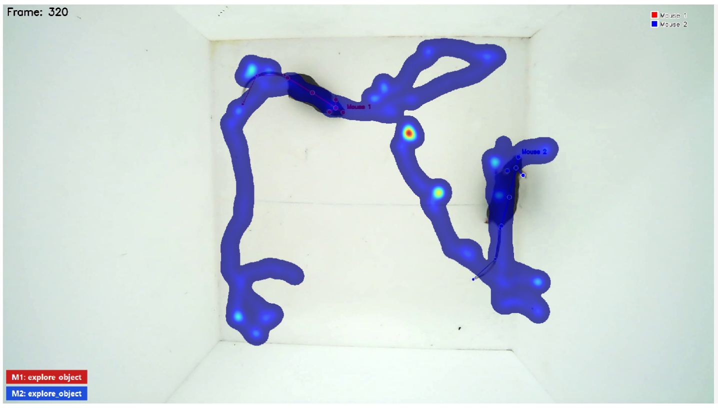


d

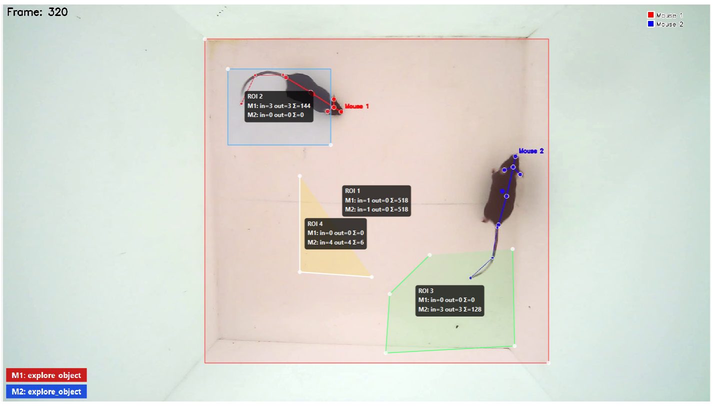
