## Supplementary Note 1 for "From pose to behavior: SABER integrates identity-resolved multi-animal pose tracking with large-language-model-based behavioral factor discovery"

This note describes the mouse dyadic behavior dataset, the feature-extraction pipeline and the data-preprocessing steps, and the composition and frame-level class distribution of the dataset. This note contains Supplementary Tables 1–2.

---

### Content

**Supplementary Table 1.** Composition of the behavioral dataset.

**Supplementary Table 2.** Frame-level class distribution of the behavioral dataset (training and validation sets).

---

### Dataset

**Supplementary Table 1. Composition of the behavioral dataset.**

| Item | Value |
|---|---|
| Subjects | 10 mouse pairs (subject + companion, 2 mice per video) |
| Training clips | 30 clips (10 pairs × 3 sessions) |
| Validation clips | 10 clips |
| Pose source | TMP pose estimation (mouse keypoints and tail keypoints, per-mouse tracking) |
| Behavior annotations | frame-level behavior labels for both mice |
| Total frames | 1,439,624 (train 1,079,616 / validation 360,008) |
| Frame rate / format | per-frame features normalized to [0,1] |

**Class distribution of the behavioral dataset (frames)**

**Supplementary Table 2. Frame-level class distribution of the behavioral dataset (training and validation sets).**

| Class | Frames (train) | Share (train) | Frames (val) | Share (val) | Frames (total) | Share (total) |
|---|---|---|---|---|---|---|
| Explore | 380,067 | 35.2% | 101,827 | 28.3% | 481,894 | 33.5% |
| Stand | 316,006 | 29.3% | 89,285 | 24.8% | 405,291 | 28.2% |
| Blank | 123,361 | 11.4% | 70,725 | 19.6% | 194,086 | 13.5% |
| Self-grooming | 108,146 | 10.0% | 54,777 | 15.2% | 162,923 | 11.3% |

| Class | Frames (train) | Share (train) | Frames (val) | Share (val) | Frames (total) | Share (total) |
|---|---|---|---|---|---|---|
| Sniffing | 53,352 | 4.9% | 19,703 | 5.5% | 73,055 | 5.1% |
| Climb | 66,413 | 6.2% | 13,851 | 3.8% | 80,264 | 5.6% |
| Approach | 32,271 | 3.0% | 9,840 | 2.7% | 42,111 | 2.9% |
| Total | 1,079,616 | 100% | 360,008 | 100% | 1,439,624 | 100% |

The dataset is strongly imbalanced (the largest class has ~10× the frames of the smallest), which motivates the class-weighted training strategies described below.

**Label map**

Explore: 0, Climb: 1, Self-grooming: 2, Stand: 3, Blank: 4, Sniffing: 5, Approach: 6 (7-class label space).

**Feature extraction and preprocessing**

- 42 engineered features per frame (see Supplementary Note 4), grouped into Skeleton / Motion / Tail / Social for both mice.
- Normalization: percentile clipping [0.1%, 99.9%] followed by min-max scaling to [0,1], fitted on the training set.
- Missing keypoints: bidirectional imputation (short-gap interpolation, max gap 10 frames).
- Boundary handling: purity_mode = nan_boundary (behavior-transition frames near annotation boundaries are masked), boundary_margin = 10, purity_threshold = 1.0.
- Training augmentation: horizontal flip, temporal jitter (±2 frames), rotation jitter (±5°).

---
