## Supplementary Note 2 for "From pose to behavior: SABER integrates identity-resolved multi-animal pose tracking with large-language-model-based behavioral factor discovery"

This note describes the training configuration of the TMP, which is trained as a single-class keypoint detector at 640 × 640 pixels, with one model for the mouse body (six keypoints: snout, head_center, body_center, tail_base, left_ear, right_ear) and a second model for the mouse tail (three keypoints: tail_start, tail_middle, and tail_end), as described in Methods ("Keypoint definition and pose annotation"). It accompanies the Methods section "Pose-estimation and identity-tracking models". This note contains Supplementary Tables 3–6.

---

### Content

**Supplementary Table 3.** TMP model variants and training data.
**Supplementary Table 4.** General training configuration of the TMP models.
**Supplementary Table 5.** Optimizer, learning-rate schedule, and loss configuration.
**Supplementary Table 6.** Data augmentation used during training.

---

### Overview

The two model variants and their training data are summarized in Supplementary Table 3. The same source videos were annotated for both the body and the tail keypoints; because the two keypoint groups differ in number, they are used to train two separate single-class models, each using only its own keypoint annotations. Both models were trained at 640 × 640-pixel input resolution with a batch size of 16. Synthetic overlap examples generated by the Geometry-Aware Mouse Synthesis Algorithm (GAMSA; see Methods "Geometry-Aware Mouse Synthesis Algorithm") were added to the training stream. Training values below are taken from the saved training configurations and training logs of the released models (TMP_mouse.pt, TMP_tail.pt).

---

**Supplementary Table 3. TMP model variants and training data.**

| Item | Body model | Tail model |
|---|---|---|
| Detected keypoints | six landmarks per mouse: snout, head_center, body_center, tail_base, left_ear, right_ear | tail keypoints: tail_start, tail_middle, tail_end |
| Class configuration | single class (mouse) | single class (mouse tail) |

| Item | Body model | Tail model |
|---|---|---|
| Training data | one shared annotated dataset for both models — frames from the two-mouse social, aggressive, and mating recordings (3,000 frames in total, 3:1 training-to-validation split; see Methods "Keypoint definition and pose annotation"), using only the body keypoint annotations | the same shared annotated dataset, using only the tail keypoint annotations |

**Supplementary Table 4. General training configuration of the TMP models.**

| Item | Value |
|---|---|
| Input resolution | 640 × 640 pixels |
| Batch size | 16 |
| Nominal epochs | 300, with early stopping (patience 10 epochs) |
| Class configuration | single class (mouse body; mouse tail) |
| Mixed precision | automatic mixed precision (AMP) enabled |
| Determinism | deterministic operations enabled, random seed 0 |
| Validation protocol | internal validation on the held-out split after every epoch; IoU threshold 0.7, up to 300 detections |

**Supplementary Table 5. Optimizer, learning-rate schedule, and loss configuration.**

| Item | Value |
|---|---|
| Optimizer | automatic selection (AdamW); initial learning rate 0.01; final learning-rate factor 0.01 |
| Momentum / β1 | 0.937 |
| Weight decay | $5 \times 10^{-4}$ |
| Learning-rate schedule | constant (cosine schedule disabled) |
| Warm-up | 3 epochs; momentum increased |

| Item | Value |
|---|---|
| | linearly from 0.8 to 0.937; bias learning rate 0.0 (body model) / 0.1 (tail model) |
| Loss gains | bounding-box regression 7.5, class 0.5, distribution focal loss 1.5, keypoint (pose) 12.0, keypoint objectness 3.0 |

**Supplementary Table 6. Data augmentation used during training.**

| Augmentation | Value |
|---|---|
| Hue / saturation / brightness (value) | ±1.5% / ±70% / ±40% |
| Translation | 0.1 (fraction of image) |
| Scale | 0.5 gain |
| Horizontal flip | probability 0.5 |
| Mosaic | probability 1.0, disabled during the last 100 epochs |
| Mixup / cutmix / copy-paste | 0 (disabled) |
| Random erasing | 0.4 |
| Automatic augmentation policy | rand-augment |
| Rotation / shear / perspective | 0 (disabled) |
| Synthetic overlap examples | GAMSA-generated (added to the training stream; see Methods) |
