## Supplementary Note 3 for "From pose to behavior: SABER integrates identity-resolved multi-animal pose tracking with large-language-model-based behavioral factor discovery"

This note presents the complete prompt system that drives the automated factor-discovery loop of SABER: the full prompt texts, the sources that fill each dynamic placeholder, and one complete real invocation. It accompanies the Methods / Software section of the paper (factor-hypothesis generation module). The prompt texts are kept identical to the prompt templates of the codebase. This note contains Supplementary Tables 7–12.

---

### Content

**Supplementary Table 7.** System-prompt modules and their functions.

**Supplementary Table 8.** Dynamic placeholders of the user-prompt template and the sources that fill them.

**Supplementary Table 9.** Feature schema injected into the {schema} placeholder.

**Supplementary Table 10.** Behavior-class definitions injected into the {behavior_rules} placeholder.

**Supplementary Table 11.** Fallback parsing levels for the LLM response.

**Supplementary Table 12.** Generation parameters and their implementation locations.

---

### Role of the Prompts in the Discovery Loop

SABER discovers behavioral factors in an iterative closed loop: *LLM hypothesis generation → sandbox execution → LightGBM validation → experience write-back*. The prompts carry the first stage (hypothesis generation):

```
┌──────────────────────────────────────────────────────────────────┐
│  MemoryManager                                                    │
│   · valid factor library (validated factors)                      │
│   · round experience (round summaries) ──┐                        │
└──────────────────────────────────────────┼────────────────────────┘
                                           │  experience summary
                                           ▼
┌──────────────────────────────────────────────────────────────────┐
│  HypothesisGenerator (prompt templates + dynamic placeholders)   │
```

```
│                                                                  │
│    SYSTEM_PROMPT                                                 │
│    + HYPOTHESIS_PROMPT_TEMPLATE                                  │
│        ├─ feature schema     ← flat_attributes / FEATURE_GROUPS│
│        ├─ behavior rules     ← hand-editable rules file         │
│        ├─ weak-class hint    ← classes with no valid factors    │
│        ├─ window config      ← seq_length                       │
│        └─ historical experience ← round experience summary      │
└──────────────────────────────────────────────────────────────────┘
                           │
                           ▼
             LLM returns a JSON array of factor hypotheses
                           ▼
┌──────────────────────────────────────────────────────────────────┐
│  FactorEngine (sandbox execution) → FactorValidator (LightGBM)  │
│  results written back to round experience → next round         │
└──────────────────────────────────────────────────────────────────┘
```

**Key design.** The prompt forces the LLM to emit *parameterized* (_P0, _P1, ...), *batch-vectorized* NumPy code rather than numeric conclusions. Every numeric constant must be written as a tunable parameter whose optimal value is later searched by an automatic parameter tuner. Behavioral knowledge is thus encoded in the *code structure*, while numeric calibration is delegated to data-driven optimization.

---

### System Prompt (Verbatim)

Defines the role, data format, coding specification, parameterization convention, and output format. Kept identical across calls.

```
You are an expert in animal behavior research, proficient in NumPy vectorize
d programming. Task: design "behavioral factors" for mouse behavior identifi
cation.

[Data Format]
- windows: np.ndarray [N, seq_length, D], full time window feature matrix. E
ach feature is normalized to [0,1].
- idx: dict, feature name → column index. Must use idx.get('name', -1) to ch
eck existence, return -1 if missing.
- np: numpy module, directly available.

[Code Specification — Batch Mode Only]
Must use batch mode (mode="batch"), processing all N windows at once with fu
lly vectorized numpy operations.
Forbidden: for loops or per-window Python logic. Variables: windows [N,seq_l
ength,D], idx, np.
Result must be assigned to _result (float array of shape [N]).
>=50% of windows must return valid values, otherwise the factor is discarde
d. No imports, no function definitions.
```

```
[Vectorized Approach for Conditional Logic]
If conditional logic is needed, use np.where:
  mask = speed_vals > _P1
  _result = np.where(mask, score * 2.0, score)

If piecewise computation is needed, use boolean indexing:
  _result = np.full(len(windows), np.nan)
  valid = (i_speed >= 0) & (i_tail >= 0)
  if valid.any():
      _result[valid] = _P0 * windows[valid, :, i_speed].mean(axis=1)

[Parameterization Convention — Important!]
Do not hardcode specific numeric constants (e.g., 0.4, 0.6, 0.35). Use _P0,
_P1, _P2... as tunable parameters,
assigning reasonable default values at the top of the code. An automatic par
ameter tuner will find optimal values later.

Parameter types and defaults:
  - Weight type (linear combination coefficient): default 1.0, search range
approx [0.05, 5.0]
  - Threshold type (condition threshold): default 0.5, search range approx
[0.01, 0.95]
  - Scaling type (amplification/attenuation factor): default 1.0, search ran
ge approx [0.1, 5.0]
  - Power type (nonlinear exponent): default 1.0, search range approx [0.5,
3.0]

Example — weight+threshold parameterization (batch vectorized):
  _P0 = 1.0   # weight coefficient for speed
  _P1 = 0.3   # activity threshold
  i_speed = idx.get('speed', -1)
  if i_speed < 0:
      _result = np.full(len(windows), np.nan)
  else:
      speed_vals = windows[:, -1, i_speed]
      base = _P0 * speed_vals
      _result = np.where(speed_vals > _P1, base * 2.0, base)

Example — temporal aggregation (batch vectorized):
  _P0 = 1.0
  i_speed = idx.get('speed', -1)
  i_tail = idx.get('tail_motion', -1)
  if i_speed < 0 or i_tail < 0:
      _result = np.full(len(windows), np.nan)
  else:
      mean_speed = np.mean(windows[:, :, i_speed], axis=1)
      std_tail = np.std(windows[:, :, i_tail], axis=1)
      _result = _P0 * mean_speed * (1.0 - std_tail)

Rule: each independently tunable value uses its own _P{n}, do not share a pa
rameter across multiple locations.
Use your best guess for initial parameter values; the tuner will override th
em.
```

```
[Output Format] Strict JSON array, each element with 5 fields:
"name" (factor_ prefix, snake_case), "description" (English), "target" (beha
vior class name),
"mode" (fixed as "batch"), "code" (Python code, batch vectorized only)
No extra fields, no text outside JSON.
```

### System prompt, section by section

### Supplementary Table 7. System-prompt modules and their functions.

| Module | Function |
|---|---|
| Role setting | Positions the model as an expert in animal behavior research and NumPy vectorized programming |
| Data format | Fixes the window matrix windows [N, seq_length, D], the feature-index dict idx, and that all features are normalized to [0,1] |
| Code specification | Enforces batch-mode full vectorization; forbids for-loops, per-window logic, imports, and function definitions; result must be written to _result[N]; ≥50% valid-value coverage |
| Conditional logic | Provides the two standard vectorization idioms: np.where and boolean indexing |
| Parameterization convention | Forbids hard-coded numeric constants; defines four parameter types (weight / threshold / scaling / power) with default values and search ranges; each tunable value gets its own _P{n} |
| Output format | Strict JSON array with exactly 5 fields per item (name, description, target, mode, code); no extra text allowed |

---

### User Prompt Template

Filled dynamically at every round. {...} placeholders and their sources are listed below.

```
[Feature Categories] (All features are normalized to [0,1], used equally, no
 preference)
{schema}

[Behavior Class Definitions]
{behavior_rules}
```

```
[Current Weak Classes] (Prioritize designing factors for these classes)
{weak_classes_hint}

[Window Configuration]
{window_context}

[Historical Experience]
{experience}

[Task] Design {n} new factors. Hard requirements:
1. Must use at least 2 features
2. Explore underused feature combinations, prioritize cross-group combinatio
ns (e.g., [Motion]+[Social], [Skeleton]+[Tail])
3. At least {n_weak} factors target weak classes
4. Do not duplicate existing factors
5. Do not repeat previously failed strategies
6. All numeric constants must use _P0, _P1... parameterization, no hardcoded
 values
7. Must use batch vectorized mode, no for loops or per-window Python logic
{temporal_requirements}
{n_start}. >=50% of windows return valid float
{n_end}. Every feature in code uses idx.get() for existence check

Output JSON array directly:
```

### Placeholder sources

### Supplementary Table 8. Dynamic placeholders of the user-prompt template and the sources that fill them.

| Placeholder | Filled from | Content / example |
|---|---|---|
| {schema} | flat_attributes from the dataset, grouped by FEATURE_GROUPS | Section 4 |
| {behavior_rules} | behavior-rules file (hand-editable ethological priors) | Section 5 |
| {weak_classes_hint} | classes with no valid factor in the previous round | With weak classes: "The following classes currently have no valid factors, please prioritize designing factors for them:\n - Climb\n - Stand"; otherwise: "(No particularly weak classes currently, explore all behavior classes evenly)" |
| {window_context} | seq_length setting | Single-frame: |

| Placeholder | Filled from | Content / example |
|---|---|---|
| | | "seq_length=1 (single frame mode, no historical frames)"; multi-frame: "seq_length=60 (multi-frame temporal mode, windows[0] earliest frame, windows[-1] latest frame)" |
| {experience} | round-experience summaries (LLM-readable rewrite) | Valid/invalid factors per round with reasons; Section 6 |
| {n} | llm.hypotheses_per_round (default 5) | number of factors requested per round |
| {n_weak} | computed as max(1, min(len(weak_classes), n // 2)); 0 if no weak classes | minimum number of factors that must target weak classes |
| {temporal_requirements} | chosen by seq_length | Single-frame: "8. Window has only 1 frame, use current frame features for mathematical combinations (ratios, differences, products, absolute values, nonlinear transformations), all vectorized"; multi-frame: "8. Window has {seq_length} frames, must utilize temporal information (e.g., np.mean/std, np.polyfit trend, first-last difference, np.max-np.min range), all vectorized" |
| {n_start} / {n_end} | fixed at 9 / 10 | continue the numbering of the hard requirements after item 8 |

**Logic of the hard requirements.** Items 1–2 guarantee combinatorial feature usage and cross-group information; item 3 steers resources toward weak classes (experience-driven); items 4–5 use historical experience to prevent duplication and the recurrence of failed strategies; items 6–7 echo the system prompt and keep the code tunable and executable; item 8 adapts to the window length (mathematical combinations for single frames, temporal statistics for multi-frame windows).

---

### Feature Schema Injected into {schema}

**Supplementary Table 9. Feature schema injected into the {schema} placeholder.**

Source: the FEATURE_GROUPS table of the hypothesis generator (synchronized with the feature registry of the feature-extraction code). The prompt asks for cross-group combinations because they carry more information than within-group permutations. The actual dataset contains exactly these 42 features.

| Group | Features |
|---|---|
| Skeleton (Self) | nose_to_head, head_to_body, body_to_tail, body_orientation, body_length, body_compactness |
| Skeleton (Other) | other_nose_to_head, other_head_to_body, other_body_to_tail, other_body_orientation, other_body_length, other_body_compactness |
| Motion (Self) | head_vel_x, head_vel_y, body_vel_x, body_vel_y, tail_vel_x, tail_vel_y, speed, acceleration |
| Motion (Other) | other_head_vel_x, other_head_vel_y, other_body_vel_x, other_body_vel_y, other_tail_vel_x, other_tail_vel_y, other_speed, other_acceleration |
| Tail (Self) | tail_angle, tail_curve, tail_motion |
| Tail (Other) | other_tail_angle, other_tail_curve, other_tail_motion |
| Social | dist_to_other, self_facing_other, other_facing_self, mutual_facing, approach_speed, relative_speed, heading_diff, body_axis_align |

The group text closes with the guidance: "Tip: prioritize cross-group feature combinations (e.g., [Motion(Self)]+[Social], [Skeleton(Self)]+[Tail(Self)]), these carry more information than permutations within the same group. Each factor must use >=2 features."

---

**Behavior Class Definitions Injected into {behavior_rules}**

**Supplementary Table 10. Behavior-class definitions injected into the {behavior_rules} placeholder.**

Source: the hand-written behavior-rules file, editable at any time. This is the main channel through which ethological priors enter the prompt.

| Class | Ethological definition (verbatim) |
|---|---|
| Approach | The subject mouse moves toward a conspecific from a distance, involving two key elements: orientation toward the other mouse AND decreasing inter-animal distance. Focus on frames where self_facing_other and approach_speed are both positive. |
| Blank | The subject mouse stands on all four paws, remaining stationary. Focus on frames where speed is near zero, all velocity features are near zero, and body_compactness is normal (not in a rearing posture). |
| Climb | The subject mouse rears up, attempting to climb the walls of the test arena. Focus on frames where body_orientation is near vertical, body_length changes (stretching), speed is low but body velocity may show specific directional patterns. |
| Explore | The subject mouse has no clear movement direction and stays continuously mobile. Focus on frames where speed > 0 but heading_diff shows high-frequency changes (no fixed direction), and dist_to_other is relatively large. |
| Sniffing | The subject mouse remains close to a conspecific and engages in sniffing behavior. Focus on frames where dist_to_other is very small, self_facing_other is high, and speed |

| Class | Ethological definition (verbatim) |
|---|---|
| | is low. |
| Self-grooming | The subject mouse is engaged in self-grooming behavior. Focus on frames where body_compactness changes (body curled up), nose_to_head distance varies (head movement), and speed is low. |
| Stand | The subject mouse maintains a bipedal rearing posture. Focus on frames where body_orientation is near vertical, nose_to_head and head_to_body ratio changes (body elongation), and speed is low. |

---

**One Complete Real Invocation (single-frame mode, seq_length = 1)**

Below is a complete message body from an actual run (placeholders filled; the experience section is abridged from the run's round-experience records).

**System** — identical to Section 2 (omitted here).

**User**:

```
[Feature Categories] (All features are normalized to [0,1], used equally, no
 preference)
Total 42 dimensions, grouped by semantics as follows:

  [Skeleton(Self)] nose_to_head, head_to_body, body_to_tail, body_orientatio
n, body_length, body_compactness
  [Skeleton(Other)] other_nose_to_head, other_head_to_body, other_body_to_ta
il, other_body_orientation, other_body_length, other_body_compactness
  [Motion(Self)] head_vel_x, head_vel_y, body_vel_x, body_vel_y, tail_vel_x,
 tail_vel_y, speed, acceleration
  [Motion(Other)] other_head_vel_x, other_head_vel_y, other_body_vel_x, othe
r_body_vel_y, other_tail_vel_x, other_tail_vel_y, other_speed, other_acceler
ation
  [Tail(Self)] tail_angle, tail_curve, tail_motion
  [Tail(Other)] other_tail_angle, other_tail_curve, other_tail_motion
  [Social] dist_to_other, self_facing_other, other_facing_self, mutual_facin
g, approach_speed, relative_speed, heading_diff, body_axis_align

Tip: prioritize cross-group feature combinations (e.g., [Motion(Self)]+[Soci
al], [Skeleton(Self)]+[Tail(Self)]),
these carry more information than permutations within the same group. Each f
actor must use >=2 features.
Use idx.get('feature_name', -1) to check existence.

[Behavior Class Definitions]
```

```
  - Approach: The subject mouse moves toward a conspecific from a distance,
involving two key elements: orientation toward the other mouse AND decreasin
g inter-animal distance. Focus on frames where self_facing_other and approac
h_speed are both positive.
  - Blank: The subject mouse stands on all four paws, remaining stationary.
Focus on frames where speed is near zero, all velocity features are near zer
o, and body_compactness is normal (not in a rearing posture).
  - Climb: The subject mouse rears up, attempting to climb the walls of the
test arena. Focus on frames where body_orientation is near vertical, body_le
ngth changes (stretching), speed is low but body velocity may show specific
directional patterns.
  - Explore: The subject mouse has no clear movement direction and stays con
tinuously mobile. Focus on frames where speed > 0 but heading_diff shows hig
h-frequency changes (no fixed direction), and dist_to_other is relatively la
rge.
  - Sniffing: The subject mouse remains close to a conspecific and engages i
n sniffing behavior. Focus on frames where dist_to_other is very small, self
_facing_other is high, and speed is low.
  - Self-grooming: The subject mouse is engaged in self-grooming behavior. F
ocus on frames where body_compactness changes (body curled up), nose_to_head
 distance varies (head movement), and speed is low.
  - Stand: The subject mouse maintains a bipedal rearing posture. Focus on f
rames where body_orientation is near vertical, nose_to_head and head_to_body
 ratio changes (body elongation), and speed is low.

[Current Weak Classes] (Prioritize designing factors for these classes)
The following classes currently have no valid factors, please prioritize des
igning factors for them:
  - Climb
  - Stand

[Window Configuration]
seq_length=1 (single frame mode, no historical frames)

[Historical Experience]
Round 1 (0 valid / 5 attempted): failed factors:
  - factor_explore_object_orientation (target=Explore): Best AUC=0.539(2) be
low threshold 0.65
  - factor_self_grooming_compact_tail (target=Self-grooming): Best AUC=0.500
(0) below threshold 0.65
  - factor_climb_vertical_axis (target=Climb): Best AUC=0.539(6) below thres
hold 0.65
  - factor_approach_relative_speed (target=Approach): Best AUC=0.586(1) belo
w threshold 0.65
  - factor_stand_still_vertical (target=Stand): Best AUC=0.548(5) below thre
shold 0.65
Round 2 (2 valid / 5 attempted): valid factors:
  - factor_climb_vertical_length_y_shift_low_social (target=Climb): Best AUC
=0.742(1)
  - factor_stand_axis_aligned_nose_lift_low_tail_activity (target=Stand): Be
st AUC=0.666(3)
  ... (older rounds omitted)

[Task] Design 5 new factors. Hard requirements:
```

```
1. Must use at least 2 features
2. Explore underused feature combinations, prioritize cross-group combinatio
ns (e.g., [Motion]+[Social], [Skeleton]+[Tail])
3. At least 2 factors target weak classes
4. Do not duplicate existing factors
5. Do not repeat previously failed strategies
6. All numeric constants must use _P0, _P1... parameterization, no hardcoded
 values
7. Must use batch vectorized mode, no for loops or per-window Python logic
8. Window has only 1 frame, use current frame features for mathematical comb
inations (ratios, differences, products, absolute values, nonlinear transfor
mations), all vectorized
9. >=50% of windows return valid float
10. Every feature in code uses idx.get() for existence check

Output JSON array directly:
```

**LLM response** (one item shown; the code of this factor exists verbatim in the factor library, its description is shown here in English translation, it targets the weak class Climb, and it satisfies requirement 3 of the task above):

```
[
  {
    "name": "factor_climb_vertical_length_y_shift_low_social",
    "description": "Climbing candidate factor: the body tends vertical and e
longates, with clear vertical body displacement, but interaction with and Ap
proach to the companion are weak, consistent with wall-facing rearing climb
s.",
    "target": "Climb",
    "mode": "batch",
    "code": "i_ori = idx.get('body_orientation', -1)\ni_len = idx.get('body_
length', -1)\ni_bvy = idx.get('body_vel_y', -1)\ni_app = idx.get('approach_s
peed', -1)\ni_sfo = idx.get('self_facing_other', -1)\ni_dist = idx.get('dist
_to_other', -1)\nif i_ori < 0 or i_len < 0 or i_bvy < 0 or i_app < 0 or i_sf
o < 0 or i_dist < 0:\n    _result = np.full(len(windows), np.nan)\nelse:\n
  ori = windows[:, 0, i_ori]\n    blen = windows[:, 0, i_len]\n    bvy = win
dows[:, 0, i_bvy]\n    app = windows[:, 0, i_app]\n    sfo = windows[:, 0, i
_sfo]\n    dist = windows[:, 0, i_dist]\n    wall_like = dist * (1 - app) *
(1 - sfo)\n    _result = np.clip(ori * blen * bvy * wall_like, 0, 1)"
  }
]
```

Notes: this factor predates the parameterization convention, so its code contains no _P parameters; factors generated after the convention was enabled follow the _P0, _P1, ... style (see, e.g., the fully parameterized code of factor_explore_object_sustained_speed_heading_tailcounter_far in Supplementary Note 6). Its per-class validation AUCs in the library are: Climb 0.7422, Sniffing 0.7159, Approach 0.6262. After passing validation it was saved to the factor library and injected into the next round's experience.

### Response Parsing and Robustness

**Supplementary Table 11. Fallback parsing levels for the LLM response.**

The LLM output is parsed by HypothesisGenerator._parse_response in four fallback levels, which keeps the loop robust to imperfect JSON:

| Level | Strategy | Notes |
|---|---|---|
| 1 | Direct json.loads | after stripping json ``fences and trailing backticks \| \| 2 \| Regex-extract the first[...]` array |
| 3 | Truncated-array rescue | scans bracket depth and recovers individual objects from truncated output |
| 4 | Abandon the round | logged warning, empty list returned, next round proceeds |

Each recovered object then passes field-level validation: field aliases are accepted (factor_name, target_behavior, target_class, behavior, behavior_class, formula, implementation); objects missing name, description or code are dropped; the name is normalized to factor_-prefixed snake_case. The LLM client adds automatic retries on empty responses or call failures (retry interval 5 s).

---

### Generation Parameters and Implementation Locations

**Supplementary Table 12. Generation parameters and their implementation locations.**

| Item | Value / location |
|---|---|
| Prompt templates | prompt builder: SYSTEM_PROMPT, HYPOTHESIS_PROMPT_TEMPLATE, _SINGLE_FRAME_REQUIREMENTS, _TEMPORAL_REQUIREMENTS_TEMPLATE |
| Feature groups | prompt builder: FEATURE_GROUPS |
| Ethological priors | behavior-rules file (hand-editable) |
| Experience injection | get_experience_summary over the round-experience store (latest 300 |

| Item | Value / location |
| --- | --- |
| | rounds kept, last 200 injected) |
| Factors per round | llm.hypotheses_per_round (default 5) |
| Sampling parameters | llm.temperature 0.7, llm.max_tokens 16,384 (as used in the reported run; code default 4,096) |
| Model backends | llm.model — OpenAI-compatible / GLM / Anthropic, unified by a single client module |
