## Supplementary Note 4 for "From pose to behavior: SABER integrates identity-resolved multi-animal pose tracking with large-language-model-based behavioral factor discovery"

This note gives the exact definitions of the 42 per-frame features computed from the two tracked mice. These features form the columns of the window matrix windows [N, seq_length, D] exposed to all factor code, and are the only inputs the LLM-generated factors are allowed to use. Definitions are reproduced verbatim from the feature registry of the feature-extraction code. This note contains Supplementary Tables 13–16.

---

### Content

**Supplementary Table 13.** Skeleton-group features (six per mouse, twelve in total).

**Supplementary Table 14.** Motion-group features (eight per mouse, sixteen in total).

**Supplementary Table 15.** Tail-group features (three per mouse, six in total).

**Supplementary Table 16.** Social-group features (eight, inter-animal).

---

### Notation

- Mouse keypoints: nose (0), head (1), body (2), tail base (3); tail midpoints and tip are tracked in the tail keypoint stream (mid = kp 7, tip = kp 9 of the combined pose).
- pts[m] = keypoint coordinates of mouse m at frame t (normalized image coordinates); box[m] = bounding-box center of mouse m; box_vel[m], box_acc[m] = its velocity and acceleration.
- "Other" = the companion mouse; all other-* features use the same definitions applied to the companion.
- Angles follow atan2 conventions; angle differences are wrapped to $(-\pi, \pi]$.

---

**Skeleton Group (6 per mouse, 12 total)**

**Supplementary Table 13. Skeleton-group features (six per mouse, twelve in total).**

| Feature | Definition |
|---|---|
| nose_to_head | ‖ nose − head ‖ (distance between nose and head keypoints) |
| head_to_body | ‖ head − body ‖ |
| body_to_tail | ‖ body − tail base ‖ |
| body_orientation | atan2( head − body ) — direction of the head–body vector |
| body_length | ‖ nose − tail base ‖ — full body extent |
| body_compactness | mean pairwise distance among the 4 keypoints {nose, head, body, tail base} — small when the body is curled up |

Companion versions: other_nose_to_head, other_head_to_body, other_body_to_tail, other_body_orientation, other_body_length, other_body_compactness.

---

**Motion Group (8 per mouse, 16 total)**

**Supplementary Table 14. Motion-group features (eight per mouse, sixteen in total).**

| Feature | Definition |
|---|---|
| head_vel_x / head_vel_y | x/y velocity of the head keypoint |
| body_vel_x / body_vel_y | x/y velocity of the body keypoint |
| tail_vel_x / tail_vel_y | x/y velocity of the tail-base keypoint |
| speed | ‖ box_vel ‖ — magnitude of bounding-box center velocity |
| acceleration | ‖ box_acc ‖ — magnitude of bounding-box center acceleration |

Companion versions carry the other_ prefix.

---

**Tail Group (3 per mouse, 6 total)**

**Supplementary Table 15. Tail-group features (three per mouse, six in total).**

| Feature | Definition |
|---|---|

| Feature | Definition |
|---|---|
| tail_angle | wrapped angle between the tail vector (tail base → tail tip) and the body vector (head → body) |
| tail_curve | ‖ tail midpoint − midpoint of the straight tail base–tip segment ‖ — deviation of the tail from a straight line |
| tail_motion | ‖ velocity of the tail tip keypoint ‖ |

Companion versions: other_tail_angle, other_tail_curve, other_tail_motion.

---

**Social Group (8, inter-animal)**

**Supplementary Table 16. Social-group features (eight, inter-animal).**

| Feature | Definition |
|---|---|
| dist_to_other | ‖ box(other) − box(self) ‖ — distance between bounding-box centers |
| self_facing_other | cos( heading(self) − direction to other's body ) — 1 when the subject faces the companion |
| other_facing_self | cos( heading(other) − direction to self's body ) |
| mutual_facing | self_facing_other × other_facing_self — 1 when both mice face each other |
| approach_speed | − ( v_rel · $\hat{d}$ ) — projection of relative velocity onto the self→other direction (positive when closing in) |
| relative_speed | ‖ v_self − v_other ‖ |
| heading_diff | ‖ heading(self) − heading(other) ‖, wrapped to [0, π] — 0 when both mice move in the same direction |
| body_axis_align | cos( heading(self) − heading(other) ) — 1 when body axes are parallel |

---

**Normalization and Availability**

- Every feature is percentile-clipped to [0.1%, 99.9%] (fitted on the training set) and then min-max scaled to [0, 1].
- Missing keypoints (tracking failures) are interpolated bidirectionally (short gaps ≤ 10 frames); remaining gaps produce NaN frames that factor code must handle via idx.get('feature', -1) existence checks and NaN propagation.

- Factor code accesses these columns through the idx name→index dictionary exactly as named above (e.g., idx.get('speed', -1)).

---
