## Supplementary Note 5 for "From pose to behavior: SABER integrates identity-resolved multi-animal pose tracking with large-language-model-based behavioral factor discovery"

This note describes the multi-resolution three-stage validation architecture, the full hyperparameter set of each stage, the final model's validation metrics, and a reproducibility summary. This note contains Supplementary Tables 17–20.

---

### Content


**Supplementary Table 17.** Stage-1 per-group LightGBM hyperparameters.

**Supplementary Table 18.** Stage-3 temporal model (BiLSTM) configuration.

**Supplementary Table 19.** Final model per-class validation metrics (validation set, 360,008 frames).

**Supplementary Table 20.** Reproducibility summary.


---

### Multi-Resolution Three-Stage Validation Architecture

Factors are grouped by their temporal window length and validated in three stages:

```
Raw frames [T, D]
  ├── Short-range factor group (seq_length ≤ 3)  ──▶ LGBM_short  ──▶ proba
[T, C]
  ├── Medium-range factor group (seq_length 4–15) ──▶ LGBM_medium ──▶ proba
[T, C]
  └── Long-range factor group (seq_length > 15)  ──▶ LGBM_long   ──▶ proba
[T, C]
                                                         │ hstack
                                                meta_X = [T, C×n_groups]
                                                         │
                                                    meta-LGBM
                                                         │
                                          Temporal sliding-window features (31
frames)
                                                         │
                                                Temporal BiLSTM (second-order)
```

### Stage 1 — per-group LightGBM (factor matrices)

#### Supplementary Table 17. Stage-1 per-group LightGBM hyperparameters.

| Hyperparameter | Value |
|---|---|

| Hyperparameter | Value |
|---|---|
| Factors per group | short 595 / medium 1,200 / long 1,153 (2,948 factors in total; top-k by validation AUC) |
| Model | LightGBM multi-class, n_estimators 500, learning_rate 0.05, num_leaves 63, max_depth 4 |
| Class handling | class_weight = "balanced", focal loss ($\alpha$ = 0.25, $\gamma$ = 2.0) |
| NaN handling | mean fill (fit on training set) + standardization, clipped to [−8, 8] |
| Early stopping | 50 rounds |
| Residual connections | stage-1 residual enabled (residual_stage1), meta-stage residual enabled (residual_stage2) |
| Seed | 42 |

**Stage 2 — meta-LightGBM stacking**

Class-aligned probability vectors of the three group models are concatenated ([T, C×3]) and fed to a meta-LightGBM (same objective and hyperparameters as Stage 1), producing the stacking probabilities.

**Stage 3 — temporal second-order model (BiLSTM)**

**Supplementary Table 18. Stage-3 temporal model (BiLSTM) configuration.**

| Item | Value |
|---|---|
| Model | BiLSTM, hidden_dim 1024, num_layers 4, dropout 0.3 |
| Input | sliding 1,024-frame chunks (stride 512 training / 256 validation) of meta-stacking probabilities (7 classes) + enriched features (2,976 dims = 7 + 2,969) |
| Enriched features | delta, entropy, margin features; multi-scale windows [7, 15, 31, 63]; autocorrelation; cross-scale; distribution-shape statistics; factor-matrix extra features (2,969 dims) |
| Training | 200 epochs, lr 0.0005, weight_decay 0.0005, batch size 64, focal loss ($\gamma$ = 2.0, class-specific $\alpha$ weights from inverse class frequency) + class weights, label smoothing 0.15 |

| Item | Value |
| --- | --- |
| | (ignored under focal loss), mixup 0.2, AMP |
| Output residual | enabled; early-stopping patience 20; seed 42 |
| Post-decoder | rule-based consistency correction (rule_correct): flicker suppression (max 3 frames, proba threshold 0.05), minimum consecutive segment 5 frames, boundary refinement, confidence margin 0.3 |

**Single-factor validation (discovery phase)**

Each candidate factor is validated independently on a video-level training–validation holdout: a one-versus-rest LightGBM classifier is fitted for each behavior class on the designated training videos and evaluated on the held-out validation videos (K-fold cross-validation is not used within the mining loop). A factor is accepted when its best class AUC ≥ 0.65 (the tuned library's minimum best AUC is 0.6502); the F1-score is recorded but is not used as an acceptance gate. The single-factor classifier used n_estimators = 200, max_depth = 4, learning_rate = 0.05, num_leaves = 31, random_state = 42, and n_jobs = 1, with early stopping on the validation AUC (early_stopping_rounds = 20). The negative class was randomly under-sampled to a 1:1 positive-to-negative ratio for training. Multi-factor joint validation of the assembled factor library, by contrast, used 5-fold stratified cross-validation (StratifiedKFold, one-vs-rest LightGBM per class).

---

**Final Model Validation Metrics (validation set, 360,008 frames)**

**Supplementary Table 19. Final model per-class validation metrics (validation set, 360,008 frames).**

The temporal BiLSTM with post-hoc calibration and rule-based correction (Calib+rule_correct) was selected as the final model (monitor: validation balanced accuracy, 0.6966).

| Class | Precision | Recall | F1 | AUC | Support |
| --- | --- | --- | --- | --- | --- |
| Explore | 0.8498 | 0.8520 | 0.8509 | 0.9358 | 101,827 |
| Climb | 0.6502 | 0.7789 | 0.7087 | 0.9568 | 13,851 |
| Self-grooming | 0.5985 | 0.7870 | 0.6799 | 0.9161 | 54,777 |
| Stand | 0.7078 | 0.7589 | 0.7325 | 0.8672 | 89,285 |

| Class | Precision | Recall | F1 | AUC | Support |
|---|---|---|---|---|---|
| Blank | 0.7080 | 0.4840 | 0.5750 | 0.9074 | 70,725 |
| Sniffing | 0.6754 | 0.5786 | 0.6232 | 0.8502 | 19,703 |
| Approach | 0.7525 | 0.6369 | 0.6899 | 0.8889 | 9,840 |
| **Macro** | 0.7060 | 0.6966 | **0.6943** | **0.9032** | |
| **Weighted** | 0.7286 | 0.7231 | **0.7190** | **0.9051** | |

Overall: accuracy 0.7231, balanced accuracy 0.6966. The full confusion matrix (raw counts and row-normalized) is shown in the corresponding heatmap figures.

For comparison, the Stage-2 stacking model (meta-LightGBM, before the temporal stage) reached substantially lower macro-F1 and macro-AUC on a down-sampled evaluation set.

---

**Reproducibility Summary**

**Supplementary Table 20. Reproducibility summary.**

| Item | Value |
|---|---|
| Random seeds | 42 (LightGBM, BiLSTM, splits) |
| Factor computation | 24 workers, 256 chunks, memmap-shared keypoints |
| GPU usage | LightGBM GPU enabled (use_gpu: true), BiLSTM on CUDA with AMP |
| Configurations | 20260626_171940/configs/common.yaml + 20260626_171940/configs/validation.yaml (merged snapshot: 20260626_171940/configs/merged_config.yaml) |
| Pipeline caches | standardized factor matrices cached in memmap-backed per-stage caches |

---
