## Supplementary Note 6 for "From pose to behavior: SABER integrates identity-resolved multi-animal pose tracking with large-language-model-based behavioral factor discovery"

This note records the complete content of the 35 factors that ranked in the top 5 of each of the seven behavior classes by one-versus-rest validation AUC (7 classes × 5 factors) within the released 2,948-factor library, supporting Fig. 4g and the Results section "LLM-based behavioral factor mining generates interpretable representations from pose dynamics". Factors are grouped by the class they were selected for. This note contains Supplementary Tables 21–34.

---

#### Content

**Supplementary Table 21.** Top-five factors selected for Explore.

**Supplementary Table 22.** One-versus-rest validation AUC of the five leading Explore factors across all seven behavior classes.

**Supplementary Table 23.** Top-five factors selected for Climb.

**Supplementary Table 24.** One-versus-rest validation AUC of the five leading Climb factors across all seven behavior classes.

**Supplementary Table 25.** Top-five factors selected for Self-grooming.

**Supplementary Table 26.** One-versus-rest validation AUC of the five leading Self-grooming factors across all seven behavior classes.

**Supplementary Table 27.** Top-five factors selected for Stand.

**Supplementary Table 28.** One-versus-rest validation AUC of the five leading Stand factors across all seven behavior classes.

**Supplementary Table 29.** Top-five factors selected for Blank.

**Supplementary Table 30.** One-versus-rest validation AUC of the five leading Blank factors across all seven behavior classes.

**Supplementary Table 31.** Top-five factors selected for Sniffing.

**Supplementary Table 32.** One-versus-rest validation AUC of the five leading Sniffing factors across all seven behavior classes.

**Supplementary Table 33.** Top-five factors selected for Approach.

**Supplementary Table 34.** One-versus-rest validation AUC of the five leading Approach factors across all seven behavior classes.

---

**Explore (class index 0)**

**Supplementary Table 21. Top-five factors selected for Explore.**

Factors are ordered by their one-versus-rest validation AUC on the selected class (Explore); the selected AUC is the class-specific AUC used for ranking.

| Rank | Factor name | Selected AUC | Designed target | seq_length |
|---|---|---|---|---|
| 1 | factor_explore_object_sustained_speed_heading_tailcounter_far | 0.7660 | Explore | 60 |
| 2 | factor_explore_object_wandering_far_social_unlocked | 0.7627 | Explore | 60 |
| 3 | factor_explore_object_far_turning_accel_bodytail_variability | 0.7615 | Explore | 60 |
| 4 | factor_explore_object_far_wander_heading_othertail_independent | 0.7607 | Explore | 60 |
| 5 | factor_explore_object_heading_jitter_accel_far_partnerquiet | 0.7595 | Explore | 60 |

**Supplementary Table 22. One-versus-rest validation AUC of the five leading Explore factors across all seven behavior classes.**

Rows correspond to the ranked factors listed in Supplementary Table 21. Bold values indicate the class-specific AUC on which each factor was selected. Factor descriptions, validation details, and executable code are provided after each table.

| Rank | Explore | Climb | Self-grooming | Stand | Blank | Sniffing | Approach |
|---|---|---|---|---|---|---|---|
| 1 | **0.7660** | 0.7207 | 0.5768 | 0.5436 | 0.6371 | 0.5441 | 0.7694 |

| Rank | Explore | Climb | Self-grooming | Stand | Blank | Sniffing | Approach |
|---|---|---|---|---|---|---|---|
| 2 | **0.7627** | 0.7517 | 0.6182 | 0.5804 | 0.6734 | 0.5039 | 0.7881 |
| 3 | **0.7615** | 0.7543 | 0.6185 | 0.5840 | 0.6651 | 0.4956 | 0.7944 |
| 4 | **0.7607** | 0.6898 | 0.6159 | 0.5629 | 0.6716 | 0.5156 | 0.7600 |
| 5 | **0.7595** | 0.8102 | 0.6490 | 0.5884 | 0.6766 | 0.4897 | 0.8251 |

**factor_explore_object_sustained_speed_heading_tailcounter_far (rank 1, AUC 0.7660)**

**Description.** During object exploration the mouse sustains locomotion with an unstable heading, accompanied by orientation changes and tail counter-balance adjustments, while remaining far from the companion with low mutual facing.

- Designed target: **Explore** · Mode: batch · seq_length: 60 · n_samples: 3595
- Best class: Approach (AUC 0.7694, F1 0.0240)
- Valid classes: Approach (AUC 0.7694, F1 0.0240), Explore (AUC 0.7660, F1 0.4826), Climb (AUC 0.7207, F1 0.0181)

**Code.**

```
_P0 = 1.0
_P1 = 1.0
_P2 = 1.0
_P3 = 1.0
_P4 = 1.0
_P5 = 1.0
_P6 = 1.0
_P7 = 0.5
_P8 = 1.0
i_speed = idx.get('speed', -1)
i_heading = idx.get('heading_diff', -1)
i_tail_motion = idx.get('tail_motion', -1)
i_dist = idx.get('dist_to_other', -1)
i_mutual = idx.get('mutual_facing', -1)
i_acc = idx.get('acceleration', -1)
if i_speed < 0 or i_heading < 0 or i_tail_motion < 0 or i_dist < 0 or i_mutu
al < 0 or i_acc < 0:
    _result = np.full(len(windows), np.nan)
else:
    sustained_speed = np.mean(windows[:, :, i_speed], axis=1) - np.std(windo
ws[:, :, i_speed], axis=1)
    heading_jitter = np.std(windows[:, :, i_heading], axis=1) + np.ptp(windo
ws[:, :, i_heading], axis=1)
    tail_counter = np.mean(windows[:, :, i_tail_motion], axis=1) + np.std(wi
ndows[:, :, i_tail_motion], axis=1)
    far_other = np.mean(windows[:, :, i_dist], axis=1)
```

```
    low_social = _P4 - np.mean(windows[:, :, i_mutual], axis=1)
    accel_var = np.std(windows[:, :, i_acc], axis=1)
    _result = (_P0 * np.maximum(sustained_speed, _P7) * np.power(heading_jit
ter, _P1) + _P2 * tail_counter + _P3 * accel_var) * far_other * np.maximum(l
ow_social, _P7) * _P5 + _P6 * np.ptp(windows[:, :, i_speed], axis=1) * _P8
```

#### factor_explore_object_wandering_far_social_unlocked (rank 2, AUC 0.7627)

**Description.** Identifies exploration without a fixed direction: sustained locomotion far from the companion with fluctuating heading difference and no stable social orientation.

- Designed target: **Explore** · Mode: batch · seq_length: 60 · n_samples: 3595
- Best class: Approach (AUC 0.7881, F1 0.0220)
- Valid classes: Approach (AUC 0.7881, F1 0.0220), Explore (AUC 0.7627, F1 0.5244), Climb (AUC 0.7517, F1 0.0274), Blank (AUC 0.6734, F1 0.5346)

**Code.**

```
_P0 = 1.0
_P1 = 1.0
_P2 = 1.0
_P3 = 1.0
_P4 = 1.0
_P5 = 1.0
_P6 = 0.5
i_speed = idx.get('speed', -1)
i_heading = idx.get('heading_diff', -1)
i_dist = idx.get('dist_to_other', -1)
i_mutual = idx.get('mutual_facing', -1)
i_tail = idx.get('tail_motion', -1)
if i_speed < 0 or i_heading < 0 or i_dist < 0 or i_mutual < 0 or i_tail < 0:
    _result = np.full(len(windows), np.nan)
else:
    speed_m = np.mean(windows[:, :, i_speed], axis=1)
    heading_std = np.std(windows[:, :, i_heading], axis=1)
    heading_rng = np.max(windows[:, :, i_heading], axis=1) - np.min(windows
[:, :, i_heading], axis=1)
    dist_m = np.mean(windows[:, :, i_dist], axis=1)
    mutual_m = np.mean(windows[:, :, i_mutual], axis=1)
    tail_m = np.mean(windows[:, :, i_tail], axis=1)
    wander = _P0 * heading_std + _P1 * heading_rng
    mobile = _P2 * speed_m + _P3 * tail_m
    social_free = (_P4 * dist_m) * (1.0 / (1.0 + _P5 * mutual_m))
    _result = mobile * wander * social_free
    _result = np.where(speed_m > _P6, _result, _result * speed_m)
```

**factor_explore_object_far_turning_accel_bodytail_variability (rank 3, AUC 0.7615)**

**Description.** Object-exploration factor: combines distance from the companion, sustained locomotion, heading-difference changes, acceleration fluctuation, and tail-curvature variation to describe active exploration without a fixed direction.

- Designed target: **Explore** · Mode: batch · seq_length: 60 · n_samples: 3595
- Best class: Approach (AUC 0.7944, F1 0.0229)
- Valid classes: Approach (AUC 0.7944, F1 0.0229), Explore (AUC 0.7615, F1 0.5284), Climb (AUC 0.7543, F1 0.0240), Blank (AUC 0.6651, F1 0.5298)

**Code.**

```
_P0 = 1.0
_P1 = 1.0
_P2 = 1.0
_P3 = 1.0
_P4 = 1.0
_P5 = 1.0
_P6 = 0.5
i_dist = idx.get('dist_to_other', -1)
i_speed = idx.get('speed', -1)
i_head = idx.get('heading_diff', -1)
i_acc = idx.get('acceleration', -1)
i_tailc = idx.get('tail_curve', -1)
i_align = idx.get('body_axis_align', -1)
if i_dist < 0 or i_speed < 0 or i_head < 0 or i_acc < 0 or i_tailc < 0 or i_
align < 0:
    _result = np.full(len(windows), np.nan)
else:
    dist = windows[:, :, i_dist]
    spd = windows[:, :, i_speed]
    hd = windows[:, :, i_head]
    acc = windows[:, :, i_acc]
    tc = windows[:, :, i_tailc]
    align = windows[:, :, i_align]
    far = np.mean(dist, axis=1)
    move = np.mean(spd, axis=1)
    turning = np.mean(np.abs(np.diff(hd, axis=1)), axis=1) + _P0 * np.std(h
d, axis=1)
    accel_var = np.std(acc, axis=1)
    tail_var = np.std(tc, axis=1)
    non_social_align = np.clip(1.0 - np.mean(align, axis=1), 0.0, 1.0)
    speed_persist = np.clip(1.0 - np.std(spd, axis=1), 0.0, 1.0)
    score = (_P1 * far + _P2 * move + _P3 * turning + _P4 * accel_var + _P5
* tail_var) * (non_social_align + speed_persist)
```

```
    score = np.where(move > _P6, score, score * move)
    _result = score.astype(float)
```

#### factor_explore_object_far_wander_heading_othertail_independent (rank 4, AUC 0.7607)

**Description.** Object-exploration factor: the mouse stays mobile far from the companion with large within-window heading-difference fluctuations, and its own tail activity exceeds the companion's tail background, capturing non-social, direction-unstable autonomous exploration.

- Designed target: **Explore** · Mode: batch · seq_length: 60 · n_samples: 3595
- Best class: Explore (AUC 0.7607, F1 0.5205)
- Valid classes: Explore (AUC 0.7607, F1 0.5205), Approach (AUC 0.7600, F1 0.0197), Climb (AUC 0.6898, F1 0.0165), Blank (AUC 0.6716, F1 0.5308)

**Code.**

```
_P0 = 1.0
_P1 = 1.0
_P2 = 1.0
_P3 = 1.0
_P4 = 1.0
_P5 = 1.0
_P6 = 1.0
_P7 = 1.0
_P8 = 1.0

i_speed = idx.get('speed', -1)
i_heading = idx.get('heading_diff', -1)
i_dist = idx.get('dist_to_other', -1)
i_face = idx.get('self_facing_other', -1)
i_tail = idx.get('tail_motion', -1)
i_otail = idx.get('other_tail_motion', -1)
i_rel = idx.get('relative_speed', -1)
if min(i_speed, i_heading, i_dist, i_face, i_tail, i_otail, i_rel) < 0:
    _result = np.full(len(windows), np.nan)
else:
    locomotion = np.mean(windows[:, :, i_speed], axis=1)
    heading_var = np.std(windows[:, :, i_heading], axis=1) + np.max(windows
[:, :, i_heading], axis=1) - np.min(windows[:, :, i_heading], axis=1)
    far = np.mean(windows[:, :, i_dist], axis=1)
    unfaced = _P0 - np.mean(windows[:, :, i_face], axis=1)
    self_tail_active = np.mean(windows[:, :, i_tail], axis=1) + np.std(windo
ws[:, :, i_tail], axis=1)
    other_tail_background = _P1 - np.mean(windows[:, :, i_otail], axis=1)
    relative_low = _P2 - np.mean(windows[:, :, i_rel], axis=1)
    _result = (_P3 * locomotion + _P4 * self_tail_active) * (_P5 * heading_v
```

```
ar) * (_P6 * far + _P7 * unfaced) * (_P8 * other_tail_background + relative_
low)
```

#### factor_explore_object_heading_jitter_accel_far_partnerquiet (rank 5, AUC 0.7595)

**Description.** Sustained locomotion with large heading changes, while keeping distance from the companion whose motion interference is low, capturing object/environment exploration without a fixed direction.

- Designed target: **Explore** · Mode: batch · seq_length: 60 · n_samples: 3595
- Best class: Approach (AUC 0.8251, F1 0.0281)
- Valid classes: Approach (AUC 0.8251, F1 0.0281), Climb (AUC 0.8102, F1 0.0449), Explore (AUC 0.7595, F1 0.5240), Blank (AUC 0.6766, F1 0.5239)

**Code.**

```
_P0 = 1.0
_P1 = 1.0
_P2 = 1.0
_P3 = 1.0
_P4 = 1.0
_P5 = 0.5
i_speed = idx.get('speed', -1)
i_heading = idx.get('heading_diff', -1)
i_acc = idx.get('acceleration', -1)
i_dist = idx.get('dist_to_other', -1)
i_other_acc = idx.get('other_acceleration', -1)
if i_speed < 0 or i_heading < 0 or i_acc < 0 or i_dist < 0 or i_other_acc <
0:
    _result = np.full(len(windows), np.nan)
else:
    sp = windows[:, :, i_speed]
    hd = windows[:, :, i_heading]
    ac = windows[:, :, i_acc]
    ds = windows[:, :, i_dist]
    oa = windows[:, :, i_other_acc]
    sustained_move = np.mean(sp, axis=1)
    heading_jitter = np.std(hd, axis=1) + np.max(hd, axis=1) - np.min(hd, ax
is=1)
    accel_var = np.std(ac, axis=1)
    far_social = np.mean(ds, axis=1)
    other_quiet = 1.0 - np.mean(oa, axis=1)
    base = _P0 * sustained_move + _P1 * heading_jitter + _P2 * accel_var + _
P3 * far_social + _P4 * other_quiet
    _result = np.where(sustained_move > _P5, base, base * sustained_move)
```

**Climb (class index 1)**

**Supplementary Table 23. Top-five factors selected for Climb.**

Factors are ordered by their one-versus-rest validation AUC on the selected class (Climb); the selected AUC is the class-specific AUC used for ranking.

| Rank | Factor name | Selected AUC | Designed target | seq_length |
|---|---|---|---|---|
| 1 | factor_self_grooming_compact_headpulse_tailcurve_pause | 0.8892 | Self-grooming | 60 |
| 2 | factor_self_grooming_head_tail_alternation_lowlocomotion | 0.8714 | Self-grooming | 60 |
| 3 | factor_blank_full_body_micro_velocity_compact_constancy | 0.8592 | Blank | 60 |
| 4 | factor_blank_allsegment_velocity_silence_compact_regular | 0.8524 | Blank | 60 |
| 5 | factor_blank_global_velocity_still_compact_stable | 0.8522 | Blank | 60 |

**Supplementary Table 24. One-versus-rest validation AUC of the five leading Climb factors across all seven behavior classes.**

Rows correspond to the ranked factors listed in Supplementary Table 23. Bold values indicate the class-specific AUC on which each factor was selected. Factor descriptions, validation details, and executable code are provided after each table.

| Rank | Explore | Climb | Self-grooming | Stand | Blank | Sniffing | Approach |
|---|---|---|---|---|---|---|---|
| 1 | 0.6206 | **0.8892** | 0.5793 | 0.5136 | 0.6089 | 0.5229 | 0.6875 |

| Rank | Explore | Climb | Self-grooming | Stand | Blank | Sniffing | Approach |
|---|---|---|---|---|---|---|---|
| 2 | 0.6281 | **0.8714** | 0.6913 | 0.5008 | 0.5640 | 0.5047 | 0.6701 |
| 3 | 0.5989 | **0.8592** | 0.5349 | 0.5203 | 0.5475 | 0.5326 | 0.4151 |
| 4 | 0.6969 | **0.8524** | 0.5861 | 0.5540 | 0.5849 | 0.4907 | 0.7747 |
| 5 | 0.5280 | **0.8522** | 0.5147 | 0.5305 | 0.5502 | 0.4664 | 0.5576 |

**factor_self_grooming_compact_headpulse_tailcurve_pause (rank 1, AUC 0.8892)**

**Description.** Self-grooming factor: a compact body with local variations in the nose-head segment and tail curvature, combined with low overall speed and weak social approach tendency, capturing in-place grooming movements.

- Designed target: **Self-grooming** · Mode: batch · seq_length: 60 · n_samples: 3595
- Best class: Climb (AUC 0.8892, F1 0.0561)
- Valid classes: Climb (AUC 0.8892, F1 0.0561), Approach (AUC 0.6875, F1 0.0191)

**Code.**

```
_P0 = 1.0  # body_compactness mean weight
_P1 = 1.0  # nose_to_head std weight
_P2 = 1.0  # tail_curve std weight
_P3 = 1.0  # speed low-motion weight
_P4 = 1.0  # approach_speed low social-approach weight
_P5 = 1.0  # head_vel_x head micro-motion weight
_P6 = 1.0  # head_vel_y head micro-motion weight
_P7 = 1.0  # high-value anchor
i_comp = idx.get('body_compactness', -1)
i_nh = idx.get('nose_to_head', -1)
i_tailc = idx.get('tail_curve', -1)
i_speed = idx.get('speed', -1)
i_app = idx.get('approach_speed', -1)
i_hvx = idx.get('head_vel_x', -1)
i_hvy = idx.get('head_vel_y', -1)
if min([i_comp, i_nh, i_tailc, i_speed, i_app, i_hvx, i_hvy]) < 0:
    _result = np.full(len(windows), np.nan)
else:
    comp_mean = np.mean(windows[:, :, i_comp], axis=1)
    nh_std = np.std(windows[:, :, i_nh], axis=1)
    tailc_std = np.std(windows[:, :, i_tailc], axis=1)
    speed_mean = np.mean(windows[:, :, i_speed], axis=1)
    app_mean = np.mean(windows[:, :, i_app], axis=1)
    hvx_std = np.std(windows[:, :, i_hvx], axis=1)
    hvy_std = np.std(windows[:, :, i_hvy], axis=1)
    _result = (_P0 * comp_mean + _P1 * nh_std + _P2 * tailc_std + _P3 * (_P7
 - speed_mean) + _P4 * (_P7 - app_mean) + _P5 * hvx_std + _P6 * hvy_std).ast
ype(float)
```

**factor_self_grooming_head_tail_alternation_lowlocomotion (rank 2, AUC 0.8714)**

**Description.** Uses within-window fluctuations of head velocity and tail motion together with low overall displacement and weak social facing to describe the non-locomotor, in-place local movements of self-grooming.

- Designed target: **Self-grooming** · Mode: batch · seq_length: 60 · n_samples: 3595
- Best class: Climb (AUC 0.8714, F1 0.0389)
- Valid classes: Climb (AUC 0.8714, F1 0.0389), Self-grooming (AUC 0.6913, F1 0.4583), Approach (AUC 0.6701, F1 0.0161)

**Code.**

```
_P0 = 1.0
_P1 = 1.0
_P2 = 1.0
_P3 = 1.0
_P4 = 1.0
_P5 = 1.0
_P6 = 1.0
_P7 = 1.0
_P8 = 0.5
i_hx = idx.get('head_vel_x', -1)
i_hy = idx.get('head_vel_y', -1)
i_tailmotion = idx.get('tail_motion', -1)
i_compact = idx.get('body_compactness', -1)
i_speed = idx.get('speed', -1)
i_face = idx.get('self_facing_other', -1)
if i_hx < 0 or i_hy < 0 or i_tailmotion < 0 or i_compact < 0 or i_speed < 0
or i_face < 0:
    _result = np.full(len(windows), np.nan)
else:
    head_micro = np.std(windows[:, :, i_hx], axis=1) + np.std(windows[:, :,
i_hy], axis=1)
    tail_var = np.std(windows[:, :, i_tailmotion], axis=1)
    compact_var = np.std(windows[:, :, i_compact], axis=1)
    compact_level = np.mean(windows[:, :, i_compact], axis=1)
    low_locomotion = _P4 / (_P4 + np.mean(windows[:, :, i_speed], axis=1) **
 _P5)
    social_disengaged = _P6 / (_P6 + np.mean(windows[:, :, i_face], axis=1)
** _P7)
    raw = _P0 * head_micro + _P1 * tail_var + _P2 * compact_var + _P3 * comp
act_level
    _result = raw * low_locomotion * social_disengaged
    _result = np.where(compact_level > _P8, _result, _result * _P8)
```

**factor_blank_full_body_micro_velocity_compact_constancy (rank 3, AUC 0.8592)**

**Description.** Speed and the velocity components of the body, head and tail are all at low levels, with small changes in body compactness and posture, indicating a four-paw standing, stationary state.

- Designed target: **Blank** · Mode: batch · seq_length: 60 · n_samples: 3595
- Best class: Climb (AUC 0.8592, F1 0.0228)
- Valid classes: Climb (AUC 0.8592, F1 0.0228)

**Code.**

```
ispeed = idx.get('speed', -1)
ibvx = idx.get('body_vel_x', -1)
ibvy = idx.get('body_vel_y', -1)
ihvx = idx.get('head_vel_x', -1)
ihvy = idx.get('head_vel_y', -1)
itvx = idx.get('tail_vel_x', -1)
itvy = idx.get('tail_vel_y', -1)
ibc = idx.get('body_compactness', -1)
ibo = idx.get('body_orientation', -1)
if ispeed < 0 or ibvx < 0 or ibvy < 0 or ihvx < 0 or ihvy < 0 or itvx < 0 or
 itvy < 0 or ibc < 0 or ibo < 0:
    _result = np.full(len(windows), np.nan)
else:
    vel_mean = (np.mean(windows[:, :, ispeed], axis=1) + np.mean(windows[:,
 :, ibvx], axis=1) + np.mean(windows[:, :, ibvy], axis=1) + np.mean(windows
[:, :, ihvx], axis=1) + np.mean(windows[:, :, ihvy], axis=1) + np.mean(windo
ws[:, :, itvx], axis=1) + np.mean(windows[:, :, itvy], axis=1)) / 7.0
    compact_stable = 1.0 - np.std(windows[:, :, ibc], axis=1)
    orient_stable = 1.0 - np.std(windows[:, :, ibo], axis=1)
    compact_normal = 1.0 - np.abs(np.mean(windows[:, :, ibc], axis=1) - 0.5)
 * 2.0
    compact_normal = np.maximum(0.0, compact_normal)
    _result = (1.0 - vel_mean) * compact_stable * orient_stable * compact_no
rmal
```

**factor_blank_allsegment_velocity_silence_compact_regular (rank 4, AUC 0.8524)**

**Description.** Stationary blank factor: head, body and tail segment velocities are all low, overall speed and acceleration are low, and body compactness stays stable; whole-window velocity variance excludes brief movements.

- Designed target: **Blank** · Mode: batch · seq_length: 60 · n_samples: 3595
- Best class: Climb (AUC 0.8524, F1 0.0400)

- Valid classes: Climb (AUC 0.8524, F1 0.0400), Approach (AUC 0.7747, F1 0.0211), Explore (AUC 0.6969, F1 0.4275)

**Code.**

```
_P0 = 1.0
_P1 = 1.0
_P2 = 1.0
_P3 = 1.0
_P4 = 1.0
_P5 = 0.5
_P6 = 1.0
_P7 = 1.0
i_hvx = idx.get('head_vel_x', -1)
i_hvy = idx.get('head_vel_y', -1)
i_bvx = idx.get('body_vel_x', -1)
i_bvy = idx.get('body_vel_y', -1)
i_tvx = idx.get('tail_vel_x', -1)
i_tvy = idx.get('tail_vel_y', -1)
i_speed = idx.get('speed', -1)
i_acc = idx.get('acceleration', -1)
i_compact = idx.get('body_compactness', -1)
if min(i_hvx, i_hvy, i_bvx, i_bvy, i_tvx, i_tvy, i_speed, i_acc, i_compact)
< 0:
    _result = np.full(windows.shape[0], np.nan)
else:
    limb_vel = np.mean(windows[:, :, i_hvx] + windows[:, :, i_hvy] + windows
[:, :, i_bvx] + windows[:, :, i_bvy] + windows[:, :, i_tvx] + windows[:, :,
i_tvy], axis=1)
    speed_mean = np.mean(windows[:, :, i_speed], axis=1)
    speed_var = np.std(windows[:, :, i_speed], axis=1)
    acc_mean = np.mean(windows[:, :, i_acc], axis=1)
    compact_mean = np.mean(windows[:, :, i_compact], axis=1)
    compact_regular = _P0 - np.abs(compact_mean - _P5)
    quiet_score = (_P1 - limb_vel) * (_P2 - speed_mean) * (_P3 - speed_var)
* (_P4 - acc_mean)
    _result = quiet_score * compact_regular ** _P6 * (_P7 - np.std(windows
[:, :, i_compact], axis=1))
    _result = _result.astype(float)
```

#### factor_blank_global_velocity_still_compact_stable (rank 5, AUC 0.8522)

**Description.** Characterizes a four-paw stationary state: own speed, acceleration, and the velocities of the head, body and tail are all low, while body compactness remains stable within the window.

- Designed target: **Blank** · Mode: batch · seq_length: 60 · n_samples: 3595
- Best class: Climb (AUC 0.8522, F1 0.0539)
- Valid classes: Climb (AUC 0.8522, F1 0.0539)

### Code.

```
i_speed = idx.get('speed', -1)
i_acc = idx.get('acceleration', -1)
i_bvx = idx.get('body_vel_x', -1)
i_bvy = idx.get('body_vel_y', -1)
i_hvx = idx.get('head_vel_x', -1)
i_hvy = idx.get('head_vel_y', -1)
i_tvx = idx.get('tail_vel_x', -1)
i_tvy = idx.get('tail_vel_y', -1)
i_comp = idx.get('body_compactness', -1)
if i_speed < 0 or i_acc < 0 or i_bvx < 0 or i_bvy < 0 or i_hvx < 0 or i_hvy
< 0 or i_tvx < 0 or i_tvy < 0 or i_comp < 0:
    _result = np.full(len(windows), np.nan)
else:
    motion = (np.mean(windows[:, :, i_speed], axis=1) + np.mean(windows[:,
 :, i_acc], axis=1) + np.mean(windows[:, :, i_bvx], axis=1) + np.mean(window
s[:, :, i_bvy], axis=1) + np.mean(windows[:, :, i_hvx], axis=1) + np.mean(wi
ndows[:, :, i_hvy], axis=1) + np.mean(windows[:, :, i_tvx], axis=1) + np.mea
n(windows[:, :, i_tvy], axis=1)) / 8.0
    comp_stability = 1.0 - np.std(windows[:, :, i_comp], axis=1)
    comp_mid = 1.0 - np.abs(np.mean(windows[:, :, i_comp], axis=1) - 0.5) *
2.0
    _result = np.clip(1.0 - motion, 0.0, 1.0) * np.clip(comp_stability, 0.0,
 1.0) * np.clip(comp_mid, 0.0, 1.0)
```

#### Self-grooming (class index 2)

#### Supplementary Table 25. Top-five factors selected for Self-grooming.

Factors are ordered by their one-versus-rest validation AUC on the selected class (Self-grooming); the selected AUC is the class-specific AUC used for ranking.

| Rank | Factor name | Selected AUC | Designed target | seq_length |
|---|---|---|---|---|
| 1 | factor_self_grooming_snout_compact_tail_curve_rhythm_llmp | 0.7593 | Self-grooming | 30 |
| 2 | factor_self_grooming_snout_compact_tail_curve_rhythm | 0.7593 | Self-grooming | 30 |
| 3 | factor_climb_vertical_extension_walllike_ | 0.7287 | Climb | 30 |

| Rank | Factor name | Selected AUC | Designed target | seq_length |
|---|---|---|---|---|
| | body_velocity | | | |
| 4 | factor_blank_wholebody_velocity_flat_compact_pose | 0.7283 | Blank | 60 |
| 5 | factor_self_grooming_compact_nose_tail_asocial | 0.7122 | Self-grooming | 60 |

**Supplementary Table 26. One-versus-rest validation AUC of the five leading Self-grooming factors across all seven behavior classes.**

Rows correspond to the ranked factors listed in Supplementary Table 25. Bold values indicate the class-specific AUC on which each factor was selected. Factor descriptions, validation details, and executable code are provided after each table.

| Rank | Explore | Climb | Self-grooming | Stand | Blank | Sniffing | Approach |
|---|---|---|---|---|---|---|---|
| 1 | 0.6848 | 0.5885 | **0.7593** | 0.5752 | 0.5836 | 0.5253 | 0.7371 |
| 2 | 0.6848 | 0.5885 | **0.7593** | 0.5752 | 0.5836 | 0.5253 | 0.7371 |
| 3 | 0.6377 | 0.7198 | **0.7287** | 0.5325 | 0.6424 | 0.5593 | 0.6523 |
| 4 | 0.6941 | 0.7331 | **0.7283** | 0.5556 | 0.6166 | 0.5169 | 0.5841 |
| 5 | 0.6570 | 0.6323 | **0.7122** | 0.5408 | 0.5304 | 0.6464 | 0.6910 |

**factor_self_grooming_snout_compact_tail_curve_rhythm_llmp (rank 1, AUC 0.7593)**

**Description.** Self-grooming factor: combines body compactness, local nose-to-head oscillation, tail-curvature variation and low speed to capture fine rhythmic head and tail activity in a curled posture.

- Designed target: **Self-grooming** · Mode: batch · seq_length: 30 · n_samples: 8353
- Best class: Self-grooming (AUC 0.7593, F1 0.4572)
- Valid classes: Self-grooming (AUC 0.7593, F1 0.4572), Approach (AUC 0.7371, F1 0.0350), Explore (AUC 0.6848, F1 0.4624)

**Code.**

```
_P0 = 0.5   # body_compactness mean weight
_P1 = 1.0   # nose_to_head fluctuation weight
_P2 = 1.0   # tail_curve fluctuation weight
```

```
_P3 = 1.0  # speed inverse weight
_P4 = 1.0  # body_length inverse weight
_P5 = 1.0  # tail_motion weight
_P6 = 1.0  # inverse baseline
_P7 = 1.0  # inverse baseline
i_comp = idx.get('body_compactness', -1)
i_nh = idx.get('nose_to_head', -1)
i_curve = idx.get('tail_curve', -1)
i_speed = idx.get('speed', -1)
i_len = idx.get('body_length', -1)
i_tailm = idx.get('tail_motion', -1)
if min(i_comp, i_nh, i_curve, i_speed, i_len, i_tailm) < 0:
    _result = np.full(windows.shape[0], np.nan)
else:
    compact_m = np.mean(windows[:, :, i_comp], axis=1)
    snout_rhythm = np.std(windows[:, :, i_nh], axis=1)
    curve_rhythm = np.std(windows[:, :, i_curve], axis=1)
    quiet = _P6 - np.mean(windows[:, :, i_speed], axis=1)
    short_body = _P7 - np.mean(windows[:, :, i_len], axis=1)
    tail_local = np.mean(windows[:, :, i_tailm], axis=1)
    _result = _P0 * compact_m + _P1 * snout_rhythm + _P2 * curve_rhythm + _P
3 * quiet + _P4 * short_body + _P5 * tail_local
_result = _result.astype(float)
```

#### factor_self_grooming_snout_compact_tail_curve_rhythm (rank 2, AUC 0.7593)

**Description.** Self-grooming factor: combines body compactness, local nose-to-head oscillation, tail-curvature variation and low speed to capture fine rhythmic head and tail activity in a curled posture.

- Designed target: **Self-grooming** · Mode: batch · seq_length: 30 · n_samples: 8353
- Best class: Self-grooming (AUC 0.7593, F1 0.4572)
- Valid classes: Self-grooming (AUC 0.7593, F1 0.4572), Approach (AUC 0.7371, F1 0.0350), Explore (AUC 0.6848, F1 0.4624)

**Code.**

```
_P0 = 1.0  # body_compactness mean weight
_P1 = 1.0  # nose_to_head fluctuation weight
_P2 = 1.0  # tail_curve fluctuation weight
_P3 = 1.0  # speed inverse weight
_P4 = 1.0  # body_length inverse weight
_P5 = 1.0  # tail_motion weight
_P6 = 1.0  # inverse baseline
_P7 = 1.0  # inverse baseline
i_comp = idx.get('body_compactness', -1)
i_nh = idx.get('nose_to_head', -1)
i_curve = idx.get('tail_curve', -1)
```

```
i_speed = idx.get('speed', -1)
i_len = idx.get('body_length', -1)
i_tailm = idx.get('tail_motion', -1)
if min(i_comp, i_nh, i_curve, i_speed, i_len, i_tailm) < 0:
    _result = np.full(windows.shape[0], np.nan)
else:
    compact_m = np.mean(windows[:, :, i_comp], axis=1)
    snout_rhythm = np.std(windows[:, :, i_nh], axis=1)
    curve_rhythm = np.std(windows[:, :, i_curve], axis=1)
    quiet = _P6 - np.mean(windows[:, :, i_speed], axis=1)
    short_body = _P7 - np.mean(windows[:, :, i_len], axis=1)
    tail_local = np.mean(windows[:, :, i_tailm], axis=1)
    _result = _P0 * compact_m + _P1 * snout_rhythm + _P2 * curve_rhythm + _P
3 * quiet + _P4 * short_body + _P5 * tail_local
_result = _result.astype(float)
```

#### factor_climb_vertical_extension_walllike_body_velocity (rank 3, AUC 0.7287)

**Description.** Climbing factor: the body orientation is near vertical with clear stretching of the body, while body velocity components differ from head velocity components, reflecting the strained posture of wall-adjacent climbing attempts.

- Designed target: **Climb** · Mode: batch · seq_length: 30 · n_samples: 8353
- Best class: Self-grooming (AUC 0.7287, F1 0.4555)
- Valid classes: Self-grooming (AUC 0.7287, F1 0.4555), Climb (AUC 0.7198, F1 0.0498), Approach (AUC 0.6523, F1 0.0253)

**Code.**

```
_P0 = 1.0
_P1 = 1.0
_P2 = 1.0
_P3 = 1.0
_P4 = 0.5
_P5 = 1.0
_P6 = 1.0
i_orient = idx.get('body_orientation', -1)
i_len = idx.get('body_length', -1)
i_hvy = idx.get('head_vel_y', -1)
i_bvy = idx.get('body_vel_y', -1)
i_tail = idx.get('body_to_tail', -1)
i_comp = idx.get('body_compactness', -1)
if min(i_orient, i_len, i_hvy, i_bvy, i_tail, i_comp) < 0:
    _result = np.full(len(windows), np.nan)
else:
    vertical = np.mean(windows[:, :, i_orient], axis=1)
    stretch_range = np.max(windows[:, :, i_len], axis=1) - np.min(windows[:,
```

```
:, i_len], axis=1)
    body_head_mismatch = np.mean(np.abs(windows[:, :, i_bvy] - windows[:, :,
 i_hvy]), axis=1)
    tail_anchor_change = np.std(windows[:, :, i_tail], axis=1)
    compact_low = _P4 - np.mean(windows[:, :, i_comp], axis=1)
    _result = (_P0 * vertical) * (_P1 * stretch_range + _P2 * tail_anchor_ch
ange) * (_P3 * body_head_mismatch) * (compact_low ** _P5) * _P6
```

#### factor_blank_wholebody_velocity_flat_compact_pose (rank 4, AUC 0.7283)

**Description.** Identifies four-paw stillness: head, body and tail velocities are all very low, and body compactness and posture remain stable within the window.

- Designed target: **Blank** · Mode: batch · seq_length: 60 · n_samples: 3595
- Best class: Climb (AUC 0.7331, F1 0.0166)
- Valid classes: Climb (AUC 0.7331, F1 0.0166), Self-grooming (AUC 0.7283, F1 0.4573), Explore (AUC 0.6941, F1 0.4447)

**Code.**

```
_P0 = 1.0
_P1 = 1.0
_P2 = 1.0
_P3 = 1.0
_P4 = 1.0
_P5 = 1.0
_P6 = 1.0
_P7 = 1.0
_P8 = 0.5
i_hx = idx.get('head_vel_x', -1)
i_hy = idx.get('head_vel_y', -1)
i_bx = idx.get('body_vel_x', -1)
i_by = idx.get('body_vel_y', -1)
i_tx = idx.get('tail_vel_x', -1)
i_ty = idx.get('tail_vel_y', -1)
i_compact = idx.get('body_compactness', -1)
i_orient = idx.get('body_orientation', -1)
if i_hx < 0 or i_hy < 0 or i_bx < 0 or i_by < 0 or i_tx < 0 or i_ty < 0 or i
_compact < 0 or i_orient < 0:
    _result = np.full(len(windows), np.nan)
else:
    vel_sum = np.mean(windows[:, :, i_hx], axis=1) + np.mean(windows[:, :, i
_hy], axis=1) + np.mean(windows[:, :, i_bx], axis=1) + np.mean(windows[:, :,
 i_by], axis=1) + np.mean(windows[:, :, i_tx], axis=1) + np.mean(windows[:,
 :, i_ty], axis=1)
    vel_flat = np.std(windows[:, :, i_hx], axis=1) + np.std(windows[:, :, i_
hy], axis=1) + np.std(windows[:, :, i_bx], axis=1) + np.std(windows[:, :, i_
by], axis=1)
    compact_stable = _P2 / (_P2 + np.std(windows[:, :, i_compact], axis=1) *
```

```
* _P3)
    orient_stable = _P4 / (_P4 + np.std(windows[:, :, i_orient], axis=1) **
_P5)
    low_velocity = _P0 / (_P0 + vel_sum ** _P1)
    low_variation = _P6 / (_P6 + vel_flat ** _P7)
    _result = low_velocity * low_variation * compact_stable * orient_stable
    _result = np.where(np.mean(windows[:, :, i_compact], axis=1) > _P8, _res
ult, _result * _P8)
```

#### factor_self_grooming_compact_nose_tail_asocial (rank 5, AUC 0.7122)

**Description.** Self-grooming factor: fluctuating body compactness and rhythmic changes of the nose-to-head distance, with tail curvature participating but overall movement low, while the mouse stays far from the companion without mutual facing.

- Designed target: **Self-grooming** · Mode: batch · seq_length: 60 · n_samples: 3595
- Best class: Self-grooming (AUC 0.7122, F1 0.4471)
- Valid classes: Self-grooming (AUC 0.7122, F1 0.4471), Approach (AUC 0.6910, F1 0.0166), Explore (AUC 0.6570, F1 0.3982)

**Code.**

```
_P0 = 1.0
_P1 = 1.0
_P2 = 1.0
_P3 = 1.0
_P4 = 1.0
_P5 = 0.01
_P6 = 1.0
i_comp = idx.get('body_compactness', -1)
i_nose = idx.get('nose_to_head', -1)
i_curve = idx.get('tail_curve', -1)
i_speed = idx.get('speed', -1)
i_dist = idx.get('dist_to_other', -1)
i_mutual = idx.get('mutual_facing', -1)
i_app = idx.get('approach_speed', -1)
if min(i_comp, i_nose, i_curve, i_speed, i_dist, i_mutual, i_app) < 0:
    _result = np.full(windows.shape[0], np.nan)
else:
    compact_pulse = np.std(windows[:, :, i_comp], axis=1) + np.max(windows
[:, :, i_comp], axis=1) - np.min(windows[:, :, i_comp], axis=1)
    nose_pulse = np.std(windows[:, :, i_nose], axis=1)
    tail_curve = np.mean(windows[:, :, i_curve], axis=1) + np.std(windows[:,
 :, i_curve], axis=1)
    low_move = _P3 / (_P5 + np.mean(windows[:, :, i_speed], axis=1))
    asocial = (_P4 + np.mean(windows[:, :, i_dist], axis=1)) / (_P5 + np.mea
n(windows[:, :, i_mutual], axis=1) + np.mean(windows[:, :, i_app], axis=1))
```

```
    _result = (_P0 * compact_pulse + _P1 * nose_pulse + _P2 * tail_curve) * low_move * (asocial ** _P6)
```

**Stand (class index 3)**

**Supplementary Table 27. Top-five factors selected for Stand.**

Factors are ordered by their one-versus-rest validation AUC on the selected class (Stand); the selected AUC is the class-specific AUC used for ranking.

| Rank | Factor name | Selected AUC | Designed target | seq_length |
|---|---|---|---|---|
| 1 | factor_stand_late_rearup_headbody_accel_damped | 0.6750 | Stand | 60 |
| 2 | factor_stand_vertical_stretch_quiet_head_ratio_plateau_mean2max | 0.6567 | Stand | 15 |
| 3 | factor_stand_upright_head_body_tail_quiet_llmp | 0.6439 | Stand | 15 |
| 4 | factor_stand_forebody_posture_tail_curve_private_stillness_llmp | 0.6426 | Stand | 30 |
| 5 | factor_stand_forebody_posture_tail_curve_private_stillness | 0.6426 | Stand | 30 |

**Supplementary Table 28. One-versus-rest validation AUC of the five leading Stand factors across all seven behavior classes.**

Rows correspond to the ranked factors listed in Supplementary Table 27. Bold values indicate the class-specific AUC on which each factor was selected. Factor descriptions, validation details, and executable code are provided after each table.

| Rank | Explore | Climb | Self-grooming | Stand | Blank | Sniffing | Approach |
|---|---|---|---|---|---|---|---|
| 1 | 0.4787 | 0.7744 | 0.6454 | **0.6750** | 0.5454 | 0.4403 | 0.4878 |
| 2 | 0.5939 | 0.5797 | 0.5524 | **0.6567** | 0.5148 | 0.5546 | 0.6659 |
| 3 | 0.5346 | 0.5417 | 0.5489 | **0.6439** | 0.5044 | 0.6568 | 0.6411 |
| 4 | 0.6647 | 0.6003 | 0.5328 | **0.6426** | 0.4987 | 0.5061 | 0.6650 |
| 5 | 0.6647 | 0.6003 | 0.5328 | **0.6426** | 0.4987 | 0.5061 | 0.6650 |

**factor_stand_late_rearup_headbody_accel_damped (rank 1, AUC 0.6750)**

**Description.** Bipedal standing factor: in the later part of the window the body axis becomes more vertical with a head-body stretching trend, accompanied by low acceleration and low speed, capturing episodes that remain standing after a late rear-up.

- Designed target: **Stand** · Mode: batch · seq_length: 60 · n_samples: 3595
- Best class: Climb (AUC 0.7744, F1 0.0201)
- Valid classes: Climb (AUC 0.7744, F1 0.0201), Stand (AUC 0.6750, F1 0.4279)

**Code.**

```
_P0 = 1.0  # body_orientation trend weight
_P1 = 1.0  # head_to_body trend weight
_P2 = 1.0  # nose_to_head mean weight
_P3 = 1.0  # body_length mean weight
_P4 = 1.0  # speed decay scaling
_P5 = 1.0  # acceleration decay scaling
_P6 = 1.0  # body_vel decay scaling
_P7 = 1.0  # total-score weight
i_ori = idx.get('body_orientation', -1)
i_hb = idx.get('head_to_body', -1)
i_nh = idx.get('nose_to_head', -1)
i_bl = idx.get('body_length', -1)
i_sp = idx.get('speed', -1)
i_acc = idx.get('acceleration', -1)
i_bvx = idx.get('body_vel_x', -1)
i_bvy = idx.get('body_vel_y', -1)
if min([i_ori, i_hb, i_nh, i_bl, i_sp, i_acc, i_bvx, i_bvy]) < 0:
    _result = np.full(windows.shape[0], np.nan)
else:
    ori_trend = windows[:, -1, i_ori] - windows[:, 0, i_ori]
    hb_trend = windows[:, -1, i_hb] - windows[:, 0, i_hb]
    nh_mean = np.mean(windows[:, :, i_nh], axis=1)
    bl_mean = np.mean(windows[:, :, i_bl], axis=1)
    sp_mean = np.mean(windows[:, :, i_sp], axis=1)
    acc_mean = np.mean(windows[:, :, i_acc], axis=1)
    bv_mag = np.sqrt(windows[:, :, i_bvx] * windows[:, :, i_bvx] + windows
```

```
[:, :, i_bvy] * windows[:, :, i_bvy])
    bv_mean = np.mean(bv_mag, axis=1)
    rearup = _P0 * np.maximum(ori_trend, 0.0) + _P1 * np.maximum(hb_trend,
0.0) + _P2 * nh_mean + _P3 * bl_mean
    quiet = _P4 / (_P4 + sp_mean)
    accel_damped = _P5 / (_P5 + acc_mean)
    body_still = _P6 / (_P6 + bv_mean)
    _result = _P7 * rearup * quiet * accel_damped * body_still
```

#### factor_stand_vertical_stretch_quiet_head_ratio_plateau_mean2max (rank 2, AUC 0.6567)

**Description.** Standing factor: persistently near-vertical body orientation and body elongation, with stable changes in the nose-to-head and head-to-body ratios and low overall speed, emphasizing the postural plateau within 15 frames. [np.mean->max]

- Designed target: **Stand** · Mode: batch · seq_length: 15 · n_samples: 18405
- Best class: Approach (AUC 0.6659, F1 0.0389)
- Valid classes: Approach (AUC 0.6659, F1 0.0389), Stand (AUC 0.6567, F1 0.4365)

**Code.**

```
_P0 = 1.0  # body_orientation weight
_P1 = 1.0  # body_length weight
_P2 = 1.0  # nose_to_head relative weight
_P3 = 1.0  # head_to_body relative weight
_P4 = 1.0  # stillness penalty weight
_P5 = 1.0  # posture-stability penalty weight
_P6 = 0.01  # division stability term
_P7 = 1.0  # nonlinear exponent

i_orient = idx.get('body_orientation', -1)
i_len = idx.get('body_length', -1)
i_nh = idx.get('nose_to_head', -1)
i_hb = idx.get('head_to_body', -1)
i_speed = idx.get('speed', -1)
if i_orient < 0 or i_len < 0 or i_nh < 0 or i_hb < 0 or i_speed < 0:
    _result = np.full(len(windows), np.nan)
else:
    orient_mean = np.max(windows[:, :, i_orient], axis=1)
    orient_std = np.std(windows[:, :, i_orient], axis=1)
    len_mean = np.max(windows[:, :, i_len], axis=1)
    nh_mean = np.max(windows[:, :, i_nh], axis=1)
    hb_mean = np.max(windows[:, :, i_hb], axis=1)
    speed_mean = np.max(windows[:, :, i_speed], axis=1)
    ratio = (_P2 * nh_mean + _P6) / (_P3 * hb_mean + _P6)
    ratio_stability = np.std((windows[:, :, i_nh] + _P6) / (windows[:, :, i_
```

```
hb] + _P6), axis=1)
    posture = _P0 * orient_mean + _P1 * len_mean + ratio
    quiet = np.power(np.maximum(_P6, 1.0 - _P4 * speed_mean), _P7)
    stable = np.maximum(_P6, 1.0 - _P5 * (orient_std + ratio_stability))
    _result = posture * quiet * stable
```

#### factor_stand_upright_head_body_tail_quiet_llmp (rank 3, AUC 0.6439)

**Description.** Bipedal standing factor: combines near-vertical body orientation, head-to-body ratio changes, low tail motion, and low overall speed over time to capture the mouse's stationary standing posture.

- Designed target: **Stand** · Mode: batch · seq_length: 15 · n_samples: 18405
- Best class: Sniffing (AUC 0.6568, F1 0.1291)
- Valid classes: Sniffing (AUC 0.6568, F1 0.1291)

**Code.**

```
_P0 = 2.0
_P1 = 0.35
_P2 = 0.05
_P3 = 1.5
_P4 = 5.0
_P5 = 1.5
i_orient = idx.get('body_orientation', -1)
i_nose = idx.get('nose_to_head', -1)
i_head = idx.get('head_to_body', -1)
i_tail = idx.get('tail_motion', -1)
i_speed = idx.get('speed', -1)
if i_orient < 0 or i_nose < 0 or i_head < 0 or i_tail < 0 or i_speed < 0:
    _result = np.full(len(windows), np.nan)
else:
    orient_mean = np.mean(windows[:, :, i_orient], axis=1)
    nose_mean = np.mean(windows[:, :, i_nose], axis=1)
    head_mean = np.mean(windows[:, :, i_head], axis=1)
    tail_mean = np.mean(windows[:, :, i_tail], axis=1)
    speed_mean = np.mean(windows[:, :, i_speed], axis=1)
    head_body_contrast = _P0 * head_mean - _P1 * nose_mean
    quiet = _P2 - _P3 * speed_mean
    tail_anchor = _P4 - _P5 * tail_mean
    _result = orient_mean * head_body_contrast * quiet * tail_anchor
```

#### factor_stand_forebody_posture_tail_curve_private_stillness_llmp (rank 4, AUC 0.6426)

**Description.** Standing factor: forebody proportion changes, vertical body posture and a low-speed state co-occur, combined with the mouse's own tail-curvature fluctuation, capturing the short pattern of forebody elevation with tail-assisted balance during standing.

- Designed target: **Stand** · Mode: batch · seq_length: 30 · n_samples: 8353
- Best class: Approach (AUC 0.6650, F1 0.0235)
- Valid classes: Approach (AUC 0.6650, F1 0.0235), Explore (AUC 0.6647, F1 0.4512)

**Code.**

```
_P0 = 1.0  # nose_to_head mean weight
_P1 = 1.0  # head_to_body mean weight
_P2 = 1.0  # forebody dynamic-range weight
_P3 = 1.0  # body-orientation weight
_P4 = 0.05  # stillness baseline threshold
_P5 = 1.0  # speed penalty weight
_P6 = 1.0  # tail-curvature dynamic weight
i_nose = idx.get('nose_to_head', -1)
i_head = idx.get('head_to_body', -1)
i_orient = idx.get('body_orientation', -1)
i_speed = idx.get('speed', -1)
i_tail_curve = idx.get('tail_curve', -1)
if i_nose < 0 or i_head < 0 or i_orient < 0 or i_speed < 0 or i_tail_curve <
 0:
    _result = np.full(len(windows), np.nan)
else:
    fore = _P0 * windows[:, :, i_nose] + _P1 * windows[:, :, i_head]
    fore_mean = np.mean(fore, axis=1)
    fore_range = np.max(fore, axis=1) - np.min(fore, axis=1)
    vertical = _P3 * np.mean(windows[:, :, i_orient], axis=1)
    quiet = _P4 - _P5 * np.mean(windows[:, :, i_speed], axis=1)
    tail_curve_dyn = _P6 * np.std(windows[:, :, i_tail_curve], axis=1)
    _result = vertical * quiet * (fore_mean + _P2 * fore_range) * tail_curve
_dyn
    _result = _result.astype(float)
```

#### factor_stand_forebody_posture_tail_curve_private_stillness (rank 5, AUC 0.6426)

**Description.** Standing factor: forebody ratio changes, vertical body posture, and low-speed states co-occur, combined with the own tail-curvature fluctuation, capturing the short-lived pattern of raised forebody with tail-assisted balance during standing.

- Designed target: **Stand** · Mode: batch · seq_length: 30 · n_samples: 8353
- Best class: Approach (AUC 0.6650, F1 0.0235)
- Valid classes: Approach (AUC 0.6650, F1 0.0235), Explore (AUC 0.6647, F1 0.4512)

**Code.**

```
_P0 = 1.0  # nose_to_head mean weight
_P1 = 1.0  # head_to_body mean weight
_P2 = 1.0  # forebody dynamic-range weight
_P3 = 1.0  # body-orientation weight
_P4 = 1.0  # stillness baseline threshold
_P5 = 1.0  # speed penalty weight
_P6 = 1.0  # tail-curvature dynamic weight
i_nose = idx.get('nose_to_head', -1)
i_head = idx.get('head_to_body', -1)
i_orient = idx.get('body_orientation', -1)
i_speed = idx.get('speed', -1)
i_tail_curve = idx.get('tail_curve', -1)
if i_nose < 0 or i_head < 0 or i_orient < 0 or i_speed < 0 or i_tail_curve <
 0:
    _result = np.full(len(windows), np.nan)
else:
    fore = _P0 * windows[:, :, i_nose] + _P1 * windows[:, :, i_head]
    fore_mean = np.mean(fore, axis=1)
    fore_range = np.max(fore, axis=1) - np.min(fore, axis=1)
    vertical = _P3 * np.mean(windows[:, :, i_orient], axis=1)
    quiet = _P4 - _P5 * np.mean(windows[:, :, i_speed], axis=1)
    tail_curve_dyn = _P6 * np.std(windows[:, :, i_tail_curve], axis=1)
    _result = vertical * quiet * (fore_mean + _P2 * fore_range) * tail_curve
_dyn
    _result = _result.astype(float)
```

### Blank (class index 4)

**Supplementary Table 29. Top-five factors selected for Blank.**

Factors are ordered by their one-versus-rest validation AUC on the selected class (Blank); the selected AUC is the class-specific AUC used for ranking.

| Rank | Factor name | Selected AUC | Designed target | seq_length |
|---|---|---|---|---|
| 1 | factor_positive_sniffs_close_mutual_headmicro_tailquiet | 0.7443 | Sniffing | 60 |
| 2 | factor_climb_vertical_bodyy_lengthrange_tailangle_lowapproach | 0.7362 | Climb | 60 |
| 3 | factor_climb_vertical_tailbr | 0.7309 | Climb | 60 |

| Rank | Factor name | Selected AUC | Designed target | seq_length |
|---|---|---|---|---|
| | ace_extension_pulses | | | |
| 4 | factor_self_grooming_head_body_decouple_tailcurve_pause | 0.7278 | Self-grooming | 60 |
| 5 | factor_climb_vertical_length_body_velocity_asymmetry | 0.7253 | Climb | 60 |

**Supplementary Table 30. One-versus-rest validation AUC of the five leading Blank factors across all seven behavior classes.**

Rows correspond to the ranked factors listed in Supplementary Table 29. Bold values indicate the class-specific AUC on which each factor was selected. Factor descriptions, validation details, and executable code are provided after each table.

| Rank | Explore | Climb | Self-grooming | Stand | Blank | Sniffing | Approach |
|---|---|---|---|---|---|---|---|
| 1 | 0.6114 | 0.5501 | 0.6455 | 0.4833 | **0.7443** | 0.6620 | 0.6487 |
| 2 | 0.6690 | 0.6266 | 0.6062 | 0.6088 | **0.7362** | 0.5055 | 0.2893 |
| 3 | 0.6683 | 0.5368 | 0.5897 | 0.6120 | **0.7309** | 0.5107 | 0.2770 |
| 4 | 0.7150 | 0.7506 | 0.6102 | 0.5706 | **0.7278** | 0.6835 | 0.7123 |
| 5 | 0.7173 | 0.5839 | 0.6621 | 0.5612 | **0.7253** | 0.5374 | 0.7464 |

**factor_positive_sniffs_close_mutual_headmicro_tailquiet (rank 1, AUC 0.7443)**

**Description.** Identifies positive sniffing: short distance and mutual facing between the pair, fine head activity of the subject with low overall speed and low tail motion.

- Designed target: **Sniffing** · Mode: batch · seq_length: 60 · n_samples: 3595
- Best class: Blank (AUC 0.7443, F1 0.5428)
- Valid classes: Blank (AUC 0.7443, F1 0.5428), Sniffing (AUC 0.6620, F1 0.1160)

**Code.**

```
_P0 = 1.0  # close-distance inverse weight
_P1 = 1.0  # mutual_facing weight
_P2 = 1.0  # self_facing_other weight
_P3 = 1.0  # head_vel_x micro-motion weight
_P4 = 1.0  # head_vel_y micro-motion weight
_P5 = 1.0  # low-speed weight
_P6 = 1.0  # low tail-motion weight
_P7 = 1.0  # baseline inverse term
i_dist = idx.get('dist_to_other', -1)
i_mutual = idx.get('mutual_facing', -1)
i_face = idx.get('self_facing_other', -1)
i_hvx = idx.get('head_vel_x', -1)
i_hvy = idx.get('head_vel_y', -1)
i_speed = idx.get('speed', -1)
i_tail = idx.get('tail_motion', -1)
if min(i_dist, i_mutual, i_face, i_hvx, i_hvy, i_speed, i_tail) < 0:
    _result = np.full(windows.shape[0], np.nan)
else:
    dist = windows[:, :, i_dist]
    mutual = windows[:, :, i_mutual]
    face = windows[:, :, i_face]
    hvx = windows[:, :, i_hvx]
    hvy = windows[:, :, i_hvy]
    speed = windows[:, :, i_speed]
    tail = windows[:, :, i_tail]
    close = _P7 - np.mean(dist, axis=1)
    social_focus = _P1 * np.mean(mutual, axis=1) + _P2 * np.mean(face, axis=
1)
    head_micro = _P3 * np.std(hvx, axis=1) + _P4 * np.std(hvy, axis=1)
    pause = _P5 * (_P7 - np.mean(speed, axis=1)) + _P6 * (_P7 - np.mean(tai
l, axis=1))
    _result = (_P0 * close) * social_focus * head_micro * pause
```

#### factor_climb_vertical_bodyy_lengthrange_tailangle_lowapproach (rank 2, AUC 0.7362)

**Description.** Climbing factor: uses near-vertical body orientation, body-length change, body-y velocity fluctuation and tail-angle change to describe wall-climbing attempts, while excluding social movement toward the companion.

- Designed target: **Climb** · Mode: batch · seq_length: 60 · n_samples: 3595
- Best class: Blank (AUC 0.7362, F1 0.5781)
- Valid classes: Blank (AUC 0.7362, F1 0.5781), Explore (AUC 0.6690, F1 0.3937)

**Code.**

```
_P0 = 1.0  # body_orientation weight
_P1 = 1.0  # body_length range weight
```

```
_P2 = 1.0  # body_vel_y fluctuation weight
_P3 = 1.0  # tail_angle range weight
_P4 = 1.0  # approach_speed low social-interaction weight
_P5 = 1.0  # speed low-displacement weight
_P6 = 1.0  # orientation fluctuation-penalty scaling
_P7 = 1.0  # approach_speed decay scaling
_P8 = 1.0  # speed decay scaling
i_ori = idx.get('body_orientation', -1)
i_len = idx.get('body_length', -1)
i_by = idx.get('body_vel_y', -1)
i_ta = idx.get('tail_angle', -1)
i_app = idx.get('approach_speed', -1)
i_speed = idx.get('speed', -1)
if i_ori < 0 or i_len < 0 or i_by < 0 or i_ta < 0 or i_app < 0 or i_speed <
0:
    _result = np.full(len(windows), np.nan)
else:
    ori_mean = np.mean(windows[:, :, i_ori], axis=1)
    ori_std = np.std(windows[:, :, i_ori], axis=1)
    len_range = np.max(windows[:, :, i_len], axis=1) - np.min(windows[:, :,
i_len], axis=1)
    by_std = np.std(windows[:, :, i_by], axis=1)
    by_range = np.max(windows[:, :, i_by], axis=1) - np.min(windows[:, :, i_
by], axis=1)
    ta_range = np.max(windows[:, :, i_ta], axis=1) - np.min(windows[:, :, i_
ta], axis=1)
    app_mean = np.mean(windows[:, :, i_app], axis=1)
    speed_mean = np.mean(windows[:, :, i_speed], axis=1)
    vertical = _P0 * ori_mean / (_P0 + _P6 * ori_std)
    stretch_change = _P1 * len_range
    wall_drive = _P2 * (by_std + by_range)
    tail_adjust = _P3 * ta_range
    non_social = _P4 / (_P0 + _P7 * app_mean)
    low_translation = _P5 / (_P0 + _P8 * speed_mean)
    _result = vertical * stretch_change * wall_drive * tail_adjust * non_soc
ial * low_translation
```

#### factor_climb_vertical_tailbrace_extension_pulses (rank 3, AUC 0.7309)

**Description.** Identifies suspected climbing: the body tends vertical, body length/stretching changes, the tail angle or curvature participates in bracing, and body velocity appears as local pulses rather than translational running.

- Designed target: **Climb** · Mode: batch · seq_length: 60 · n_samples: 3595
- Best class: Blank (AUC 0.7309, F1 0.5681)
- Valid classes: Blank (AUC 0.7309, F1 0.5681), Explore (AUC 0.6683, F1 0.4322)

**Code.**

```
_P0 = 1.0
_P1 = 1.0
_P2 = 1.0
_P3 = 1.0
_P4 = 1.0
_P5 = 1.0
_P6 = 0.5
_P7 = 1.0
i_orient = idx.get('body_orientation', -1)
i_len = idx.get('body_length', -1)
i_bvy = idx.get('body_vel_y', -1)
i_bvx = idx.get('body_vel_x', -1)
i_tail_angle = idx.get('tail_angle', -1)
i_tail_curve = idx.get('tail_curve', -1)
if i_orient < 0 or i_len < 0 or i_bvy < 0 or i_bvx < 0 or i_tail_angle < 0 o
r i_tail_curve < 0:
    _result = np.full(len(windows), np.nan)
else:
    orient_mean = np.mean(windows[:, :, i_orient], axis=1)
    orient_stable = _P5 - np.std(windows[:, :, i_orient], axis=1)
    len_range = np.max(windows[:, :, i_len], axis=1) - np.min(windows[:, :,
i_len], axis=1)
    len_late_gain = windows[:, -1, i_len] - windows[:, 0, i_len]
    vertical_pulse = np.std(windows[:, :, i_bvy], axis=1)
    lateral_quiet = _P4 - np.mean(windows[:, :, i_bvx], axis=1)
    tail_brace = _P2 * np.std(windows[:, :, i_tail_angle], axis=1) + _P3 * n
p.mean(windows[:, :, i_tail_curve], axis=1)
    extension = _P1 * len_range + len_late_gain
    score = _P0 * orient_mean * orient_stable * extension * vertical_pulse *
 lateral_quiet * tail_brace
    _result = np.where(orient_mean > _P6, score * _P7, score)
```

#### factor_self_grooming_head_body_decouple_tailcurve_pause (rank 4, AUC 0.7278)

**Description.** Self-grooming factor: during a low-speed pause, head velocity fluctuates more than body velocity, accompanied by tail-curvature and body-compactness changes, emphasizing local head movements rather than whole-body displacement.

- Designed target: **Self-grooming** · Mode: batch · seq_length: 60 · n_samples: 3595
- Best class: Climb (AUC 0.7506, F1 0.0191)
- Valid classes: Climb (AUC 0.7506, F1 0.0191), Blank (AUC 0.7278, F1 0.5713), Explore (AUC 0.7150, F1 0.4425), Approach (AUC 0.7123, F1 0.0170), Sniffing (AUC 0.6835, F1 0.1032)

**Code.**

```
_P0 = 1.0  # head_vel_x fluctuation weight
_P1 = 1.0  # head_vel_y fluctuation weight
_P2 = 1.0  # body_vel_x fluctuation-suppression scaling
_P3 = 1.0  # body_vel_y fluctuation-suppression scaling
_P4 = 1.0  # tail_curve fluctuation weight
_P5 = 1.0  # body_compactness fluctuation weight
_P6 = 1.0  # speed suppression scaling
_P7 = 1.0  # dist_to_other weight
_P8 = 1.0  # other_facing_self suppression scaling
i_hvx = idx.get('head_vel_x', -1)
i_hvy = idx.get('head_vel_y', -1)
i_bvx = idx.get('body_vel_x', -1)
i_bvy = idx.get('body_vel_y', -1)
i_tc = idx.get('tail_curve', -1)
i_bc = idx.get('body_compactness', -1)
i_sp = idx.get('speed', -1)
i_dist = idx.get('dist_to_other', -1)
i_of = idx.get('other_facing_self', -1)
if min(i_hvx, i_hvy, i_bvx, i_bvy, i_tc, i_bc, i_sp, i_dist, i_of) < 0:
    _result = np.full(windows.shape[0], np.nan)
else:
    hvx = windows[:, :, i_hvx]
    hvy = windows[:, :, i_hvy]
    bvx = windows[:, :, i_bvx]
    bvy = windows[:, :, i_bvy]
    tc = windows[:, :, i_tc]
    bc = windows[:, :, i_bc]
    sp = windows[:, :, i_sp]
    dist = windows[:, :, i_dist]
    of = windows[:, :, i_of]
    head_micro = _P0 * np.std(hvx, axis=1) + _P1 * np.std(hvy, axis=1)
    body_still = (_P2 / (_P2 + np.std(bvx, axis=1))) * (_P3 / (_P3 + np.std
(bvy, axis=1)))
    grooming_posture = _P4 * np.std(tc, axis=1) + _P5 * np.std(bc, axis=1)
    locomotion_pause = _P6 / (_P6 + np.mean(sp, axis=1))
    social_detached = (_P7 * np.mean(dist, axis=1)) * (_P8 / (_P8 + np.mean
(of, axis=1)))
    _result = head_micro * body_still * grooming_posture * locomotion_pause
* social_detached
```

#### factor_climb_vertical_length_body_velocity_asymmetry (rank 5, AUC 0.7253)

**Description.** Climbing factor: near-vertical body orientation with body-length stretching, moderate overall speed, and asymmetric body-velocity changes along x/y, matching the posture transitions of wall-climbing attempts.

- Designed target: **Climb** · Mode: batch · seq_length: 60 · n_samples: 3595
- Best class: Approach (AUC 0.7464, F1 0.0173)

- Valid classes: Approach (AUC 0.7464, F1 0.0173), Blank (AUC 0.7253, F1 0.5657), Explore (AUC 0.7173, F1 0.4581), Self-grooming (AUC 0.6621, F1 0.4342)

**Code.**

```
io = idx.get('body_orientation', -1)
ibl = idx.get('body_length', -1)
ispeed = idx.get('speed', -1)
ibvx = idx.get('body_vel_x', -1)
ibvy = idx.get('body_vel_y', -1)
ita = idx.get('tail_angle', -1)
if min(io, ibl, ispeed, ibvx, ibvy, ita) < 0:
    _result = np.full(len(windows), np.nan)
else:
    orient = windows[:, :, io]
    blen = windows[:, :, ibl]
    speed = windows[:, :, ispeed]
    bvx = windows[:, :, ibvx]
    bvy = windows[:, :, ibvy]
    tail_angle = windows[:, :, ita]
    vertical = np.mean(orient, axis=1)
    extension_change = np.max(blen, axis=1) - np.min(blen, axis=1)
    low_translation = 1.0 - np.mean(speed, axis=1)
    vel_asym = np.abs(np.mean(bvx[:, -20:], axis=1) - np.mean(bvx[:, :20], a
xis=1)) + np.abs(np.mean(bvy[:, -20:], axis=1) - np.mean(bvy[:, :20], axis=
1))
    tail_adjust = np.std(tail_angle, axis=1)
    vertical_persistence = 1.0 - np.std(orient, axis=1)
    _result = vertical * extension_change * np.clip(low_translation, 0, 1) *
 vel_asym * (0.5 + tail_adjust) * np.clip(vertical_persistence, 0, 1)
```

#### Sniffing (class index 5)

**Supplementary Table 31. Top-five factors selected for Sniffing.**

Factors are ordered by their one-versus-rest validation AUC on the selected class (Sniffing); the selected AUC is the class-specific AUC used for ranking.

| Rank | Factor name | Selected AUC | Designed target | seq_length |
|---|---|---|---|---|
| 1 | factor_self_grooming_compact_headtail_micro_asocial_still | 0.8532 | Self-grooming | 60 |
| 2 | factor_approach_face_closing_other_alig | 0.8499 | Approach | 15 |

| Rank | Factor name | Selected AUC | Designed target | seq_length |
|---|---|---|---|---|
| | nment_trend_llmp | | | |
| 3 | factor_approach_facing_closure_body_drive_trend | 0.8480 | Approach | 5 |
| 4 | factor_positive_sniffs_close_nose_face_partner_still | 0.8477 | Sniffing | 30 |
| 5 | factor_self_grooming_compact_head_tail_ripple_private | 0.8442 | Self-grooming | 30 |

**Supplementary Table 32. One-versus-rest validation AUC of the five leading Sniffing factors across all seven behavior classes.**

Rows correspond to the ranked factors listed in Supplementary Table 31. Bold values indicate the class-specific AUC on which each factor was selected. Factor descriptions, validation details, and executable code are provided after each table.

| Rank | Explore | Climb | Self-grooming | Stand | Blank | Sniffing | Approach |
|---|---|---|---|---|---|---|---|
| 1 | 0.6765 | 0.7122 | 0.4810 | 0.4908 | 0.5801 | **0.8532** | 0.6159 |
| 2 | 0.6682 | 0.6018 | 0.4995 | 0.5531 | 0.6499 | **0.8499** | 0.7891 |
| 3 | 0.6309 | 0.6186 | 0.5571 | 0.5539 | 0.6108 | **0.8480** | 0.6437 |
| 4 | 0.6420 | 0.6644 | 0.5422 | 0.5534 | 0.5644 | **0.8477** | 0.6046 |
| 5 | 0.6682 | 0.6604 | 0.5548 | 0.4781 | 0.5240 | **0.8442** | 0.7242 |

**factor_self_grooming_compact_headtail_micro_asocial_still (rank 1, AUC 0.8532)**

**Description.** Self-grooming factor: under low overall movement, body curling/compactness fluctuation, nose-head micro-motion and tail-curvature fluctuation co-occur, and the mouse stays far from the companion.

- Designed target: **Self-grooming** · Mode: batch · seq_length: 60 · n_samples: 3595
- Best class: Sniffing (AUC 0.8532, F1 0.1869)

- Valid classes: Sniffing (AUC 0.8532, F1 0.1869), Climb (AUC 0.7122, F1 0.0191), Explore (AUC 0.6765, F1 0.4339)

**Code.**

```
_P0 = 1.0
_P1 = 1.0
_P2 = 1.0
_P3 = 1.0
_P4 = 1.0
_P5 = 1.0
_P6 = 1.0
_P7 = 1.0
i_compact = idx.get('body_compactness', -1)
i_nose = idx.get('nose_to_head', -1)
i_tailcurve = idx.get('tail_curve', -1)
i_speed = idx.get('speed', -1)
i_dist = idx.get('dist_to_other', -1)
i_acc = idx.get('acceleration', -1)
if i_compact < 0 or i_nose < 0 or i_tailcurve < 0 or i_speed < 0 or i_dist <
 0 or i_acc < 0:
    _result = np.full(len(windows), np.nan)
else:
    compact_std = np.std(windows[:, :, i_compact], axis=1)
    compact_mean = np.mean(windows[:, :, i_compact], axis=1)
    nose_std = np.std(windows[:, :, i_nose], axis=1)
    tailcurve_std = np.std(windows[:, :, i_tailcurve], axis=1)
    speed_mean = np.mean(windows[:, :, i_speed], axis=1)
    acc_mean = np.mean(windows[:, :, i_acc], axis=1)
    dist_mean = np.mean(windows[:, :, i_dist], axis=1)
    micro_activity = _P0 * compact_std + _P1 * nose_std + _P2 * tailcurve_st
d + _P3 * compact_mean
    stillness = _P4 - _P5 * speed_mean - _P6 * acc_mean
    _result = micro_activity * stillness * (_P7 * dist_mean)
```

#### factor_approach_face_closing_other_alignment_trend_llmp (rank 2, AUC 0.8499)

**Description.** Approach factor: jointly uses facing the companion, approach speed, a decreasing distance trend, mutual body-axis alignment and the partner's motion state to characterize active approach from afar.

- Designed target: **Approach** · Mode: batch · seq_length: 15 · n_samples: 18405
- Best class: Sniffing (AUC 0.8499, F1 0.2035)
- Valid classes: Sniffing (AUC 0.8499, F1 0.2035), Approach (AUC 0.7891, F1 0.0609), Explore (AUC 0.6682, F1 0.4747)

**Code.**

```
_P0 = 1.0
_P1 = 1.0
_P2 = 1.0
_P3 = 1.0
_P4 = 1.0
_P5 = 0.35
_P6 = 1.0
_P7 = 1.0
_P8 = 1.0
i_face = idx.get('self_facing_other', -1)
i_app = idx.get('approach_speed', -1)
i_dist = idx.get('dist_to_other', -1)
i_align = idx.get('body_axis_align', -1)
i_ospeed = idx.get('other_speed', -1)
i_hdiff = idx.get('heading_diff', -1)
if i_face < 0 or i_app < 0 or i_dist < 0 or i_align < 0 or i_ospeed < 0 or i
_hdiff < 0:
    _result = np.full(len(windows), np.nan)
else:
    face_mean = np.mean(windows[:, :, i_face], axis=1)
    app_mean = np.mean(windows[:, :, i_app], axis=1)
    align_mean = np.mean(windows[:, :, i_align], axis=1)
    ospeed_mean = np.mean(windows[:, :, i_ospeed], axis=1)
    hdiff_std = np.std(windows[:, :, i_hdiff], axis=1)
    dist_drop = windows[:, 0, i_dist] - windows[:, -1, i_dist]
    dist_mean = np.mean(windows[:, :, i_dist], axis=1)
    directed_close = (_P0 * face_mean) * (_P1 * app_mean) * (_P2 * dist_dro
p)
    partner_context = _P3 * align_mean + _P4 * ospeed_mean
    stable_heading = _P5 - hdiff_std
    _result = _P6 * directed_close * partner_context * stable_heading + _P7
 * app_mean * (_P8 * dist_mean)
```

#### factor_approach_facing_closure_body_drive_trend (rank 3, AUC 0.8480)

**Description.** Approach factor: jointly uses facing the companion, approach speed, body-motion intensity, and a rising approach-speed trend within 5 frames to characterize active approach toward the companion.

- Designed target: **Approach** · Mode: batch · seq_length: 5 · n_samples: 59412
- Best class: Sniffing (AUC 0.8480, F1 0.2080)
- Valid classes: Sniffing (AUC 0.8480, F1 0.2080)

**Code.**

```
_P0 = 1.0
_P1 = 1.0
_P2 = 1.0
_P3 = 1.0
_P4 = 1.0
```

```
_P5 = 1.0
_P6 = 1.0
_P7 = 1.0

i_face = idx.get('self_facing_other', -1)
i_app = idx.get('approach_speed', -1)
i_speed = idx.get('speed', -1)
i_body_vx = idx.get('body_vel_x', -1)
i_body_vy = idx.get('body_vel_y', -1)
i_dist = idx.get('dist_to_other', -1)
i_head = idx.get('heading_diff', -1)
if min(i_face, i_app, i_speed, i_body_vx, i_body_vy, i_dist, i_head) < 0:
    _result = np.full(windows.shape[0], np.nan)
else:
    face = np.mean(windows[:, :, i_face], axis=1)
    closure = np.mean(windows[:, :, i_app], axis=1)
    closure_trend = np.mean(np.diff(windows[:, :, i_app], axis=1), axis=1)
    self_drive = np.mean(windows[:, :, i_speed], axis=1) + _P0 * np.mean(np.
abs(windows[:, :, i_body_vx]) + np.abs(windows[:, :, i_body_vy]), axis=1)
    distance_not_close = np.mean(windows[:, :, i_dist], axis=1)
    heading_consistency = _P1 - np.std(windows[:, :, i_head], axis=1)
    _result = (_P2 * face + _P3 * closure + _P4 * closure_trend) * (_P5 * se
lf_drive) * (_P6 * distance_not_close) * (_P7 * heading_consistency)
```

#### factor_positive_sniffs_close_nose_face_partner_still (rank 4, AUC 0.8477)

**Description.** Positive-sniffing factor: combines short distance, facing the companion, fine nose-segment activity, low own speed and low partner speed, using distance stability to distinguish close sniffing from fast chasing.

- Designed target: **Sniffing** · Mode: batch · seq_length: 30 · n_samples: 8353
- Best class: Sniffing (AUC 0.8477, F1 0.2151)
- Valid classes: Sniffing (AUC 0.8477, F1 0.2151), Climb (AUC 0.6644, F1 0.0302)

**Code.**

```
_P0 = 0.5
_P1 = 1.0
_P2 = 1.0
_P3 = 0.5
_P4 = 0.5
_P5 = 1.0
_P6 = 0.5
_P7 = 1.0
_P8 = 1.0
i_dist = idx.get('dist_to_other', -1)
i_face = idx.get('self_facing_other', -1)
```

```
i_nh = idx.get('nose_to_head', -1)
i_sp = idx.get('speed', -1)
i_osp = idx.get('other_speed', -1)
if i_dist < 0 or i_face < 0 or i_nh < 0 or i_sp < 0 or i_osp < 0:
    _result = np.full(len(windows), np.nan)
else:
    close = np.maximum(_P0 - np.mean(windows[:, :, i_dist], axis=1), 0.0)
    face = np.mean(windows[:, :, i_face], axis=1)
    nose_micro = np.std(windows[:, :, i_nh], axis=1)
    self_quiet = np.maximum(_P3 - np.mean(windows[:, :, i_sp], axis=1), 0.0)
    other_quiet = np.maximum(_P4 - np.mean(windows[:, :, i_osp], axis=1), 0.
0)
    dist_stable = np.maximum(_P6 - np.std(windows[:, :, i_dist], axis=1), 0.
0)
    score = (_P1 * close) * (_P2 * face + _P5 * nose_micro) * (self_quiet +
other_quiet + _P7 * dist_stable)
    _result = score ** _P8
```

#### factor_self_grooming_compact_head_tail_ripple_private (rank 5, AUC 0.8442)

**Description.** Self-grooming factor: combines body compactness, local nose-to-head segment changes, small tail-curvature/tail-motion ripples, and distance from the companion, using window standard deviations to capture fine-grained in-place self-grooming.

- Designed target: **Self-grooming** · Mode: batch · seq_length: 30 · n_samples: 8353
- Best class: Sniffing (AUC 0.8442, F1 0.1908)
- Valid classes: Sniffing (AUC 0.8442, F1 0.1908), Approach (AUC 0.7242, F1 0.0289), Explore (AUC 0.6682, F1 0.4557), Climb (AUC 0.6604, F1 0.0311)

**Code.**

```
_P0 = 1.0
_P1 = 1.0
_P2 = 1.0
_P3 = 0.5
_P4 = 1.0
_P5 = 1.0
_P6 = 0.5
_P7 = 1.0
_P8 = 1.0
i_bc = idx.get('body_compactness', -1)
i_nh = idx.get('nose_to_head', -1)
i_tc = idx.get('tail_curve', -1)
i_tm = idx.get('tail_motion', -1)
i_sp = idx.get('speed', -1)
i_dist = idx.get('dist_to_other', -1)
```

```
if i_bc < 0 or i_nh < 0 or i_tc < 0 or i_tm < 0 or i_sp < 0 or i_dist < 0:
    _result = np.full(len(windows), np.nan)
else:
    compact_m = np.mean(windows[:, :, i_bc], axis=1)
    head_ripple = np.std(windows[:, :, i_nh], axis=1)
    tail_curve_ripple = np.std(windows[:, :, i_tc], axis=1)
    tail_motion_m = np.mean(windows[:, :, i_tm], axis=1)
    still = np.maximum(_P3 - np.mean(windows[:, :, i_sp], axis=1), 0.0)
    private = np.mean(windows[:, :, i_dist], axis=1)
    tail_moderate = np.maximum(_P6 - tail_motion_m, 0.0) + _P7 * tail_curve_
ripple
    score = (_P0 * compact_m + _P1 * head_ripple + _P2 * tail_moderate) * (_
P4 * still) * (_P5 * private)
    _result = score ** _P8
```

#### Approach (class index 6)

#### Supplementary Table 33. Top-five factors selected for Approach.

Factors are ordered by their one-versus-rest validation AUC on the selected class (Approach); the selected AUC is the class-specific AUC used for ranking.

| Rank | Factor name | Selected AUC | Designed target | seq_length |
|---|---|---|---|---|
| 1 | factor_approach_speed_facing_distdrop_axisalign | 0.8389 | Approach | 60 |
| 2 | factor_approach_facing_closure_motion_alignment_llmp | 0.8321 | Approach | 15 |
| 3 | factor_approach_mutual_axis_closure_speed_ramp | 0.8302 | Approach | 15 |
| 4 | factor_approach_facing_closure_speed_tailcalm_consistency | 0.8276 | Approach | 60 |
| 5 | factor_explore_object_speed_heading_jitter_social_far | 0.8267 | Explore | 60 |

**Supplementary Table 34. One-versus-rest validation AUC of the five leading Approach factors across all seven behavior classes.**

Rows correspond to the ranked factors listed in Supplementary Table 33. Bold values indicate the class-specific AUC on which each factor was selected. Factor descriptions, validation details, and executable code are provided after each table.

| Rank | Explore | Climb | Self-grooming | Stand | Blank | Sniffing | Approach |
|---|---|---|---|---|---|---|---|
| 1 | 0.6903 | 0.5643 | 0.5703 | 0.5559 | 0.6080 | 0.5496 | **0.8389** |
| 2 | 0.7042 | 0.6273 | 0.6157 | 0.5408 | 0.6304 | 0.5064 | **0.8321** |
| 3 | 0.7091 | 0.6256 | 0.6057 | 0.5177 | 0.6298 | 0.4925 | **0.8302** |
| 4 | 0.6921 | 0.6287 | 0.5800 | 0.5575 | 0.6040 | 0.5223 | **0.8276** |
| 5 | 0.7530 | 0.8018 | 0.6431 | 0.5819 | 0.6539 | 0.5168 | **0.8267** |

**factor_approach_speed_facing_distdrop_axisalign (rank 1, AUC 0.8389)**

**Description.** Approach factor: facing the companion with high approach speed, a decreasing within-window distance trend, combined with body-axis alignment information.

- Designed target: **Approach** · Mode: batch · seq_length: 60 · n_samples: 3595
- Best class: Approach (AUC 0.8389, F1 0.0222)
- Valid classes: Approach (AUC 0.8389, F1 0.0222), Explore (AUC 0.6903, F1 0.4526)

**Code.**

```
_P0 = 1.0
_P1 = 1.0
_P2 = 1.0
_P3 = 1.0
_P4 = 1.0
i_app = idx.get('approach_speed', -1)
i_face = idx.get('self_facing_other', -1)
i_dist = idx.get('dist_to_other', -1)
i_speed = idx.get('speed', -1)
i_axis = idx.get('body_axis_align', -1)
if i_app < 0 or i_face < 0 or i_dist < 0 or i_speed < 0 or i_axis < 0:
    _result = np.full(windows.shape[0], np.nan)
else:
    app_mean = np.mean(windows[:, :, i_app], axis=1)
    app_stable = np.maximum(_P1 - np.std(windows[:, :, i_app], axis=1), 0)
    face_mean = np.mean(windows[:, :, i_face], axis=1)
    dist_drop = np.maximum(windows[:, 0, i_dist] - windows[:, -1, i_dist],
0)
    speed_mean = np.mean(windows[:, :, i_speed], axis=1)
```

```
    axis_mean = np.mean(windows[:, :, i_axis], axis=1)
    _result = _P0 * app_mean * face_mean * (_P2 * dist_drop + _P3 * speed_me
an) * (_P4 * axis_mean + app_stable)
```

#### factor_approach_facing_closure_motion_alignment_llmp (rank 2, AUC 0.8321)

**Description.** Approach factor: jointly characterizes facing the companion, approach speed, head-to-tail distance shrinkage, and the alignment between self-motion and body axis, emphasizing active approach from far to near.

- Designed target: **Approach** · Mode: batch · seq_length: 15 · n_samples: 18405
- Best class: Approach (AUC 0.8321, F1 0.0694)
- Valid classes: Approach (AUC 0.8321, F1 0.0694), Explore (AUC 0.7042, F1 0.5032)

**Code.**

```
_P0 = 1.0
_P1 = 1.0
_P2 = 1.0
_P3 = 0.1
_P4 = 1.0
_P5 = 1.0
_P6 = 1.0
_P7 = 1.0
_P8 = 1.0
i_face = idx.get('self_facing_other', -1)
i_app = idx.get('approach_speed', -1)
i_dist = idx.get('dist_to_other', -1)
i_sp = idx.get('speed', -1)
i_align = idx.get('body_axis_align', -1)
if min(i_face, i_app, i_dist, i_sp, i_align) < 0:
    _result = np.full(len(windows), np.nan)
else:
    face_m = np.mean(windows[:, :, i_face], axis=1)
    app_m = np.mean(windows[:, :, i_app], axis=1)
    sp_m = np.mean(windows[:, :, i_sp], axis=1)
    align_m = np.mean(windows[:, :, i_align], axis=1)
    closure = np.maximum(windows[:, 0, i_dist] - windows[:, -1, i_dist], np.
zeros(len(windows)))
    dist_var = np.std(windows[:, :, i_dist], axis=1)
    directed = _P0 * face_m + _P1 * app_m + _P2 * align_m
    drive = (_P3 * sp_m + _P4 * closure) ** _P5
    _result = directed * drive * (_P6 + _P7 * dist_var) ** _P8
```

#### factor_approach_mutual_axis_closure_speed_ramp (rank 3, AUC 0.8302)

**Description.** Approach factor: uses facing the companion, a shortening distance trend, strengthening approach speed, body-axis alignment and relative speed to characterize the temporal process of active movement toward the companion from afar.

- Designed target: **Approach** · Mode: batch · seq_length: 15 · n_samples: 18405
- Best class: Approach (AUC 0.8302, F1 0.0597)
- Valid classes: Approach (AUC 0.8302, F1 0.0597), Explore (AUC 0.7091, F1 0.4963)

**Code.**

```
_P0 = 1.0
_P1 = 1.0
_P2 = 1.0
_P3 = 1.0
_P4 = 1.0
_P5 = 1.0
i_face = idx.get('self_facing_other', -1)
i_dist = idx.get('dist_to_other', -1)
i_app = idx.get('approach_speed', -1)
i_axis = idx.get('body_axis_align', -1)
i_rel = idx.get('relative_speed', -1)
i_speed = idx.get('speed', -1)
if i_face < 0 or i_dist < 0 or i_app < 0 or i_axis < 0 or i_rel < 0 or i_spe
ed < 0:
    _result = np.full(len(windows), np.nan)
else:
    face_mean = np.mean(windows[:, :, i_face], axis=1)
    dist_closure = windows[:, 0, i_dist] - windows[:, -1, i_dist]
    app_ramp = windows[:, -1, i_app] - windows[:, 0, i_app]
    axis_mean = np.mean(windows[:, :, i_axis], axis=1)
    rel_mean = np.mean(windows[:, :, i_rel], axis=1)
    speed_mean = np.mean(windows[:, :, i_speed], axis=1)
    social_drive = _P0 * face_mean + _P1 * dist_closure + _P2 * app_ramp + _
P3 * axis_mean
    motion_drive = _P4 * rel_mean + _P5 * speed_mean
    _result = social_drive * motion_drive
```

#### factor_approach_facing_closure_speed_tailcalm_consistency (rank 4, AUC 0.8276)

**Description.** Approach factor: emphasizes facing the companion with simultaneously high approach speed and own speed, while requiring the tail motion to stay moderate and the approach process to persist throughout the window.

- Designed target: **Approach** · Mode: batch · seq_length: 60 · n_samples: 3595
- Best class: Approach (AUC 0.8276, F1 0.0243)
- Valid classes: Approach (AUC 0.8276, F1 0.0243), Explore (AUC 0.6921, F1 0.4495)

**Code.**

```
_P0 = 1.0
_P1 = 1.0
_P2 = 1.0
_P3 = 1.0
_P4 = 1.0
_P5 = 1.0
_P6 = 1.0
_P7 = 1.0
_P8 = 1.0
_P9 = 1.0
i_face = idx.get('self_facing_other', -1)
i_app = idx.get('approach_speed', -1)
i_speed = idx.get('speed', -1)
i_tailmotion = idx.get('tail_motion', -1)
i_dist = idx.get('dist_to_other', -1)
if i_face < 0 or i_app < 0 or i_speed < 0 or i_tailmotion < 0 or i_dist < 0:
    _result = np.full(windows.shape[0], np.nan)
else:
    face_m = np.mean(windows[:, :, i_face], axis=1)
    app_m = np.mean(windows[:, :, i_app], axis=1)
    speed_m = np.mean(windows[:, :, i_speed], axis=1)
    tail_m = np.mean(windows[:, :, i_tailmotion], axis=1)
    app_std = np.std(windows[:, :, i_app], axis=1)
    dist_drop = windows[:, 0, i_dist] - windows[:, -1, i_dist]
    dist_drop_pos = np.maximum(dist_drop, np.zeros_like(dist_drop))
    social_drive = _P0 * np.power(face_m, _P1) * (_P2 * app_m + _P3 * dist_d
rop_pos)
    locomotion = _P4 * speed_m
    calm_tail = _P5 / (_P6 + tail_m)
    consistency = _P7 / (_P8 + app_std)
    _result = social_drive * locomotion * calm_tail * consistency + _P9 * di
st_drop_pos * face_m
```

#### factor_explore_object_speed_heading_jitter_social_far (rank 5, AUC 0.8267)

**Description.** Object-exploration factor: sustained locomotion with large direction changes while staying far from the companion with weak social facing, characterizing direction-unstable exploration of non-social targets.

- Designed target: **Explore** · Mode: batch · seq_length: 60 · n_samples: 3595

- Best class: Approach (AUC 0.8267, F1 0.0270)
- Valid classes: Approach (AUC 0.8267, F1 0.0270), Climb (AUC 0.8018, F1 0.0205), Explore (AUC 0.7530, F1 0.5237), Blank (AUC 0.6539, F1 0.5258)

**Code.**

```
_P0 = 1.0
_P1 = 1.0
_P2 = 1.0
_P3 = 1.0
_P4 = 1.0
_P5 = 1.0
_P6 = 1.0
_P7 = 0.5
i_speed = idx.get('speed', -1)
i_heading = idx.get('heading_diff', -1)
i_dist = idx.get('dist_to_other', -1)
i_face = idx.get('self_facing_other', -1)
i_app = idx.get('approach_speed', -1)
i_acc = idx.get('acceleration', -1)
if i_speed < 0 or i_heading < 0 or i_dist < 0 or i_face < 0 or i_app < 0 or i_acc < 0:
    _result = np.full(len(windows), np.nan)
else:
    move_level = np.mean(windows[:, :, i_speed], axis=1) * (_P0 - np.std(windows[:, :, i_speed], axis=1))
    heading_jitter = _P1 * np.std(windows[:, :, i_heading], axis=1) + _P2 * (np.max(windows[:, :, i_heading], axis=1) - np.min(windows[:, :, i_heading], axis=1))
    far_social = _P3 * np.mean(windows[:, :, i_dist], axis=1) * (_P4 - np.mean(windows[:, :, i_face], axis=1))
    nondirected = (_P5 - np.abs(windows[:, -1, i_app] - windows[:, 0, i_app])) * (_P6 - np.mean(windows[:, :, i_app], axis=1))
    accel_texture = np.std(windows[:, :, i_acc], axis=1)
    base = move_level * heading_jitter * far_social + nondirected + accel_texture
    _result = np.where(move_level > _P7, base, base * move_level)
```
